# *Plasmodium falciparum* contains functional SCF and CRL4 ubiquitin E3 ligases, and CRL4 is critical for cell division and membrane integrity

**DOI:** 10.1101/2023.04.18.537323

**Authors:** Zeba Rizvi, G. Srinivas Reddy, Somesh M. Gorde, Priyanka Pundir, Divya Das, Puran Singh Sijwali

## Abstract

Protein ubiquitination is essential for cellular homeostasis and regulation of several processes, including cell division and genome integrity. Ubiquitin E3 ligases determine substrate specificity for ubiquitination, and Cullin-RING ubiquitin E3 Ligases (CRLs) make the largest group among the ubiquitin E3 ligases. Although conserved and most studied in model eukaryotes, CRLs remain underappreciated in *Plasmodium* and related parasites. To investigate the CRLs of human malaria parasite *Plasmodium falciparum*, we generated parasites expressing tagged *P. falciparum* cullin-1 (PfCullin-1), cullin-2 (PfCullin-2), Rbx1 (PfRbx1) and Skp1 (PfSkp1). PfCullin-1 and PfCullin-2 were predominantly expressed in erythrocytic trophozoite and schizont stages, with nucleocytoplasmic localization and chromatin association, suggesting their roles in different cellular compartments and DNA-associated processes. Immunoprecipitation, *in vitro* protein-protein interaction and ubiquitination assay confirmed the presence of a functional SCF (PfSCF), comprising of PfCullin-1, PfRbx1, PfSkp1, PfFBXO1 and calcyclin binding protein. Immunoprecipitation, sequence analysis and ubiquitination assay indicated that PfCullin-2 forms a functional human CRL4-like complex (PfCRL4), consisting of PfRbx1, cleavage and polyadenylation specific factor subunit_A and WD40 repeat proteins. PfCullin-2 knock-down at the protein level, which would hinder PfCRL4 assembly, significantly decreased asexual and sexual erythrocytic stage development. Several pathways, including protein translation and folding, lipid biosynthesis and transport, DNA replication, and protein degradation were dysregulated upon PfCullin-2-depletion, which likely reflects association of PfCRL4 with multiple pathways. Consistent with dysregulation of multiple pathways, PfCullin-2-depleted schizonts had poorly delimited merozoites and internal membraned structures, suggesting a role of PfCRL4 in maintaining membrane integrity. PfCullin-2-depleted parasites had significantly lower number of nuclei/parasite than the normal parasites, indicating a crucial role of PfCRL4 in cell division. Taken together, we for the first time demonstrate the presence of functional CRLs in *P. falciparum*, with crucial roles for PfCRL4 in cell division and maintaining membrane integrity. This study will benefit investigation of similar ligases in related parasites.

## Introduction

*Plasmodium* species cause malaria in humans and several other vertebrates. Human malaria is a huge burden, with the latest estimates of about 247 million cases and 6,19,000 deaths in 2021 (The World malaria report 2022, WHO)*. Plasmodium* species undergo multi-stage development in diverse cellular environments, with massive changes in the stage-specific proteomes and organelles, which must require protein turnover pathways like the ubiquitin proteasome system (UPS) and autophagy. UPS has key roles in diverse cellular processes, including cell cycle progression, differentiation, endocytic transport, transcription, DNA repair, signalling and antigen presentation [1–7]. The *Plasmodium* UPS is a validated drug target, and its inhibition has synergistic effect on artemisinin potency on artemisinin susceptible and resistant *P. falciparum* strains, which further strengthens the therapeutic potential and warrants investigation of *Plasmodium* UPS [8–15],[11, 16–19].

The core UPS components are ubiquitin, ubiquitination enzymes and proteasome. Ubiquitination, the covalent conjugation of ubiquitin to the ε-amino group of a lysine residue of the substrate protein, involves sequential action of ubiquitin-activating enzyme E1, transfer of the activated ubiquitin to ubiquitin-conjugating enzyme E2, followed by transfer of ubiquitin from the E2-ubiquitin conjugate to the substrate lysine residue by ubiquitin E3 ligase. Finally, the ubiquitinated proteins are degraded by the proteasome. However, ubiquitination of substrate also acts as a non-degradation signal, which depends on the ubiquitination chain topology [20].

Ubiquitin E3 ligases recognize substrate proteins, thereby, dictate target specificity of the UPS. There are 600-700 putative ubiquitin E3 ligases in humans, which are classified into RING E3s, HECT E3s and RBR E3s depending on the domain organization and the mechanism of ubiquitin transfer to the substrate [21, 22]. RING E3s contain structurally similar RING (Really Interesting New Gene) or U-box domain proteins, which bind the E2-ubiquitin conjugate and facilitate ubiquitin transfer to the substrate. HECT E3s contain a characteristic HECT domain (homologous to the E6AP carboxyl terminus) and catalyse ubiquitin transfer to their own catalytic cysteine residue and then to the substrate. RBR (RING-in-between-RING) E3s contain a canonical RING finger domain (RING1), an In-Between-RING domain (IBR) and a RING2 domain. The RING1 domain accepts the E2-ubiquitin conjugate and mediates transfer of ubiquitin to the catalytic cysteine residue in RING2, which then transfers ubiquitin to the substrate. Some of the best studied RING E3s are cullin-RING E3 ligases (CRLs), which make about 40% of the total ubiquitin E3 ligases. CRLs contain a cullin subunit that serves as a scaffold for assembly of the complex, a RING-box protein that binds E2 of the E2-ubiquitin conjugate, and a substrate adaptor for recruitment of the substrate. Skp1-Cullin-1-F-box (SCF) and anaphase promoting complex/cyclosome (APC/C) are the two most notable CRLs, which mediate timely degradation of cell cycle regulatory proteins [23, 24]. Cullin proteins are evolutionary conserved, with 8 in humans (Cul 1-3, 4A, 4B, 5, 7 and 9/PARC) and 3 in *S. cerevisiae* (cul1, cul3, cul8) [25–27]. CRLs have been mostly named based on the cullin subunit present in the complex; for example, cullin-1 makes CRL1 that is also known as SCF, and Cullin-4A and Cullin-4B make CRL4A and CRL4B, respectively.

CRLs have been shown to have key roles in cell cycle regulation [28], DNA replication [29], and DNA damage repair [30]. A key regulatory role of SCF and APC/C is timely degradation of cyclins and cyclin-dependent kinase inhibitors (CKIs) during eukaryotic cell cycle [28], which ensures chromosome replication, uniform segregation of chromosomes to opposite poles, followed by cytokinesis into two daughter cells. However, interestingly, *Plasmodium* cell division involves multiple rounds of nuclear divisions prior to cytokinesis, followed by sequestration of each nucleus with organelles and other cellular contents at the parasite plasma membrane, resulting into multiple daughter cells. This likely requires unique mode of regulation, including a role of the UPS.

More than half of the *P. falciparum* proteome represents possible targets of ubiquitination, suggesting both regulatory and homeostatic roles of the UPS [31]. The studies on the role of UPS in *Plasmodium* biology are limited to few UPS components. The *P. falciparum* ubiquitin-activating enzyme 1 (PfUBA1) and ubiquitin C-terminal hydrolase 37 (PfUCH37) have been shown to be essential for erythrocytic stage development [32, 33]. The HECT E3 ligase has been shown to have roles in cell cycle progression, invasion of erythrocytes by merozoites, and virulence [34, 35]. The knock-out of ubiquitin-conjugating enzyme 13 (UBC13) decreased parasite growth and increased susceptibility to DNA mutagens and dihydroartemisinin [36]. The *P. falciparum* ubiquitin C-terminal hydrolase 3 (PfUCH3) has been shown to have dual-hydrolase activity for ubiquitin and neuronal precursor cell-expressed developmentally down-regulated protein 8 (NEDD8) conjugates [37, 38]. An ERAD-like complex, including the ubiquitination enzymes, has been shown to be localized to the apicoplast, a plastid remnant, and is required for protein transport to the organelle [39, 40]. Although *Plasmodium* does not appear to have APC/C, the homologs of human APC/C subunits CDC20/CDH1 and APC3 have been shown to be vital for male gametogenesis [41, 42]. In a recent study, Rashpa, *et. al*. proposed an SCF-like complex in the mouse malaria parasite *P. berghei*, and showed that FBXO1 is required for cell division, gamete egress and integrity of the innermembrane complex in *P. berghei* [43].

We have previously shown that the *P. falciparum* cullin homologs are conjugated to NEDD8, which marks the active state of a CRL, and thus suggested active CRLs in *P. falciparum* [44]. However, it remains to be investigated if *Plasmodium* cullins form functional complexes, and what are the core components and roles of *Plasmodium* CRLs during parasite development. Given the regulatory roles and possibility of exploitation of ongoing CRL-targeted drug discovery efforts, we investigated *Plasmodium* cullins. We report the core components of the human malaria parasite *P. falciparum* Cullin-1 and Cullin-2 complexes, demonstrate that these two form functional SCF-like (PfSCF) and CRL-4-like (PfCRL4) ubiquitin E3 ligases, respectively. We further show that PfCRL4 is critical for cell division and membrane integrity.

## RESULTS

### *Plasmodium* genomes code for putative cullin, Rbx1 and Skp1 homologs

The cullin family proteins are conserved in eukaryotes and function as scaffold proteins in Cullin RING ubiquitin E3 ligases (CRLs), which form the largest group of ubiquitin E3 ligases. Although conserved across eukaryotes, the number of cullin genes is variable, with eight in *H. sapiens* and three in *S. cerevisiae* [4]. All *Plasmodium* species contain two putative cullin homologs, and we have previously reported that *Plasmodium falciparum* cullins are substrates of neddylation [44, 45]. The *P. falciparum* Cullin-1 (PfCullin-1, PF3D7_0811000) and Cullin-2 (PfCullin-2, PF3D7_0629800) share 13.9%-24.9% sequence identity with *H. sapiens* and *S. cerevisiae* cullins. Among the *H. sapiens* and *S. cerevisiae* cullins, PfCullin-1 is marginally closer to human Cullin-1 (HsCullin-1)/*S. cerevisiae* Cullin-1 (ScCullin-1) and PfCullin-2 is marginally closer to human Cullin-4B (HsCullin-4B)/*S. cerevisiae* Cullin-8 (ScCullin-8).

We also searched for PfCullin-1 and PfCullin-2 homologs in related Apicomplexan parasites, and compared Apicomplexan cullins with human cullins. All the Apicomplexans sequenced to date have 2-3 cullins, which share 13.3%-66.5% sequence identity with PfCullins, and also share the typical cullin domain architecture, consisting of the signature cullin homology domain and neddylation motif (Figure S1A, Table S1). The Apicomplexans cullins share maximum identity with HsCullin-1, HsCullin-3, and HsCullin-4A or HsCullin-4B (Table S1). Phylogenetic analysis showed clustering of Apicomplexan cullins into three separate groups: PfCullin-1 group, PfCullin-2 group, and the 3^rd^ group contained the 3^rd^ cullin homolog present in the Sarcocystidae family. *C. parvum* cullin domain containing protein (CDCP) also clustered with the 3^rd^ group. This suggested that PfCullin-1 and PfCullin-2 homologs are conserved across Apicomplexa (Figure S1B). PfCullin-1 and PfCullin-2 occupied separate branches on the phylogenetic tree and do not share node with any of the cullins of model organisms, indicating that *Plasmodium* cullins are distantly related to those of the model organisms (Figure S1C).

Cullin-1 associates with several other proteins, including Rbx1 and Skp1 proteins, to form the SCF ubiquitin E3 ligase. BLAST search of the Apicomplexan parasite genomes using *H. sapiens* and *S. cerevisiae* Rbx1 and Skp1 amino acid sequences identified homologs in all the parasites (Table S1). *P. falciparum* Rbx1 (PfRbx1, PF3D7_0319100) shares 64.7% identity with *H. sapiens* Rbx1 (HsRbx1) and 56.6% with *S. cerevisiae* Rbx1 (ScRbx1). *P. falciparum* Skp1 (PfSkp1, PF3D7_1367000) shares 44.7% sequence identity with *H. sapiens* Skp1 (HsSkp1) and 51.9% sequence identity with *S. cerevisiae* Skp1 (ScSkp1). Apicomplexan Rbx1 proteins share 21.6%-98.4% sequence identity and contain the characteristic RING_Ubox superfamily (RING_Ubox SF) domain (Figure S2A). Apicomplexan Skp1 proteins share 44.1%-97.1% sequence identity and contain the characteristic Skp1 superfamily (SS) domain (Figure S2B). The presence of cullin, Rbx1 and Skp1 homologs in Apicomplexan parasites suggests for the presence of at least two CRL ubiquitin E3 ligases.

### PfCullin-1 and PfCullin-2 are expressed in asexual erythrocytic stages, localize to both cytoplasm and nucleus, and associated with chromatin

To study the expression and localization of PfCullins, we replaced wild type PfCullin-1 and PfCullin-2 coding regions with PfCullin-1/GFP and PfCullin-2/cDD_HA_ coding sequences in *P. falciparum* D10 strain, respectively. Cloned PfCullin-1/GFP knock-in (PfCul1GFPKI) and PfCullin-2/cDD_HA_ knock-in (PfCul2KD_D10_) parasites were evaluated for replacement of the target gene by PCR and expression of fusion protein by western blotting. PfCul1GFPKI and PfCul2KD_D10_ parasites showed replacement of the respective wild type coding region with PfCullin-1/GFP and PfCullin-2/cDD_HA_ coding sequences, respectively (Figure S3). PfCullin-1/GFP was predominantly expressed in late trophozoite and schizont stages (Figure 1A), which is similar to its reported stage-specific transcription profile [46, 47]. PfCullin-2/cDD_HA_ is a fusion of PfCullin-2 with HA-tagged mutant *E. coli* DHFR, and it was predominantly expressed in trophozoite and schizont stages compared to ring stage (Figure 2A), which is in agreement with its reported transcription profile [46, 47]. The western blot of PfCul1GFPKI parasites had smaller PfCullin-1/GFP species, and the full-length protein (∼128 kDa) was observed at higher exposure only (Figure 1A). The western blot of PfCul2KD_D10_ parasites also had truncated species in addition to the full-length PfCullin-2/cDD_HA_ (∼155.7 kDa) (Figure 2A). We did not see any signal in the western blot of wild type parasite lysates with the antibodies used for detection of PfCullin-1/GFP and PfCullin-2/cDD_HA_ (Figure S3C and F), indicating that the smaller size bands are the degraded products of PfCullin-1/GFP and PfCullin-2/cDD_HA_. Both PfCullin-1/GFP and PfCullin-2/cDD_HA_ were localized all over the parasite except the black pigment-containing region, which is hemozoin in the food vacuole, a lysosome-equivalent organelle of *Plasmodium* (Figure 1B, Figure 2B). Confocal fluorescence images showed colocalization of PfCullin-1/GFP and PfCullin-2/cDD_HA_ with the nuclear stain DAPI (Figure 1C, Figure 2C), indicating that both these proteins are present in the nucleus as well. As some nuclear CRLs are chromatin-associated and regulate DNA-associated processes [48, 49], we checked PfCullin-1/GFP and PfCullin-2/cDD_HA_ for chromatin association. Both the proteins were present in cytoplasmic and chromatin fractions along with the marker proteins (H2B for nuclear fraction and the α2 proteasome subunit for cytoplasmic fraction) (Figure 1D, Figure 2D), suggesting that these proteins have DNA-associated functions.

**Figure 1.**
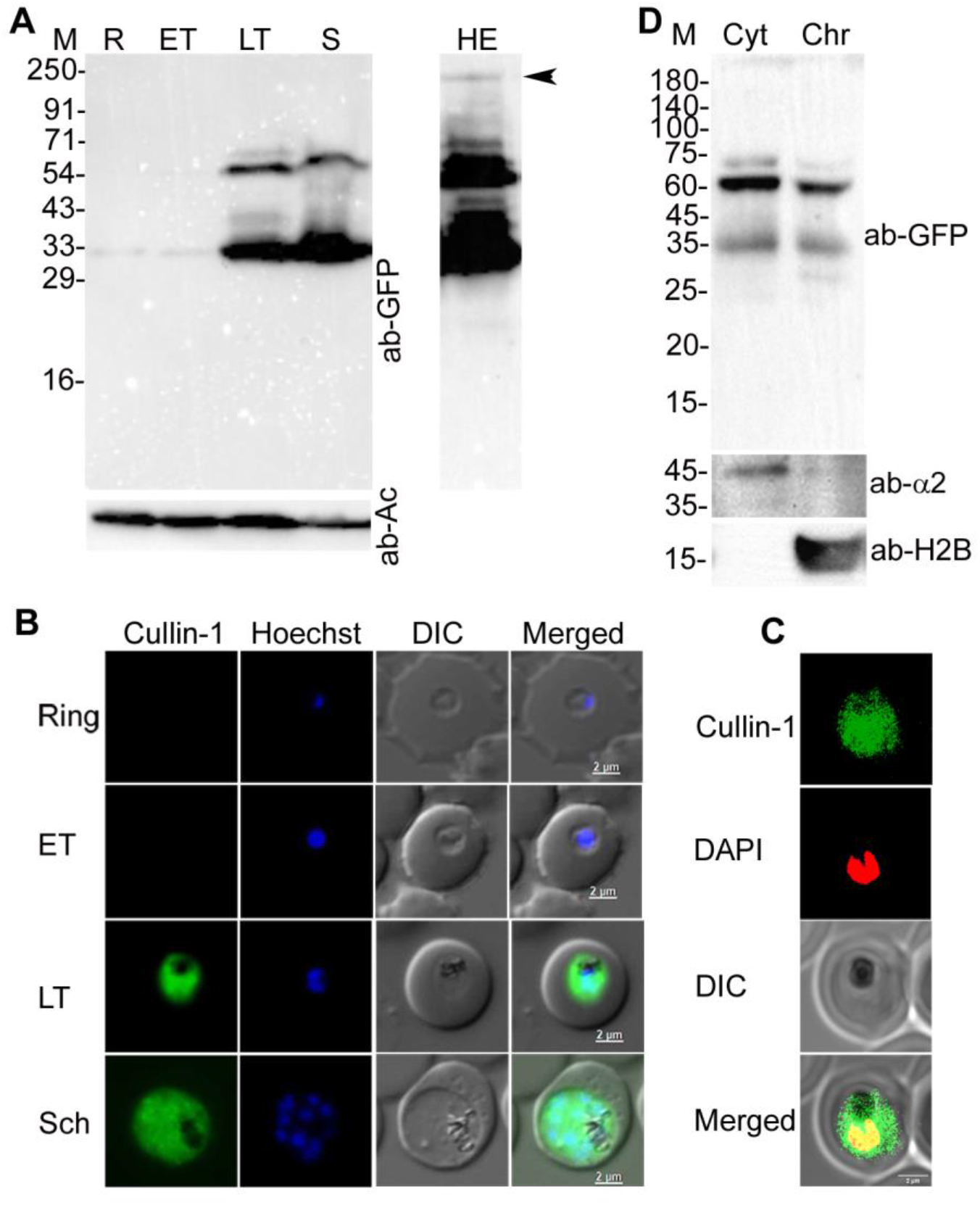
Expression and localization of PfCullin-1. **A.** The lysates of PfCul1GFPKI parasites corresponding to ring (R), early trophozoite (ET), late trophozoite (LT) and schizont (S) stages were assessed for expression of PfCullin-1/GFP and β-actin as a loading control by western blot using anti-GFP (ab-GFP) and mouse anti-β-actin-HRP (ab-Ac) antibodies, respectively. The full-length PfCullin-1/GFP protein (≈128 kDa) is indicated by arrow head in the higher exposure panel (HE). The lower molecular species could be truncated forms of the full-length PfCullin-1/GFP protein. **B.** The ring, early trophozoite (ET), late trophozoite (LT) and schizont (Sch) stages of PfCul1GFPKI parasites were assessed for localization of PfCullin-1/GFP by live-cell fluorescence microscopy. The panels show signal for PfCullin-1/GFP (Cullin-1), nuclear stain (Hoechst), bright field with RBC and parasite boundaries (DIC) and overlap of the three images (Merged). **C.** The confocal images of fixed PfCul1GFPKI trophozoite stage show colocalization of PfCullin-1/GFP with the nuclear stain DAPI (Pearson’s coefficient 0.59±0.10 from 27 images). The panels are as in B except DAPI was used for nuclear staining. **D.** The cytoplasmic (Cyt) and chromatin (Chr) fractions of PfCul1GFPKI trophozoites were assessed for the presence of PfCullin-1/GFP (ab-GFP), cytoplasmic protein α2 proteasome subunit (ab-α2), and nuclear protein histone-2B (ab-H2B) by western blotting. The sizes of protein markers (M) are in kDa and the size scale bars are in the merged panel.

**Figure 2.**
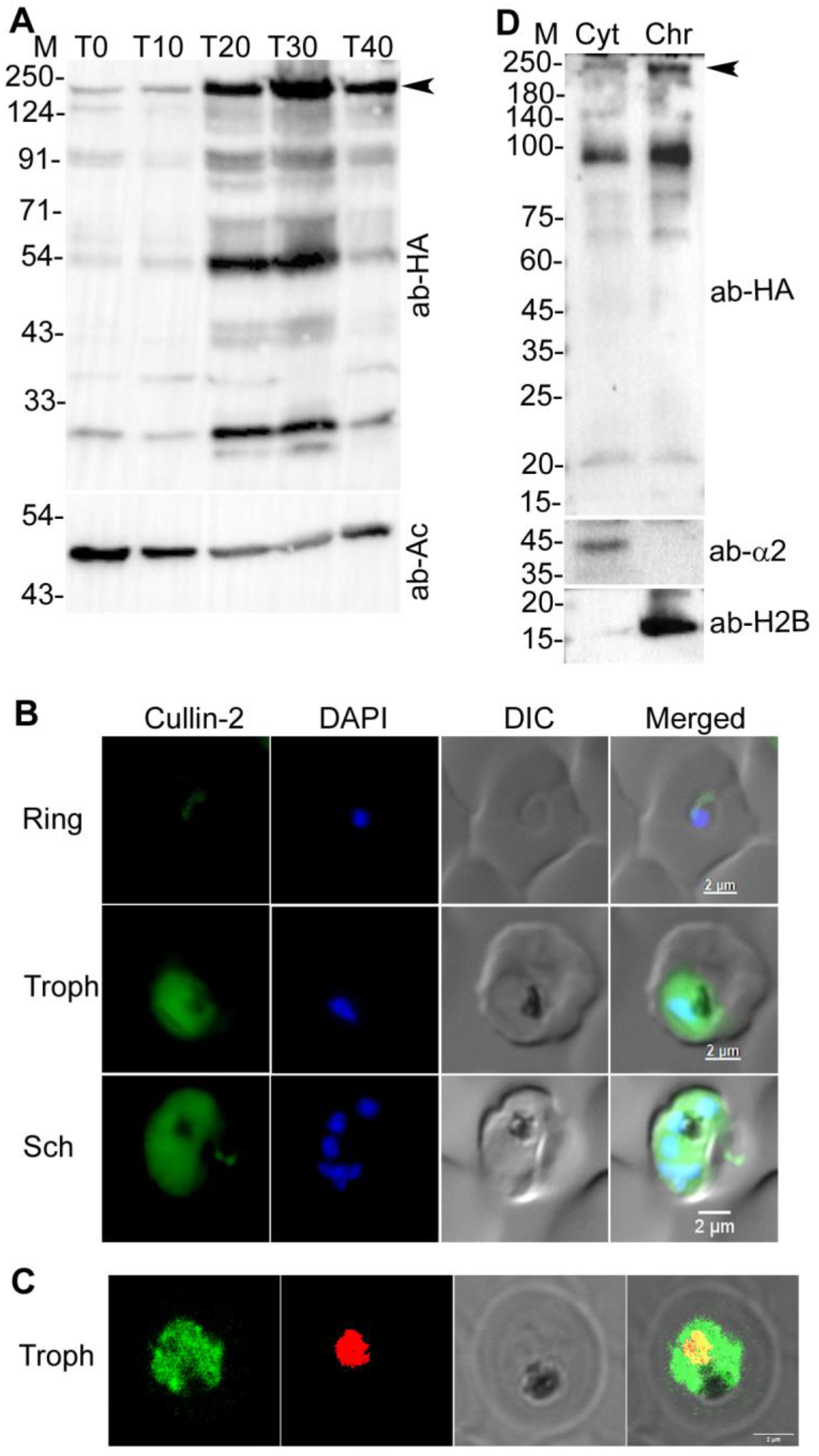
Expression and localization of PfCullin-2. **A.** Lysates of PfCul2KD_D10_ parasites corresponding to early ring (T0), late ring (T10), early trophozoite (T20), late trophozoite (T30) and schizont (T40) stages were assessed for expression of PfCullin-2/cDD_HA_ and β-actin as a loading control by western blot using anti-HA (ab-HA) and anti-β-actin (ab-Ac) antibodies, respectively. The full-length PfCullin-2/cDD_HA_ protein (≈155.74 kDa) is indicated with arrow, and the lower molecular bands could be truncated forms of the full-length protein. **B.** The ring, trophozoite (Troph), and schizont (Sch) stages of PfCul2KDD10 parasites were assessed for localization of PfCullin-2/cDD_HA_ by IFA using anti-HA antibodies. The panels show signal for PfCullin-2/cDD_HA_ (Cullin-2), nuclear stain (DAPI), bright field with RBC and parasite boundaries (DIC) and overlap of the three images (Merged). **C.** The confocal images of PfCul2KDD10 trophozoite (Troph) show colocalization of PfCullin-2/cDD_HA_ with DAPI (Pearson’s coefficient 0.59±0.11 from 13 images). **D.** The cytoplasmic (Cyt) and chromatin (Chr) fractions of PfCul2KDD10 trophozoites were assessed for the presence of PfCullin-2/cDD_HA_ (ab-HA), cytoplasmic protein α2 proteasome subunit (ab-α2), and chromatin protein histone-2B (ab-H2B) by western blotting. The sizes of protein markers (M) are in kDa and the size scale bars are in the merged panel.

### PfRbx1 and PfSkp1 localize to both cytoplasm and nucleus, and are associated with chromatin

The coding regions of PfRbx1 with C-terminal Myc-tag (PfRbx1myc) and PfSkp1 with C-terminal GFP-tag (PfSkp1GFP) were episomally expressed in *P. falciparum* D10 strain, and the recombinant parasites (PfRbx1myc-epi and PfSkp1GFP-epi) were assessed for localization of tagged-proteins. Western blots of PfRbx1myc-epi and PfSkp1GFP-epi parasites expressed the respective fusion proteins (Figure 3A, D), which were localized throughout the parasite except the food vacuole and colocalized with the nuclear stain DAPI (Pearson’s coefficient of 0.73±0.11 for PfRbx1myc and 0.55±0.12 for PfSkp1GFP) (Figure 3B, E), indicating that PfRbx1 and PfSkp1 are localized to the cytoplasm and nucleus. PfRbx1myc and PfSkp1GFP proteins were also observed in the chromatin fraction along with the control proteins (H2B for nuclear fraction and the α2 proteasome subunit for cytoplasmic fraction) (Figure 3C, F). The nucleocytoplasmic localization and chromatin association of PfCullin-1, PfCullin-2, PfRbx1 and PSkp1 suggest their roles in both the compartments and DNA-associated processes.

**Figure 3.**
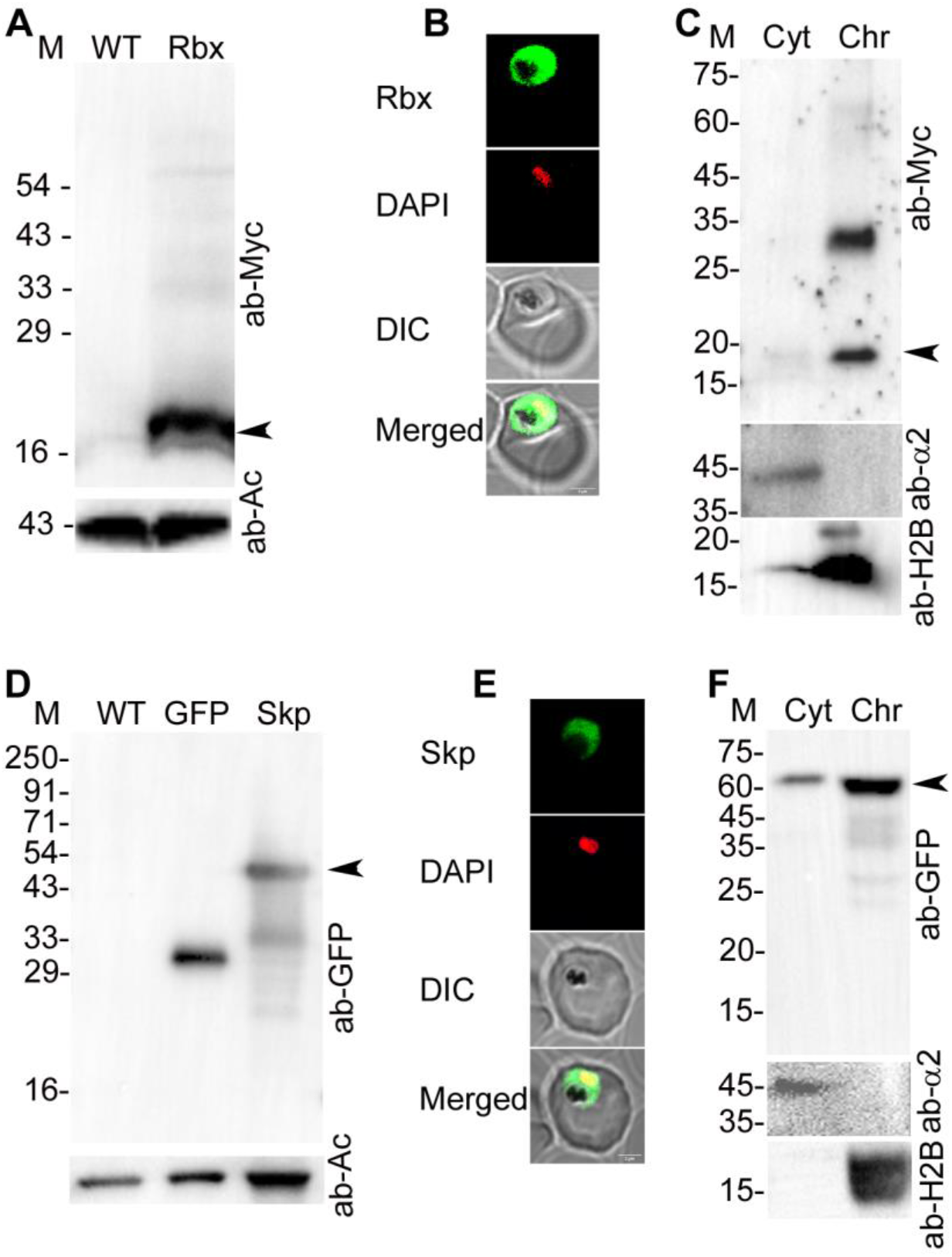
Localization of PfRbx1 and PfSkp1. **A.** The western blot of wild type (WT) and PfRbx1myc-epi (Rbx) parasite lysates shows expression of PfRbx1myc (ab-Myc) and β-actin as a loading control (ab-Ac). The arrow indicates PfRbx1myc protein (≈15 kDa). **B.** The confocal images of PfRbx1myc-epi trophozoite show colocalization of PfRbx1myc with the nuclear stain DAPI (Pearson’s coefficient 0.73±0.11 from 20 images). The panels show signal for PfRbx1myc (Rbx), DAPI, bright field with RBC and parasite boundaries (DIC) and overlap of the three images (Merged). **C.** The cytoplasmic (Cyt) and chromatin (Chr) fractions of PfRbx1myc-epi trophozoites were assessed for the presence of PfRbx1myc (ab-Myc), cytoplasmic protein α2 proteasome subunit (ab-α2) and nuclear protein histone-2B (ab-H2B) by western blotting. The arrow indicates PfRbx1myc protein (≈15 kDa). **D.** The western blot of wild type (WT), GFP control (GFP) and PfSkp1GFP-epi (Skp) parasite lysates shows expression of PfSkp1GFP or GFP (ab-GFP) and β-actin as a loading control (ab-Ac). The arrow indicates PfSkp1GFP protein (≈48 kDa). **E.** The confocal images of fixed PfSkp1GFP-epi trophozoite show colocalization of PfSkp1GFP with the nuclear stain DAPI. The panels show signal for PfSkp1GFP (Skp), DAPI, bright field with RBC and parasite boundaries (DIC), and overlap of the three images (Merged). **F.** The western blot of cytoplasmic (Cyt) and chromatin (Chr) fractions of PfSkp1GFP-epi trophozoites indicates the presence of PfSkp1GFP (ab-GFP), cytoplasmic protein α2 proteasome subunit (ab-α2), and nuclear protein histone-2B (ab-H2B). The arrow indicates PfSkp1GFP protein (≈48 kDa). The protein size markers (M) are in kDa, and the size scale bars are shown in the merged image panels.

### PfCullin-1 forms an SCF-like complex

Since cullin-1, Rbx1 and Skp1 together with other proteins form SCF ubiquitin E3 ligase in several model organisms, we asked if *Plasmodium* has an SCF-like complex. We immunoprecipitated PfCullin-1/GFP, PfRbx1myc, PfSkp1GFP from PfCul1GFPKI, PfRbx1myc-epi and PfSkp1GFP-epi parasites, respectively, using Myc-trap (for PfRbx1myc-epi) or GFP-trap (for PfCul1GFPKI and PfSkp1GFP-epi) antibodies (Figure S4). As a control, wild type *P. falciparum* D10 and GFP-expressing parasites (GFP control) were also processed for immunoprecipitation using Myc-trap and GFP-trap antibodies, respectively (Figure S4). The immunoprecipitates (IP) were subjected to LC-MS/MS and the proteins present were analysed.

Twenty-five proteins were reproducibly present in PfCullin-1/GFP immunoprecipitates from two of the three biological repeats (Table S2), including PfCullin-1 and PfRbx1. PfNEDD8 was present in one of the repeats; conjugation of NEDD8 to the cullin subunit of CRLs promotes ubiquitination activity, including the SCF. Rbx1 binds to the C-terminal of cullin, and recruits the E2-ubiquitin conjugate. The presence of PfNEDD8 in PfCullin-1/GFP immunoprecipitates corroborates our previous report of the presence of PfCullin-1 in PfNEDD8 immunoprecipitates, and substantiates neddylation of PfCullin-1 [6]. DDI1 and Histone H2B variant, which are relevant to DNA-associated processes, were also present in the PfCullin-1/GFP immunoprecipitates.

Forty-two proteins were present in PfRbx1myc immunoprecipitates from two of the three biological repeats (Table S3). Some of the top hits included PfRbx1, PfCullin-1 and PfNEDD8, which are well known components of an SCF. Tubulin gamma chain was also a hit, which is required for microtubule nucleation and is a component of microtubule organizing centre (MTOC) and substrate of human CRL4 E3 ligases [7]. PfAtg18, cell cycle proteins (nucleosome assembly proteins, actin related protein, regulator of chromosome condensation), spliceosome components, chromatin re-modellers, and proteins involved in chromosome condensation were also present in the PfRbx1myc immunoprecipitates.

Eighteen proteins were reproducibly present in the PfSkp1GFP immunoprecipitates from two of the three biological repeats (Table S4), including PfSkp1, PfCullin-1, a putative FBXO1 protein, and calcyclin binding protein (CacyBP), which are also the known components of human SCF complex, suggesting that these proteins form an SCF complex in *P. falciparum*. Polyubiquitin (pUB), DDI1 that is involved in DNA-protein crosslink repair in *P. falciparum* [8], DNA repair and recombination protein RAD54, exported protein PHISTb that is involved in host cell remodelling, and anti-oxidant protein DJ1 were some other proteins present in the PfSkp1GFP immunoprecipitates.

The presence of PfCullin-1, PfNEDD8, PfRbx1, PfSkp1, PfFBXO1 and PfCacyBP in the immunoprecipitates suggested the presence of a typical SCF-like complex in *P. falciparum*. In human SCF complex, cullin-1 binds with Skp1 via its N-terminal and with Rbx1 via its C-terminal. Skp1 associates with FBXO1 through its C-terminal, which then binds to the substrate to be ubiquitinated. Rbx1 recruits an E2-ubiquitin conjugate, and NEDD8 conjugation to cullin-1 promotes transfer of ubiquitin to the substrate [9]. To check whether these proteins directly interact with each other, we produced GST-tagged recombinant N- and C-terminals of PfCullin-1 (GST/Cul1N, 1-300 aa; GST/Cul1C, 601-829 aa), PfFBXO1 (GST/FBXO), PfCacyBP (GST/CacyBP), PfRbx1 (GST/Rbx), and only GST using a bacterial expression system. We also produced 6×His/SUMO-tagged PfRbx1 (_His_SUMO/Rbx), 6×His/SUMO only (_His_SUMO), and 6×His-tagged PfSkp1 (_His_Skp) using a bacterial expression system (Figure S5). The recombinant proteins were assessed for interaction by dot blot protein overlay assay. _His_Skp interacted with GST/Cul1N, GST/FBXO and GST/CacyBP, but not with GST/Rbx and GST (Figure 4), indicating that the interaction is specific. _His_SUMO/Rbx interacted with GST/Cul1C, but not with GST/CacyBP, GST/FBXO and GST (Figure 4). _His_SUMO alone did not interact with GST/Cul1C and GST, indicating that the interaction of _His_SUMO/Rbx with GST/Cul1C is specific. Thus, our immunoprecipitation and protein overlay data for the first time demonstrate that *P. falciparum* contains an SCF-like complex of PfCullin-1, PfSkp1, PfRbx1, PfFBXO1 and PfCacyBP. Furthermore, the presence of PfNEDD8 in the immunoprecipitates of PfCullin-1/GFP and PfRbx1myc, and our previous report of neddylation of PfCullin-1 indicate the PfSCF undergoes neddylation.

**Figure 4.**
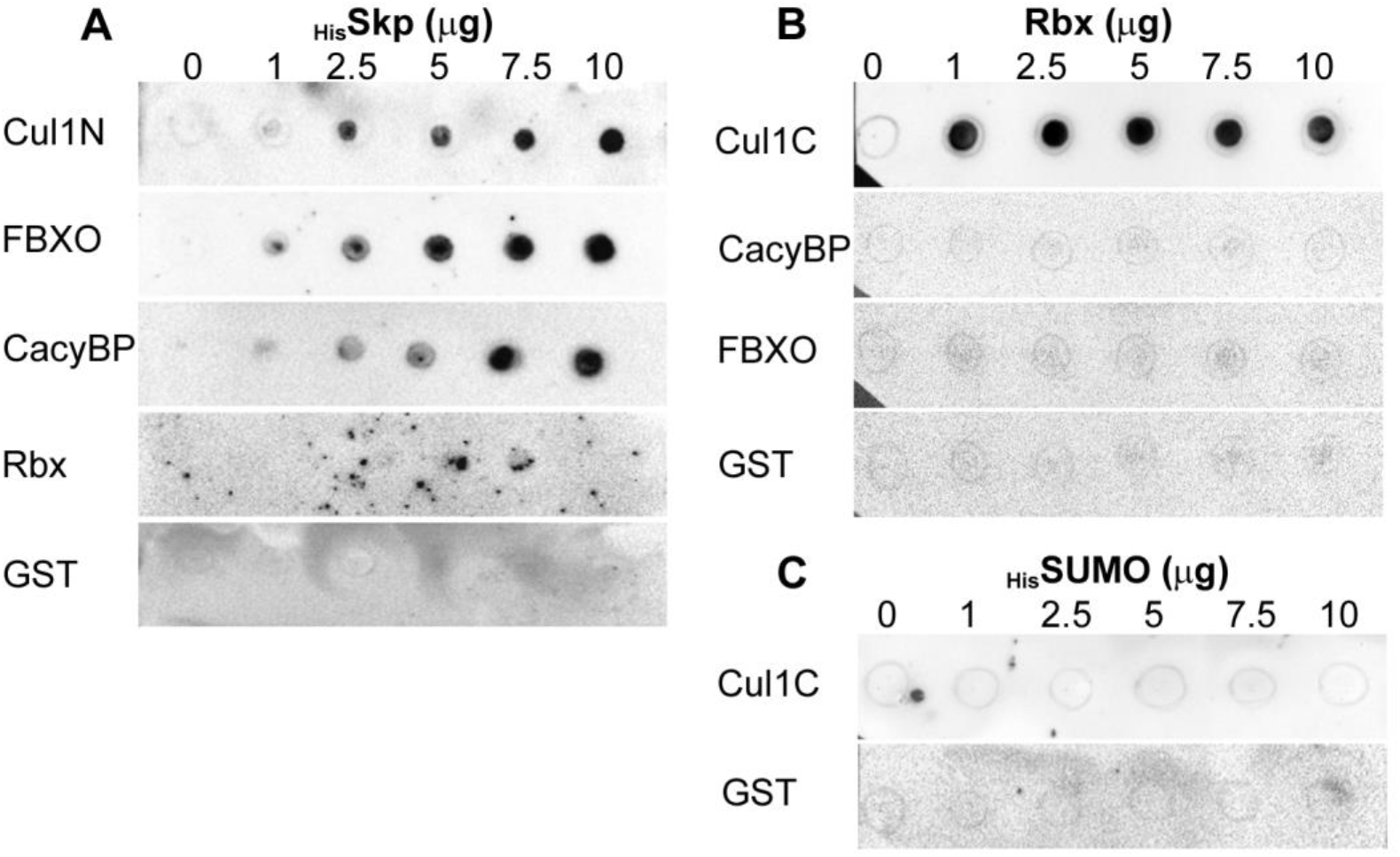
Interactions among PfSCF subunits. **A.** Blots containing spots of the indicated amounts of recombinant HisSkp were overlaid with recombinant GST/Cul1N (Cul1N), GST/FBXO1 (FBXO), GST/CacyBP (CacyBP), GST/Rbx (Rbx) and GST proteins. The blots were probed with anti-GST antibodies, which revealed interaction of HisSkp with Cul1N, FBXO, and CacyBP proteins. **B.** The blots containing spots of the indicated amounts of recombinant HisSUMO/Rbx (Rbx) were overlaid with recombinant GST/Cul1C (Cul1C), GST/CacyBP (CacyBP), GST/FBXO1 (FBXO), and GST proteins. The blots were probed with anti-GST antibodies, which revealed interaction of Rbx with Cul1C. **C.** The blots containing spots of the indicated amounts of recombinant 6×His/SUMO (HisSUMO) were overlaid with recombinant GST/Cul1C (Cul1C) and GST proteins, and probed with anti-GST antibodies. The blots indicated that HisSUMO and GST do not interact.

### PfCullin-2 appears to form a CRL4B-like complex

As cullin is an essential core component of CRLs, we investigated whether PfCullin-2 forms a CRL. We immunoprecipitated PfCullin-2/cDD_HA_ from PfCul2KD_D10_ parasites and subjected the immunoprecipitates to mass spectrometry (Figure S6). Forty-nine proteins were present in two of the three biological repeats (Table S5). Among the top 10 hits that are relevant to CRLs were PfCullin-2, cleavage and polyadenylation specific factor subunit A (PfCPSF_A), two WD40 repeat proteins (PfWD40A and PfWD40B), PfNEDD8, and PfRbx1 (Figure 5A). Sequence analysis of these hits indicated similarity in domain architecture with human Cullin-4B ubiquitin E3 ligase subunits, which includes Cullin-4B as a scaffold protein, DNA damage-specific binding protein 1 (DDB1) as an adaptor protein, Rbx1 as a ubiquitin-E2 recruiter, NEDD8 that promotes the ligase activity, and several WD40 repeat proteins as substrate receptors [50]. DDB1 binds to the N-terminal of Cullin-4B and recruits the substrate recognition subunit DDB1-CUL4-associated factor (DCAF), which is a WD40 repeat-containing FBXO protein. Rbx1 binds to the C-terminal of Cullin-4B.

**Figure 5.**
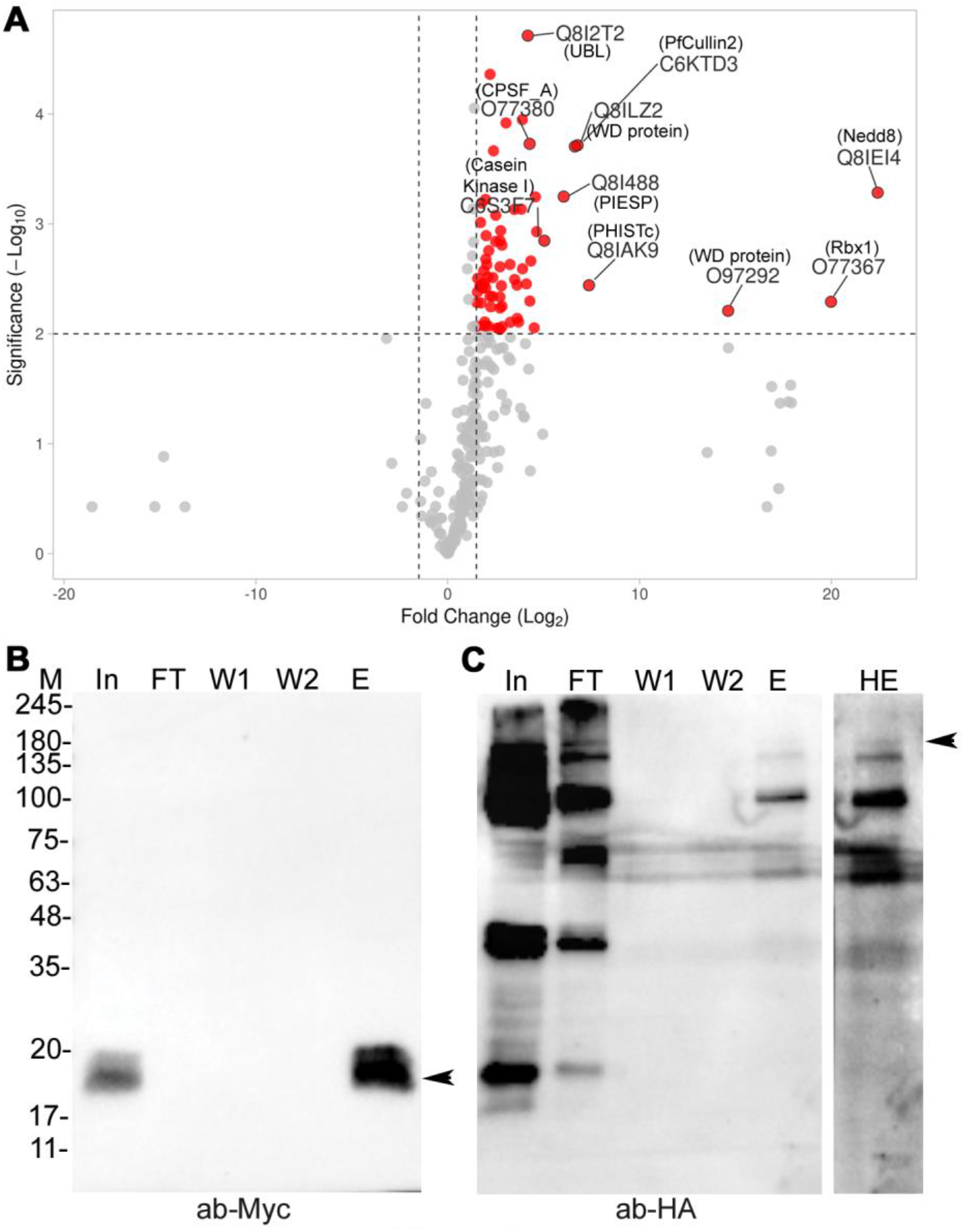
Analysis of the immunoprecipitated proteins and co-immunoprecipitation of PfCullin-2 with PfRbx1. **A.** The plot shows Log2-fold-enrichment (X-axis) of the proteins identified in PfCullin-2 immunoprecipitates over those in the control immunoprecipitates with significant -Log10 *P*-value (Y-axis). The top 10 hits are labelled. **B.** Equal amounts of PfRbx1myc-epi and PfCul2KDD10 parasite lysates were mixed and processed for immunoprecipitation using Myc-TRAP antibodies. Aliquots of the mixed lysate (In), flow through (FT), washes (W1, W2) and eluate (E) samples were evaluated for the presence of Rbx1myc by western blotting using anti-Myc antibodies (ab-Myc). The arrow indicates PfRbx1myc protein (≈16 kDa). **C.** The blot in B was stripped and probed to check for PfCullin-2/cDD_HA_ using anti-HA antibodies (ab-HA). The arrow indicates full-length PfCullin-2/cDD_HA_ protein (≈155 kDa) at higher exposure (HE). The sizes of protein markers (M) are in kDa.

PfCullin-2 shares maximum sequence identity with human Cullin-4B (HsCullin-4B) among all human cullins. Like HsCullin-4B, PfCullin-2 has a long N-terminus upstream of the putative cullin homology domain (703 aa in PfCullin-2 and 216 aa in HsCullin-4B) and a putative nuclear localization signal (Figure S7), suggesting that PfCullin-2 could be the *Plasmodium* counterpart of HsCullin-4B. A BLAST search of the Uniprot database with PfCPSF_A amino acid sequence identified DDB1 proteins of *A. thaliana* (2e-15), *Xenopus laevis* (5e-15), *Rattus norvegicus* (5e-14), *Homo sapiens* (2e-13) and *Mus musculus* (2e-13) as the top five hits. PfCPSF_A also shares domain architecture with DDB1 proteins, suggesting that it could be the *P. falciparum* DDB1 homolog (Figure S7). A BLAST search of the PlasmoDB with human DCAF1 did not show up any homolog. Human DCAF1, PfWD40A and PfWD40B share WD40 domain, and WD40 repeat proteins have been shown to function as substrate receptors for human CRL4 and some ubiquitin E3 ligases [50, 51]. It is likely that PfWD40A and PfWD40B serve as substrate receptors. Consistent with our previous report of the presence of PfCullin-2 in PfNEDD8 immunoprecipitates, the presence of PfNEDD8 in PfCullin-2/cDD_HA_ immunoprecipitates indicates neddylation of PfCullin-2 [44].

Attempts to express PfCPSF_A, PfWD40A, PfWD40B and PfCullin-2 in *E. coli* were unsuccessful, and we could not assess their direct interaction with each other *in vitro*. Nonetheless, successful co-immunoprecipitation of PfCullin-2/cDD_HA_ with PfRbx1myc from the mixed lysate of PfCul2KD_D10_ and PfRbx1myc-epi parasites strengthens PfRbx1-PfCullin-2 interaction (Figure 5B). The presence of PfCPSF_A, WD40 repeat proteins, PfNEDD8 and PfRbx1 as top hits in the PfCullin-2/cDD_HA_ immunoprecipitates, co-IP of PfCullin-2 with PfRbx1, similarity of PfCPSF_A with HsDDB1 and that of PfCullin-2 with HsCullin-4B together indicate that PfCullin-2 forms a human Cullin-4B-like ubiquitin ligase, which we have termed the *P. falciparum* CRL4 (PfCRL4).

PfCullin-2/cDD_HA_ immunoprecipitates also contained several proteins associated with diverse pathways, including phosphorylation (casein kinase 1), virulence (parasite-infected erythrocyte surface protein 2, KAHRP, translocon component PTEX88), exported proteins (exported protein 1 and PHISTc), transport to nucleus (exportin-7 and karyopherin alpha), RBC invasion by merozoites (rhoptry neck protein 3 and rhoptry neck protein 5). This implicates PfCRL4 in a variety of pathways. More work will be required to determine a direct link of PfCRL4 association with the proteins present in the immunoprecipitates.

### PfSCF and PfCRL4 are functional ubiquitin E3 ligases

We next asked whether PfSCF and PfCRL4 are functional ubiquitin E3 ligases. Although *in vitro* assembly of PfSCF and PfCRL4 would have been ideal to check their ubiquitination activity, attempts to obtain full-length recombinant PfCullin-1, PfCullin-2 and other subunits were unsuccessful due to their large sizes. We used bead-bound PfSkp1GFP immunoprecipitate, which contains most of the PfSCF components (PfCullin-1, PfSkp1, PfFBXO1 and PfCacyBP) except PfRbx1, along with recombinant GST/Rbx, ubiquitin-activating enzyme E1 and ubiquitin-conjugating enzyme E2 for ubiquitination assay in the presence or absence of ATP. The immunoprecipitate showed ATP-dependent ubiquitination activity (Figure 6A, B, C), indicating that PfSCF is a functional ubiquitin E3 ligase. Similarly, the bead-bound PfCullin-2/cDD_HA_ immunoprecipitate, which contains all the core components (PfCullin-2, PfCPSF_A, PfRbx1 and WD40 repeat proteins), showed ATP-dependent ubiquitination activity (Figure 6D, E, F), indicating that PfCRL4 is a functional ubiquitin E3 ligase. Interestingly, PfCullin-2 appeared to undergo auto-ubiquitination, which could be a regulatory mechanism of PfCRL4 function. Of note, a putative ubiquitin carboxy-terminal hydrolase was consistently co-purified with PfCullin-2/cDD_HA_, which may modulate the activity of PfCRL4.

**Figure 6.**
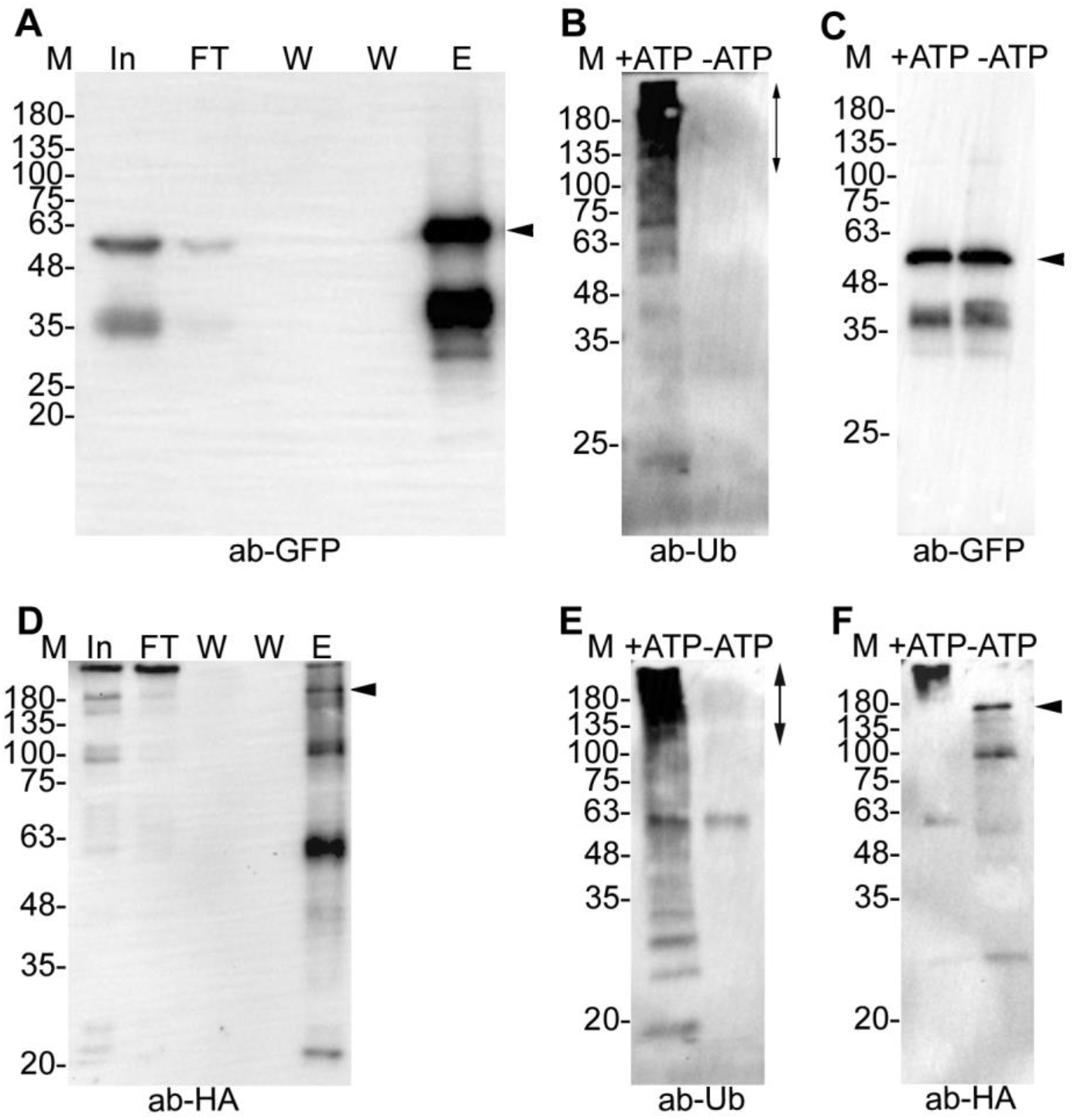
Ubiquitination activity of PfSCF and PfCRL4. PfSkp1GFP-epi and PfCul2KDD10 parasite lysates were processed for immunoprecipitation of PfSkp1GFP and PfCullin-2/cDD_HA_ using GFP-Trap and anti-HA antibodies, respectively. The bead-bound immunoprecipitates were washed, eluted or added to ubiquitination reactions with (+ATP) or without (-ATP) ATP as has been described in the methods section. **A.** Aliquots of the parasite lysate input (In), flow through (FT), washes (W), and eluate (E) were evaluated for Skp1GFP by western blotting using anti-GFP antibodies (ab-GFP). **B.** The western blot of PfSkp1GFP-containing ubiquitination reactions with (+ATP) or without (-ATP) ATP was probed with anti-ubiquitin antibodies (ab-Ub). It showed ATP-dependent ubiquitination activity. The vertical arrow indicates predominant ubiquitination. **C.** The blot in B was stripped and re-probed with anti-GFP antibodies (ab-GFP) to show that the reactions contained similar amount of PfSkp1GFP. **D.** As in A, the samples of PfCullin-2/cDD_HA_ immunoprecipitation were evaluated for PfCullin-2/cDD_HA_ by western blotting using anti-HA antibodies (ab-HA). **E.** The western blot of PfCullin-2/cDD_HA_-containing ubiquitination reactions with (+ATP) or without (-ATP) ATP was probed with anti-ubiquitin antibodies (ab-Ub). It showed ATP-dependent ubiquitination activity. The vertical arrow indicates predominant ubiquitination. **F.** The blot in E was stripped and probed with anti-HA antibodies (anti-HA). The full-length PfCullin-2/cDD_HA_ in +ATP lane migrated at higher size than that in –ATP lane and corresponds to the size of top most ubiquitination signal +ATP lane, suggesting auto-ubiquitination of PfCullin-2. The sizes of protein markers (M) are in kDa, and arrows indicate the respective proteins in the blots.

### PfCRL4 is crucial for asexual and sexual erythrocytic stage development

PfCullin-2 expression in asexual erythrocytic stages suggested a role for it during asexual stage parasite development. Hence, we generated a conditional PfCullin-2 knock-down line in *P. falciparum* 3D7 strain by replacing the wild type PfCullin-2 gene with PfCullin-2 coding sequence fused to HA-tagged mutant *E. coli* DHFR coding sequence (cDD_HA_). cDD or cDD-fusion protein is stable when bound to its ligand trimethoprim (TMP), but undergoes proteasomal degradation in the absence of TMP, allowing inducible knock-down at the protein level [52]. Accordingly, these parasites will have stable PfCullin-2/cDD_HA_ protein in the presence of trimethoprim (+TMP), which would be depleted in the absence of trimethoprim (-TMP). As cullin protein is the scaffold for CRL assembly, depletion of PfCullin-2 would prevent the formation of PfCRL4 complex and would indicate the effect of PfCRL4 depletion on parasite development.

PCR and western blot of a cloned recombinant line (PfCul2KD_3D7_) confirmed the replacement of wild type PfCullin-2 with PfCullin-2/cDD_HA_ coding sequence and expression of PfCullin-2/cDD_HA_ protein, respectively (Figure S8A, B). IFA showed PfCullin-2/cDD_HA_ expression in asexual stages of PfCul2KD_3D7_ parasites (Figure S8C). PfCul2KD_3D7_ parasites cultured in the absence of TMP showed decreased PfCullin-2/cDD_HA_ protein level and significantly lower parasitemia than those grown in the presence of TMP (Figure 7A-C). The knock-down effect was partial in the 1^st^ cycle and became prominent in 2^nd^ and 3^rd^ cycles. Hence, we primarily focussed on 2^nd^ and 3^rd^ cycle parasites for further evaluation of the effect of PfCullin-2 depletion on parasite development. Giemsa smears of PfCul2KD_3D7_ parasites cultured without TMP, but not with TMP, revealed clear morphological defects (distorted membrane, scattered hemozoin and shrunken nuclei) in a large number of trophozoites/schizonts (Figure 8A, B). Transmission electron micrographs of PfCul2KD_3D7_ parasites cultured with TMP showed clear demarcation of merozoites inside the schizont with distinct outer membrane and intracellular organelles, whereas the schizonts of parasites cultured without TMP showed loss of overall morphology with poor demarcation of the merozoite membrane and intracellular organelles (Figure 8C). Scanning electron microscopy images of PfCul2KD_3D7_ parasites cultured without TMP showed aberrant surface topology in 58% of the cells as compared to only 5.6% of the cells in parasites cultured with TMP (Figure S9). Together, these data indicate that PfCRL4 is critical for asexual erythrocytic development and membrane integrity.

**Figure 7.**
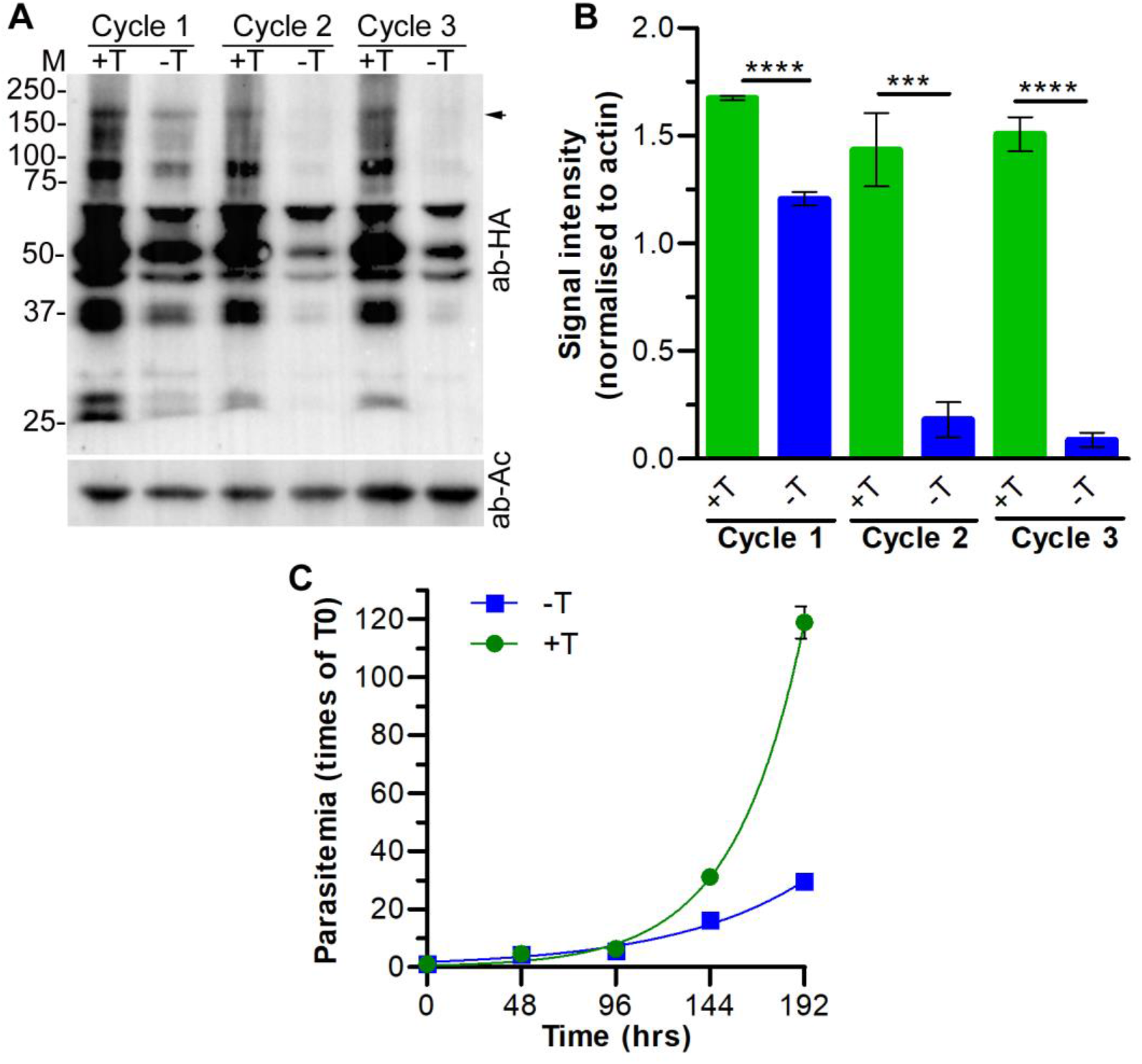
Effect of PfCullin-2 knock-down on asexual erythrocytic stage development. **A.** Synchronized PfCull2KD_3D7_ parasites were cultured with (+T) or without (-T) trimethoprim for three consecutive cycles, each spanning approximately 48 hrs. Equal aliquots of the cultures at the trophozoite stage were harvested in each cycle, and parasite lysates were assessed for PfCullin-2/cDD_HA_ protein level by western blotting using anti-HA (ab-HA) and anti-β-actin (ab-AC) antibodies. The arrow indicates full-length PfCullin-2/cDD_HA_ and the sizes of protein markers (M) are in kDa. **B.** The signal intensity of full-length PfCullin-2/cDD_HA_ and β-actin bands in A were measured, and the signal intensity of full-length PfCullin-2/cDD_HA_ was normalised with the signal intensity of corresponding β-actin band. The plot shows signal intensity of full-length PfCullin-2/cDD_HA_ band on Y-axis for the indicated parasite lysates on X-axis. The data is mean of three independent experiments with SD error bar. **C.** Synchronized PfCull2KD_3D7_ parasites were cultured with (+T) or without (-T) trimethoprim for four consecutive cycles. Parasitemia was determined at the beginning and end of each cycle, and shown as fold multiplication of the starting parasitemia over time. The data is mean of three independent experiments with SD error bar.

**Figure 8.**
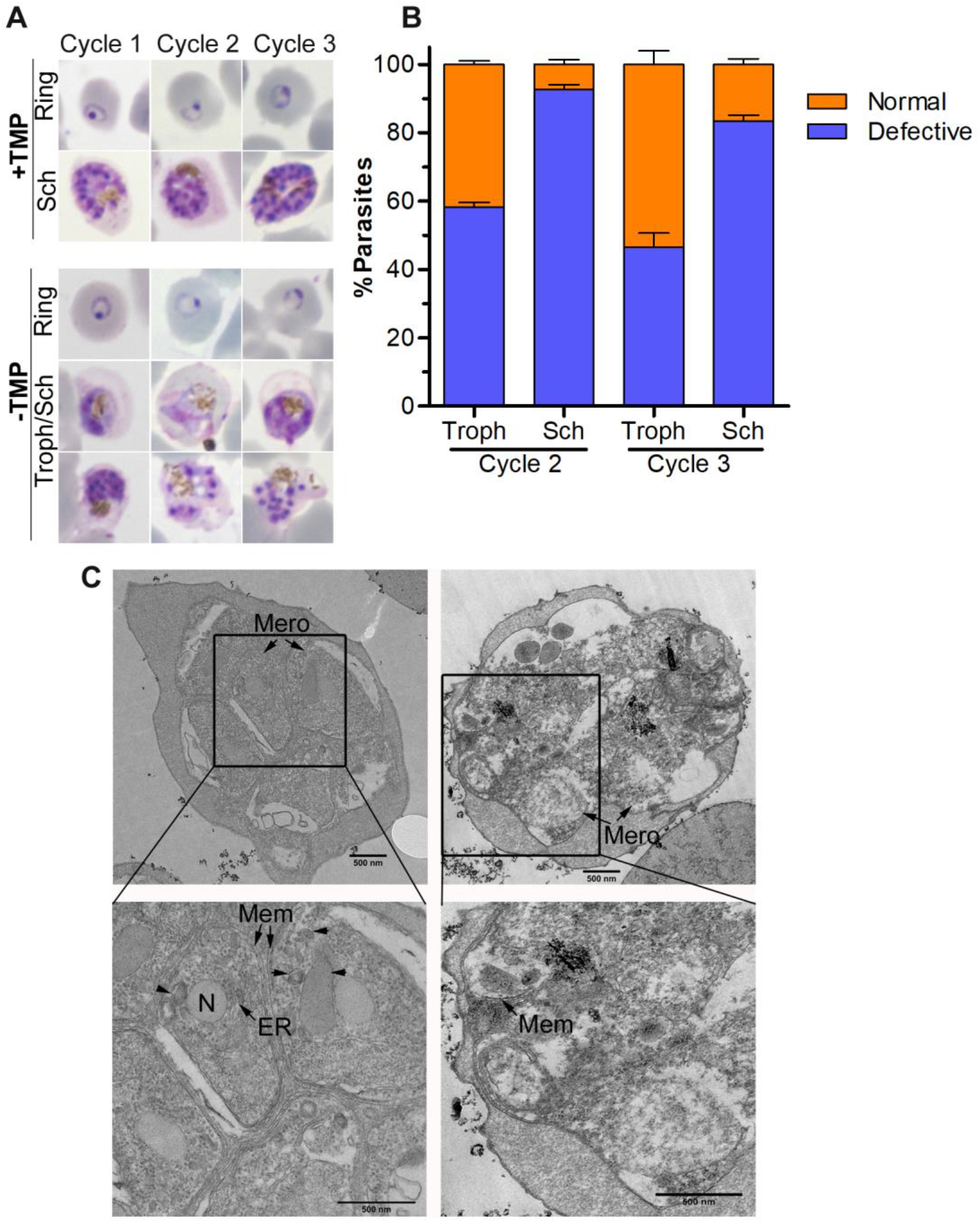
Synchronized PfCull2KD_3D7_ parasites were cultured with (+TMP) or without (-TMP) trimethoprim for three consecutive cycles, and parasites were observed for morphology at the end of each cycle. **A.** Shown are the Giemsa-stained smears of PfCull2KD_3D7_ parasite cultures from three cycles. **B.** The plot shows percentage of normal and morphologically defective (fragmented cell, indistinct nuclei and scattered hemozoin) trophozoites (Troph) and schizonts (Sch) in cycle 2 and cycle 3. The data is from two experiments, each based on at least 100 trophozoites/schizonts observed. **C.** Shown are the TEM images of 3^rd^ cycle schizonts with multiple merozoites (Mero). The zoomed-in section shows clearly delimited membraned structures (N: nucleus, ER: endoplasmic reticulum, Mem: membrane) in +TMP parasites, which are poorly visible in –TMP parasites.

The IFA of PfCul2KD_3D7_ gametocytes indicated prominent expression of PfCullin-2/cDD_HA_ in all gametocyte stages (Figure 9A), which prompted us to assess the effect of PfCullin-2 knock-down during sexual erythrocytic stage development. We induced gametocytogenesis and maintained gametocytes with or without TMP in the culture. The western blot of day 4 gametocytes without TMP showed decreased PfCullin-2 protein level as compared to those grown with TMP (Figure 9B), and there was a significant reduction in the number of gametocytes in –TMP culture (Figure 9C), indicating an important role of PfCRL4 during sexual erythrocytic stage development.

**Figure 9.**
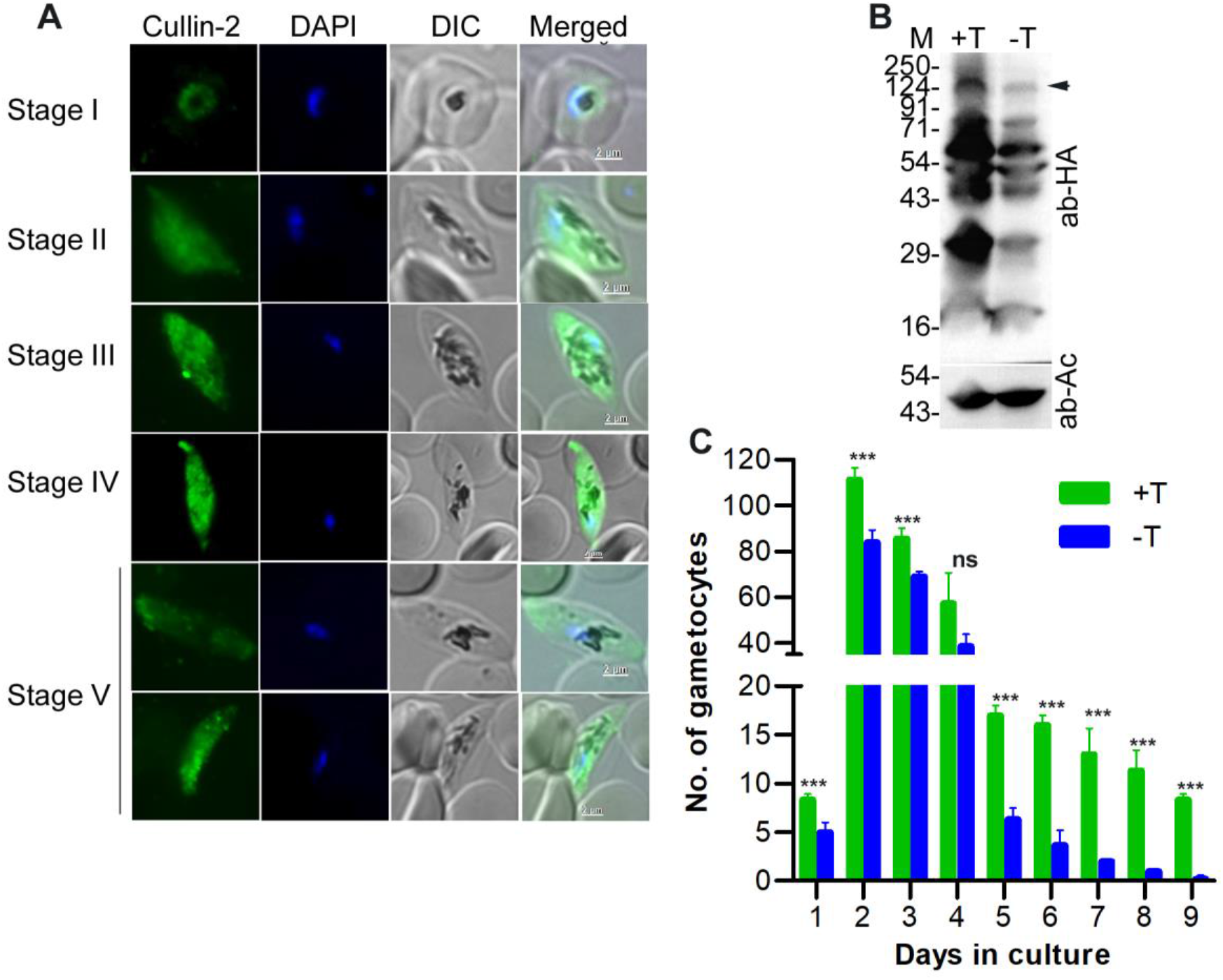
Effect of PfCullin-2 knock-down on sexual erythrocytic stage development. Synchronized PfCull2KD_3D7_ parasite cultures with (+T) or without (-T) trimethoprim were used to induce gametocytogenesis, and treated with NAG to eliminate asexual stages. **A.** The +T culture was evaluated for expression of PfCullin-2/cDD_HA_ in gametocytes by IFA using anti-HA antibodies. The images for the indicated gametocyte stages show PfCullin-2/cDD_HA_ signal (Cullin-2), nuclear staining (DAPI), parasite and erythrocyte boundaries in the infected cell (DIC), and overlap of all the three images (Merged). The size scale bars are shown in the merged image panels. **B.** Parasites were isolated from +T and –T cultures on day-4, and assessed for PfCullin-2/cDD_HA_ protein level by western blotting using anti-HA (ab-HA) and anti-β-actin (ab-AC) antibodies. The full-length PfCullin-2/cDD_HA_ is indicated with arrow, and the sizes of protein markers (M) are in kDa. **C.** The number of gametocytes/2000 cells in +T and -T cultures were estimated, and shown as the number of gametocytes over days in culture. The data is mean of three independent experiments with SD error bar.

### PfCullin-2 knock-down altered the levels of proteins associated with proteasome, lipid biosynthesis and transport, protein translation and folding, DNA repair, and apicoplast

Since CRLs are involved in ubiquitination-dependent turnover and signalling of a large number of proteins, we quantitated and compared the proteomes of 3^rd^ cycle PfCul2KD_3D7_ trophozoites cultured in the presence and absence of TMP by label free quantification (LFQ) method. PfCullin-2 was absent in the western blot as well as the mass spectrometry data of -TMP parasites (Figure 10A), which confirmed successful knock-down. Compared to the +TMP, - TMP parasite sample had 128 proteins downregulated (log2 fold change = ≤1), 199 proteins upregulated (log2 fold change = ≥1) and 557 proteins unchanged (log2 fold change between = ±1) in two biological replicates. Of these, 37 proteins were upregulated and 23 proteins were downregulated with a *P*-value of ≤0.05 (Figure 10B, Figure S10, Table S6). Some of the upregulated proteins are associated with proteasome and peptidases, transcription/translation and protein folding, ion transport, DNA replication and repair, and metabolism. The majority of down-regulated proteins are associated with lipid biosynthesis and transport, nucleoside/metabolite transport, and ribosome biogenesis. Proliferating cell nuclear antigen 2 (PCNA2) has been shown to be associated with DNA damage in malaria parasites [53], and it was elevated, suggesting DNA damage stress in PfCullin-2-depleted parasites. Seven apicoplast-associated proteins, a major site of fatty acid synthesis, and 8 proteins associated with lipid synthesis and transport were found to be dysregulated. Dysregulation of these proteins could compromise lipid biosynthesis and subsequently membrane integrity, which might be responsible for the disrupted morphology of PfCul2KD_3D7_ trophozoites/schizonts. Further investigation is required to understand how PfCRL4 is associated with the above pathways.

**Figure 10.**
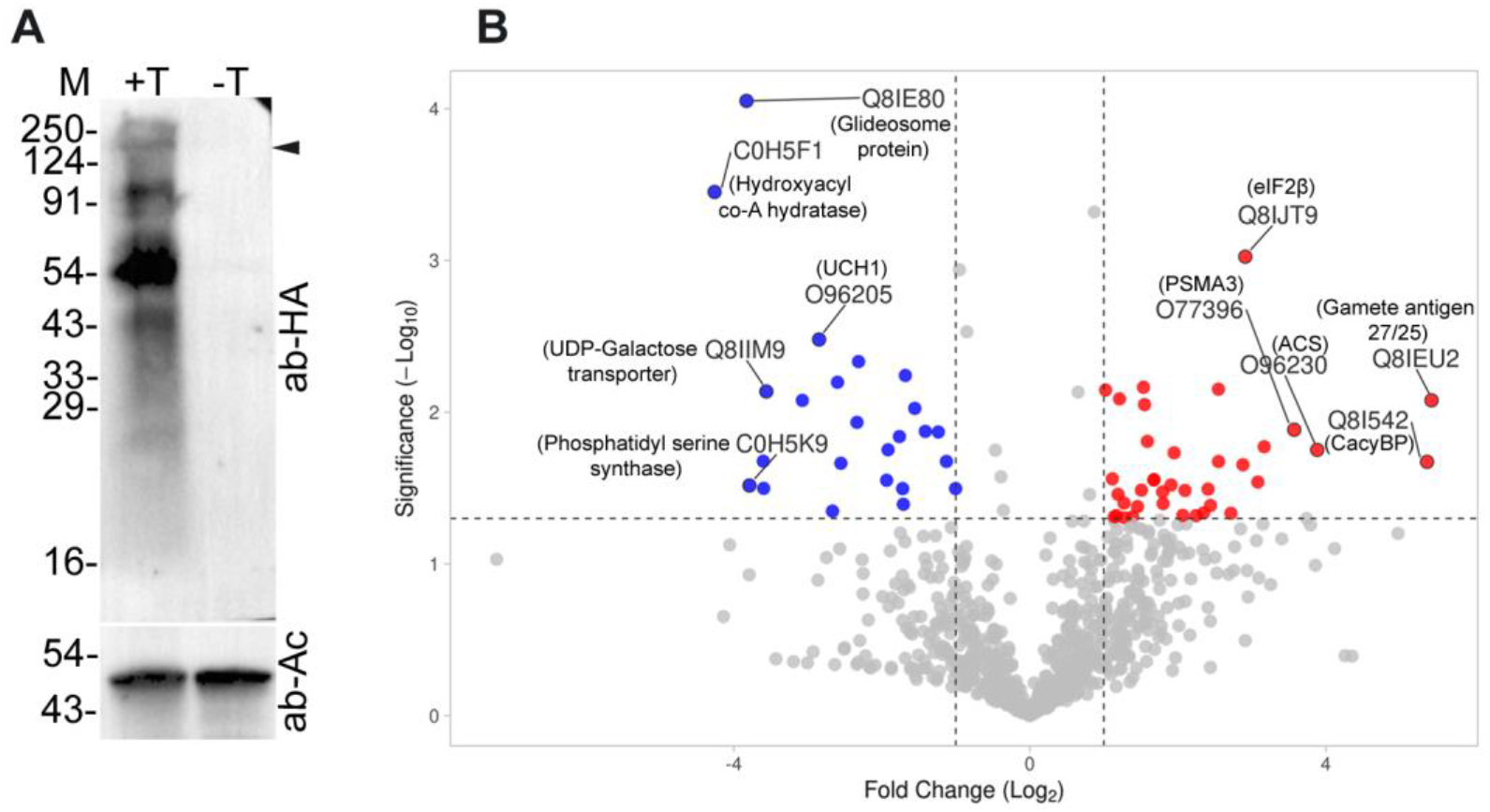
Changes in protein levels upon PfCullin2 knock-down. Synchronized PfCul2KD_3D7_ parasites were grown with (+T) or without (-T) trimethoprim for three cycles, and the lysates of 3^rd^ cycle trophozoites were compared for PfCullin-2/cDD_HA_ protein level by western blotting and changes in the proteome by label free quantification mass spectrometry. **A.** The western blot shows PfCullin-2/cDD_HA_ (ab-HA) and β-actin (ab-Ac) protein levels. The full-length PfCullin-2/cDD_HA_ is indicated with arrow, and the sizes of protein markers (M) are in kDa. **B.** The volcano plot shows up- and down-regulated proteins with at least 2-fold change (*P*-value of ≤0.05) in –T parasites. The X-axis shows Log2 fold change and Y-axis shows significance as -Log10 *P*-value (cut off 1.3 = 0.05).

### PfCullin-2 knock-down increased susceptibility to DNA damage and ER stress, and decreased mitochondrial membrane potential

The proliferating cell nuclear antigen 2 (PCNA2) was upregulated upon PfCullin-2-depletion, which is an indicator of DNA damage response [54], and suggested that these parasites might have genotoxic stress-like condition. We evaluated PfCul2KD_3D7_ parasites for susceptibility to the DNA-damaging agent methyl methanesulfonate (MMS), which has been shown to alkylate DNA [55]. PfCul2KD_3D7_ parasites were grown with or without TMP for 2 cycles and the 3^rd^ cycle trophozoite stage parasites were treated with DMSO or MMS, and the number of rings in each case was counted to assess recovery. The MMS-treated –TMP parasites produced almost 50% lesser number of rings than +TMP parasites (Figure 11A), indicating increased susceptibility to DNA-damaging agent upon PfCullin-2-depletion, which is in agreement with upregulation of PCNA2.

**Figure 11.**
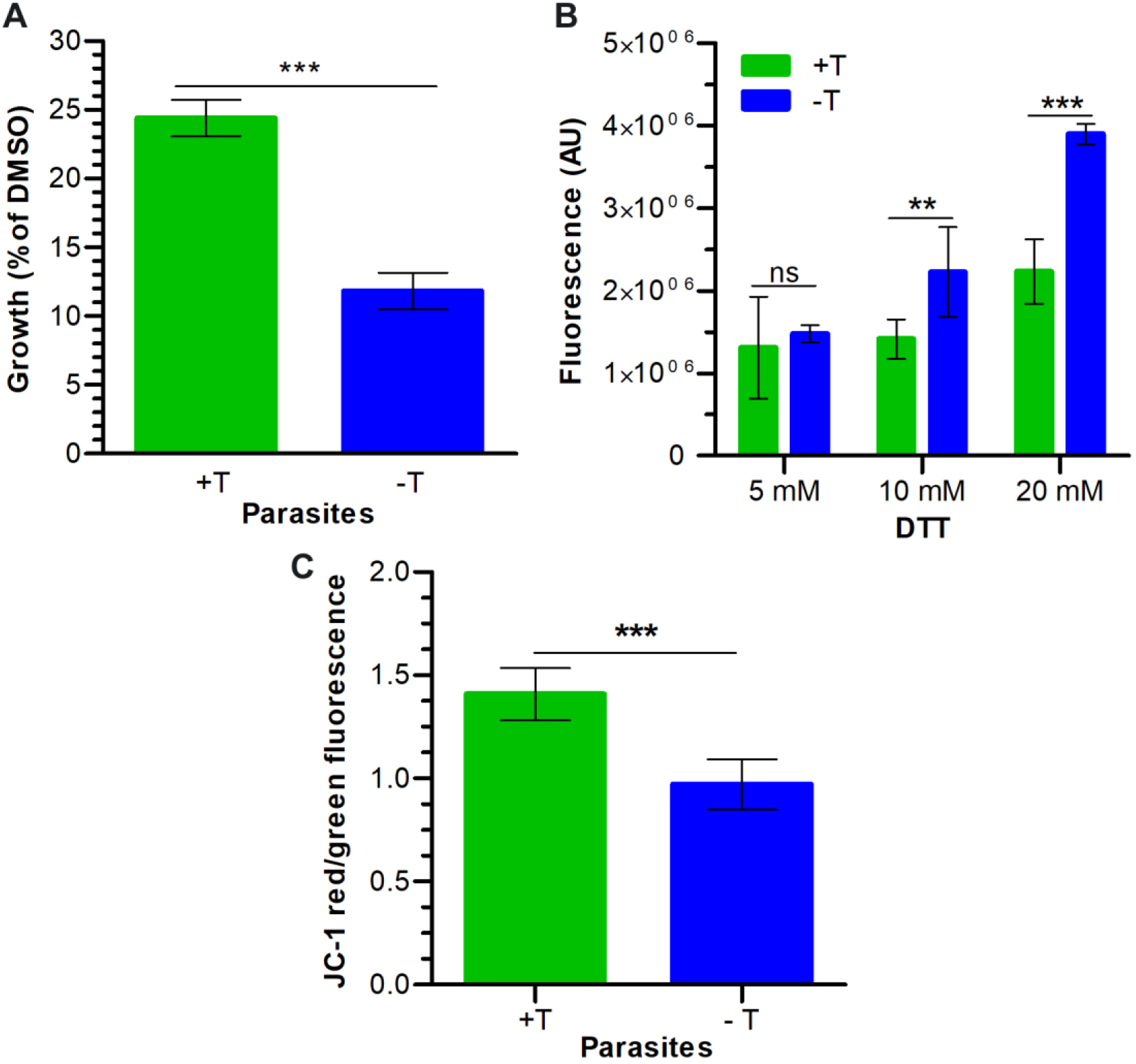
Effect of PfCullin-2 knock-down on susceptibility to stresses. PfCul2KD_3D7_ parasites were grown with (+T) or without (-T) trimethoprim for 2 cycles, and the 3^rd^ cycle trophozoite stage parasites were evaluated for mitochondrial membrane potential, and susceptibility to DNA-damaging agent and ER stress. **A.** The trophozoite stage parasites were treated with DMSO or the DNA-damaging agent MMS for 6 hours, and the number of fresh rings formed 12 hrs after the treatment was counted. The number of fresh rings in each culture is shown as % of its DMSO control for the same. **B.** The trophozoite stage parasites were treated with the indicated concentrations of DTT and ROS production was measured using H2DCFDA. The plot shows fluorescence values upon DTT treatment. **C.** The trophozoite stage parasites were stained with JC-1, and the JC-1 red and green fluorescence were measured. The plot shows the ratio of JC-1 red/green fluorescence (Y-axis) for the indicated parasites (X-axis). Each data is mean of three independent experiments with SD error bar.

PfCullin-2-depleted parasites had disrupted morphology and dysregulation of several proteins associated with lipid biosynthesis and transport, proteasome and peptidases, protein translation and folding, which suggested perturbed proteostasis and ER stress. DTT has been shown to induce ER stress in a variety of cells, including *Plasmodium* [11], which results in the production of reactive oxygen species (ROS) that converts non-fluorescent 5-(and-6)-carboxy-2’,7’-dichlorofluorescein diacetate (H_2_DCFDA) into fluorescent DCF, and this increase in fluorescence is directly proportional to ROS. Accordingly, PfCul2KD3_D7_ parasites were grown with or without TMP for 3 cycles, and the 3^rd^ cycle parasites were treated with DTT followed by with H_2_DCFDA. The PfCul2KD3_D7_ parasites grown without TMP showed more fluorescence than those grown with TMP condition, indicating increased ER stress upon PfCullin-2 depletion (Figure 11B).

PfCullin-2-depleted parasites also showed dysregulation of three proteins associated with mitochondria, which prompted us to assess these parasites for mitochondrial membrane potential as a measure of mitochondrial activity. For this, PfCul2KD_3D7_ parasites were grown with or without TMP for 2 cycles, the 3^rd^ cycle trophozoite stage parasites were stained with JC-1, and the red/green ratio was determined (Figure 11C). The –TMP parasites showed lower red/green ratio of J-aggregates than +TMP parasites, indicating decreased membrane potential, which is an indicator of increased mitochondrial stress.

### PfCullin-2 knock-down blocked cell division

PfCullin-2-depleted parasites showed significantly reduced growth, which prompted us to compare total DNA content and the number of nuclei/parasite in PfCul2KD_3D7_ parasites grown with or without TMP for 3 cycles. The DNA contents of +TMP and –TMP parasites increased as parasites matured and were largely similar until the 3^rd^ cycle ring stage (10 hr time point) (Figure 12A). As expected, the DNA content of +TMP parasites increased with maturation in the 3^rd^ cycle, whereas the DNA content of –TMP parasites was significantly lower than that of +TMP parasites (Figure 12A). Since the total DNA content does not accurately reflect the division state of parasite, we counted the number of nuclei/parasite at different time points during the 3^rd^ cycle. The number of multinucleated cell (2-4N to >12N) increased with the maturation of +TMP parasites, whereas the majority of -TMP parasites had single nucleus (1N) throughout the development, indicating replication arrest (Figure 12B).

**Figure 12.**
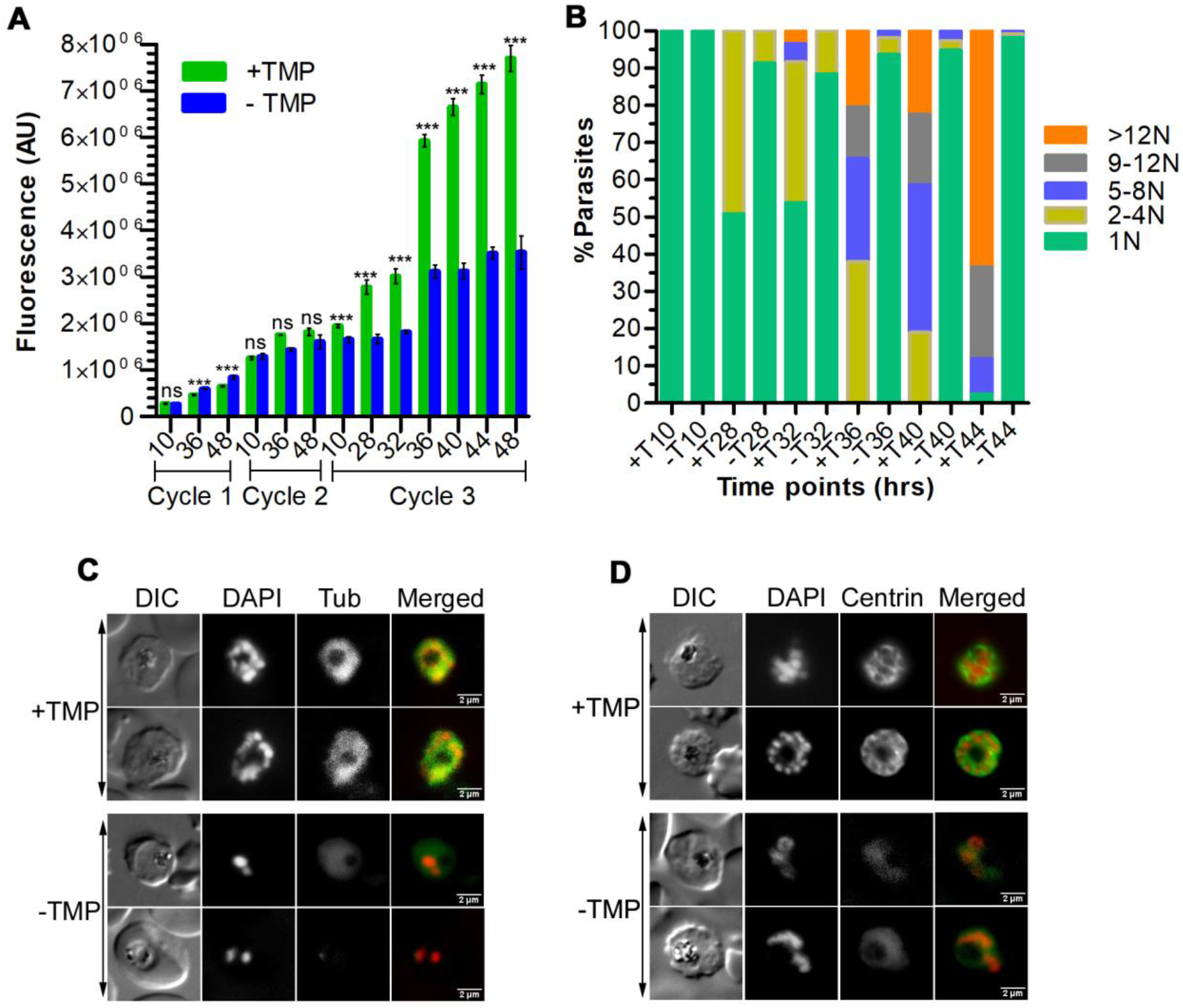
Effect of PfCullin-2 knock-down on cell division. Synchronized PfCul2KD_3D7_ parasites were grown with (+TMP) or without (-TMP) trimethoprim for three cycles, beginning with equal parasitemia. **A.** Equal aliquots of the cultures were collected at different time points over three cycles, stained with SYBR green I, and the fluorescence of each sample was measured. The plot shows fluorescent in arbitrary units (AU) on Y-axis for samples corresponding to the indicated time points of each cycle (X-axis). The data is from a representative of three experiments, each done in triplicate. “*” indicates *P*-value for the significance of difference between +TMP and –TMP data. **B.** The aliquots of 3^rd^ cycle cultures were smeared on glass slides, stained with DAPI, and the number of nuclei/parasite were counted using a microscope. The plot shows percentage of parasites with the indicated number of nuclei/parasite (Y-axis) at the indicated time points (X-axis). The data is based on at least 100 parasites for each time point from three experiments. **C and D.** Trophozoite/schizont stages during the 3^rd^ cycle were processed for IFA using anti-α-tubulin (C) or anti-centrin (D) antibodies. The panels are for the bright field image showing parasite and RBC boundaries (DIC), nuclear stain (DAPI), α-tubulin (Tub) or centrin, and the overlap of all three images (Merged). The size scale bar is in the merged image.

To further examine nuclear division, we examined the 3^rd^ cycle trophozoites/schizonts (40 hr time point) of +TMP and –TMP parasites for mitotic structures (spindle and microtubule organizing centre) using antibodies to α-tubulin and centrin-1, which have been used successfully earlier for studying *Plasmodium* cell division in [56–58]. The +TMP parasites showed prominent tubulin- and centrin-labeled structures in association with each nucleus, which were significantly diminished in -TMP parasites (Figure 12C, D). In particular, the +TMP parasites showed prominent tubular centrin signal with each nucleus, whereas the –TMP parasites had faint and diffuse centrin signal, suggesting that mitotic structures did not form upon PfCullin-2 depletion. The decreased DNA content, less number of nuclei/parasite and diminished mitotic structures in PfCullin-2-depleted parasites together indicate that PfCRL4 is required for cell division.

## Discussion

Cullin-RING ubiquitin E3 ligases (CRLs) form the largest group of ubiquitin E3 ligases, and have key roles in maintaining protein homeostasis, genomic integrity and fidelity of the cell division [28–30]. CRLs are associated with several human diseases and are the focus of many ongoing drug discovery efforts [59]. However, it remains to be investigated if *Plasmodium* cullins form functional complexes. Furthermore, the core components and roles of CRLs during the development of *Plasmodium* are not known. In this study, we demonstrate two functional CRLs in the human malaria parasite *P. falciparum*: a typical SCF-like (PfSCF) and a human CRL4-like (PfCRL4) ubiquitin E3 ligase. PfSCF comprises of PfCullin-1, PfSkp1, PfRbx1, PfFBXO1 and PfCacyBP. PfCRL4 appears to contain PfCullin-2, PfCPSF_A, two WD40 repeat proteins, and PfRbx1. PfCRL4 is crucial for cell division and membrane integrity, and likely contributes to proper functioning of mitochondria, ER, and genomic maintenance.

The homologs of cullins, Skp1 and Rbx1 have been predicted for *Plasmodium* and related Apicomplexan parasites (https://veupathdb.org/veupathdb/app; https://plasmodb.org/plasmo/app) [60]. We analyzed individual putative cullins, Skp1 and Rbx1 sequences of the representative Apicomplexan parasites to identify their closest *Plasmodium* and human counterparts. All the Apicomplexan parasites analyzed in this study have 2-3 putative cullins, one Skp1 and one Rbx1. We could not annotate Apicomplexan cullins as a particular human cullin homolog due to their low sequence identity with human cullins (<35%), which was also reflected as separation of *P. falciparum* cullins from those of model organisms on the phylogenetic tree. This suggests parasite-specific adaptation in function and composition of *P. falciparum* CRLs to regulate the complex and multi-stage parasite development. This is also supported by the absence of canonical CRL substrates like cyclins and CDKs in *P. falciparum* [61]. The presence of homologs for 2-3 cullins, Skp1 and Rbx1 in Apicomplexan parasites suggests that there are at least two CRLs, including an SCF-like ubiquitin E3 ligase.

PfCullin-1 and PfCullin-2 expression was predominant in trophozoite and schizont stages, which corresponds to their reported transcription profiles [46, 47]. The nucleocytoplasmic localization of PfCullin1, PfCullin-2, PfSkp1, and PfRbx1 suggests that CRLs containing these subunits function in both the compartments. Similar localization pattern and function in nucleus as well as cytoplasm have also been shown for most of the cullins and associated proteins in model systems [62].

Mass spectrometry of the immunoprecipitates followed by direct protein-protein interaction confirmed that PfCullin-1, PfSkp1, PfFBXO1, PfCacyBP and PfRbx1 form a SCF-like complex (PfSCF). PfSkp1 and PfRbx1 likely function as substrate adaptor and ubiquitin E2 recruiter, respectively. PfFBXO1 was present in the PfSkp1 immunoprecipitate only, interacted with PfSkp1, and has an F-box motif, indicating that it is an F-box protein and likely functions as a substrate receptor, as has been reported for F-box proteins [63]. PfCacyBP was present in the PfSkp1 immunoprecipitate and interacted with PfSkp1. Human CacyBP binds to multiple proteins, including Skp1, and has been shown to have roles in p53-induced degradation of non-phosphorylated human β-catenin [64], cytoskeleton organization, dephosphorylation of ERK1/2, cell proliferation and cancer development [65]. The homologs of p53, β-catenin and ERK1/2 appear to be absent in *Plasmodium*, suggesting that PfCacyBP has *Plasmodium*-specific roles.

PfCullin-2 likely forms a human CRL4B-like complex (PfCRL4), as several top hits (PfCullin-2, PfCPSF_A, PfRbx1 and WD40 repeat proteins) present in the PfCullin-2 immunoprecipitates share domain architecture with human CRL4B core subunits. The human CRL4B core subunits include HsCullin-4B, HsDDB1, HsDCAF1 and HsRbx1, which serve as a scaffold protein, substrate adaptor, substrate receptor and ubiquitin E2 recruiter, respectively [50]. PfCullin-2, like HsCullin-4B, has a long N-terminus upstream of the predicted cullin homology domain (CH) and a putative nuclear localization signal (NLS). The PfCPSF_A domain architecture is similar to that of HsDDB1 protein, and a BLAST search of the Uniprot database using PfCPSF_A sequence identified DDB1 proteins, suggesting that PfCPSF_A could be an HsDDB1 homolog of *P. falciparum*. HsDDB1 has been proposed to interact with several WD40 repeat proteins, including DCAF1, which have been shown or predicted to function as substrate receptors [66]. Several F-box proteins with WD40 repeats function as substrate receptors [63], and PfWD40 repeat proteins may serve as substrate receptors. PfCullin-2, PfCPSF_A and WD40 repeat proteins are significantly larger than their human CRL4B counterparts, and these extended regions are also present in their homologs in other *Plasmodium* species. We speculate that these extended regions might impart parasite-specific functions like interaction with parasite-specific proteins. Notably, PfCullin-2 immunoprecipitates also contained a putative ubiquitin carboxy-terminal hydrolase, which showed most similarity with ubiquitin carboxy-terminal hydrolase 7 (UCH7) proteins, and we have termed it the *P. falciparum* ubiquitin carboxy-terminal hydrolase 7 (PfUCH7). The *H. sapiens* ubiquitin specific protease 7 (USP7), also called the Herpes-Associated Ubiquitin-Specific Protease (HAUSP), has been shown to be involved in a variety of pathways and associated with multiple ubiquitin E3 ligases, including MDM2 and RAD18. This association has been shown to be important for regulation of the functions of HsUSP7 and the interacting ligases [67]. PfUCH7 or its *Plasmodium* orthologs have not been characterized yet to our knowledge; it may regulate PfCRL4 function by performing ubiquitin chain editing and/or deubiquitinating the PfCRL4 subunits, which agrees with autoubiquitination of PfCullin-2. On the other hand, PfCRL4 may impart specificity to PfUCH7 for substrates. Some other noteworthy proteins identified in the immunoprecipitate included subunits of the translation initiation factor 3 (A, C, D, E, I), Atg18, and proteins involved in nuclear transport (exportin 7 and karyopherin). This suggests a role of PfCRL4 in multiple processes, and some of these proteins could be substrates or modulators of PfCRL4.

Due to large sizes of most of the PfSCF and PfCRL4 subunits and AT-richness of their coding genes, we could not produce these subunits as recombinant proteins for assembling the complexes *in vitro* to perform functional assays. Hence, we evaluated PfSkp1 immunoprecipitate, which contained all the core subunits except Rbx1, together with recombinant Rbx1 for *in vitro* ubiquitination activity. Similarly, PfCullin-2 immunoprecipitate, which contained all the core CRL4 subunits, was assessed for ubiquitination activity. Both the immunoprecipitates showed ATP-dependent ubiquitination activity, indicating that PfSCF and PfCRL4 complexes are functional ubiquitin E3 ligases. Since the immunoprecipitates did not contain any other ubiquitin E3 ligase, the ubiquitination activity is of PfSCF and PfCRL4. PfCullin-2 appeared to undergo autoubiquitination, which may regulate PfCRL4 functions, as autoubiquitination has been shown to regulate the stability and functions of several ubiquitin E3 ligase [68]. Since the ubiquitinated proteins are the co-immunoprecipitated proteins, these could be the substrates of PfSCF or PfCRL4. Further investigation is needed to identify specific substrates of PfSCF and PfCRL4, which will provide insights into their specific targets. A previous study also identified Cullin-1, Skp1, Rbx1, and FBXO1 in the Skp1-immunoprecipitate of *T. gondii*, and proposed an SCF-like CRL in *T. gondii* [69]. Rashpa, *et. al.* in a recent study performed extensive immunoprecipitation of Rbx1, Skp1 and FBXO1 of the mouse malaria parasite *P. berghei*, and proposed that these proteins form an SCF in *P. berghei* [43]. While our results for PfSCF are in agreement with these two reports, we extended the composition, confirmed one-one interaction between different components, and demonstrated a functional SCF in the human malaria parasite *P. falciparum*. Furthermore, the identification of functional PfCRL4 in our study is the first report for *Plasmodium* to our knowledge, and a similar ubiquitin E3 ligase may be present in related Apicomplexan parasites.

We generated a trimethoprim-inducible PfCullin-2 knock-down line in *P. falciparum* 3D7 strain (PfCul2KD_3D7_) to investigate the roles of PfCRL4 during parasite development. PfCul2KD_3D7_ showed depletion of PfCullin-2 in the absence of trimethoprim (-TMP), which would prevent the formation of the PfCRL4 complex, thereby allowed us a comparative assessment of –TMP and +TMP parasites. The PfCullin-2-depleted parasites showed significantly decreased growth as compared with the parasites grown in the presence of trimethoprim (+TMP). The knock-down effect was partial in the 1^st^ cycle and became prominent in the 2^nd^ and 3^rd^ cycles. The partial knock-down effect could be due to essentiality of PfCullin-2 for parasite development, which has also been observed with other essential proteins [70, 71]. Despite near complete depletion of full-length PfCullin-2 in the 2^nd^ and 3^rd^ cycles, a significant reduction in parasite growth was observed in the 3^rd^ cycle and onward. We speculate that there could be some level of functional redundancy in PfCRL4 functions, which may compensate for it for a short period but not for prolonged parasite development. This is somewhat similar to the delayed death phenomenon observed in *P. falciparum* upon treatment with translation inhibitor antibiotics [72].

PfCullin-2 immunoprecipitate contained proteins belonging to various metabolic pathways and PfCullin-2-depletion caused dysregulation of proteins associated with a variety of pathways, particularly lipid biosynthesis and transport, protein translation and folding, DNA repair and nucleoside/metabolite transport. This might explain decreased mitochondrial membrane potential, and increased susceptibility to DNA damage and ER stress upon PfCullin-2-depletion. Notably, the levels of 7 apicoplast-associated proteins and 8 proteins associated with lipid synthesis/transport were highly dysregulated in PfCullin-2-depleted parasites. Apicoplast is an essential, non-photosynthetic plastid remnant that provides isoprenoid precursors, fatty acids and heme for parasite development [73–75]. One of the down-regulated apicoplast proteins is a CDC48-subfamily protein, _AP_CDC48, which has been proposed to be part of an apicoplast-localized ERAD-like complex [39, 40]. The *Toxoplasma* _AP_CDC48 has been shown to be required for protein translocation across the apicoplast membrane [76], and it is likely that *Plasmodium* _AP_CDC48 has roles in transport of nuclear-encoded proteins to the apicoplast and maintenance of apicoplast protein quality control. We speculate that dysregulation of the apicoplast-associated proteins and the proteins associated with lipid synthesis/transport would adversely affect apicoplast and lipid biosynthesis, thereby affecting the membrane integrity and biogenesis. Inhibition of apicoplast-associated processes has been shown to cause delayed parasite death in the subsequent cycles [77], which is similar to the delayed effects of PfCullin-2-depletion in our study. Further investigation is required to identify a direct link between PfCRL4 and apicoplast. A link between lipid metabolism and ER stress has been shown both at cellular and organismal level [78, 79], which is in-line with our results. Deletion of CRL4 *in S. cerevisiae*, mammalian cells, *C. elegans*, and plants has been shown to cause compromised DNA repair and replication [30, 50, 80], and this is in agreement with our data showing increased susceptibility of PfCullin-2-depleted parasites. These data suggest pleiotropic roles of CRL4, including lipid biosynthesis.

A large number of PfCullin-2-depleted trophozoites/schizonts were morphologically defective. In SEM, +TMP schizonts had multiple merozoites as lobules extending to the surface, whereas -TMP schizonts had fewer of such lobules. The TEM images of +TMP schizonts showed properly segmented merozoites with clearly demarcated outer membrane and internal membraned structures, whereas -TMP schizonts had fewer merozoites with poorly demarcated outer membrane and internal membraned structures. Decreased number of merozoites with poorly demarcated outer merozoite membrane and membraned structures in – TMP parasites indicate a defect in the formation and segregation of merozoites and suggest a key role of PfCRL4 in cell division. As has been explained earlier, the morphological abnormalities of –TMP parasites could be due to impaired lipid biosynthesis.

Compared to the +TMP parasites, PfCullin-2-depleted parasites had less total DNA content, fewer number of nuclei/parasite, and diminished mitotic structures, which confirmed a block in cell division. Nuclear localization and chromatin-association of PfCullin-2, the presence of several nuclear proteins in the PfCullin-2 immunoprecipitate, and altered levels of several nuclear proteins upon PfCullin-2 depletion suggest that PfCullin-2 regulates several nuclear processes (transport, replication, transcription, nucleotide synthesis, replication and repair). Hence, depletion of PfCullin-2 could have adversely affected these processes, causing a block in cell division. The effects of PfCullin-2 depletion were not noticeable in the 1^st^ and 2^nd^ cycles, which could be due to partial depletion of PfCullin-2 in the 1^st^ cycle and there could be some level of functional redundancy in PfCRL4 functions, which together with partial depletion of PfCullin-2 might be sufficient for parasite development in the first two cycles.

Rashpa, *et. al*. recently reported that genetic ablation or depletion of the *P. berghei* SCF component FBXO1 blocked cell division during erythrocytic schizogony, prevented gamete egress from the host erythrocyte, and affected integrity of the apical and the inner membrane complexes in merozoite and ookinete [43]. Our data showing key role of CRL4 in cell division and maintenance of membrane integrity during asexual erythrocytic development of *P. falciparum* complements the findings of Rashpa, *et. al*., and extends the CRL repertoire and the roles of *Plasmodium* CRLs. These two studies together indicate that *Plasmodium* SCF and CRL4 have essential, but non-overlapping, functions during the development of malaria parasites.

In summary, our study shows that *P. falciparum* Cullin-1 and Cullin-2 form functional PfSCF and PfCRL4 ligases, which likely have roles in the cytoplasmic, nuclear and DNA-associated processes. PfCRL4 is crucial for asexual and sexual erythrocytic development, and has key roles in cell division and maintenance of membrane integrity (Figure 13). Our demonstration of functional CRLs in the human malaria parasite *P. falciparum* will pave the way for similar studies in other Apicomplexan parasites, which is necessary to uncover the biology and drug-target potential of CRLs in parasitic protozoa.

**Figure 13.**
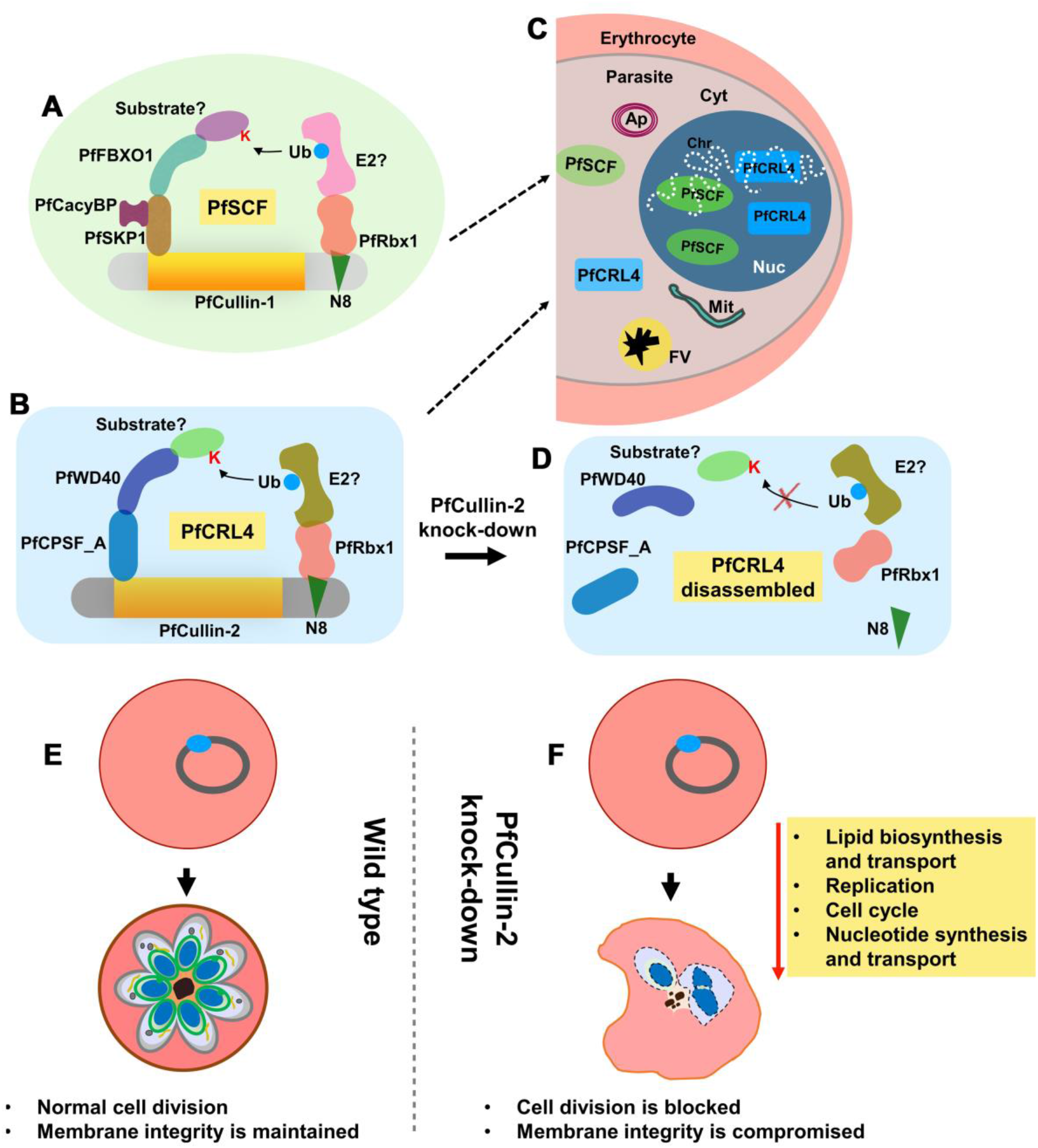
Highlights of the study. PfCullin-1 and PfCullin-2 serve as scaffolds for assembly of the indicated proteins into functional PfSCF (A) and PfCRL4 (B) ubiquitin E3 ligases, respectively. Both the complexes localize (C) to cytoplasm (Cyt) and nucleus (Nuc), and associate with chromatin (Chr), suggesting their roles in these compartments, including DNA-associated processes. Apicoplast (Ap), mitochondria (Mit) and food vacuole (FV) with haemozoin (black structure) are also shown in the parasite within the erythrocyte. PfCullin-2 knock-down at protein level would prevent assembly of the indicated subunits into functional PfCRL4 (D), thereby affecting the associated processes. Wild type asexual erythrocytic parasite developed normally from ring to multinucleate schizonts (E), whereas PfCullin-2 knock-down parasites exhibited downregulation of several key pathways, resulting in a block in cell division and compromised membrane integrity (F). Contrary to the wild type schizont that showed multiple nuclei with distinct outer membrane and intracellular membraned structures, the PfCullin-2 knock-down schizont had less number of nuclei and poorly demarcated membraned structures.

## Materials and Methods

### Materials

All the routinely used biochemicals were of molecular biology grade and procured from MERCK or similar sources unless otherwise stated. Cell culture reagents were from Lonza and Thermo Fisher Scientific. Restriction enzymes and DNA modifying enzymes were from New England Biolabs and Thermo Fisher Scientific. DNA isolation kits were from QIAGEN and MACHEREY-NAGEL. ProLong Diamond Antifade Mountant, Hoechst, DAPI, SuperSignal Chemiluminescent substrate and 2’,7’-dichlorodihydrofluorescein diacetate acetyl ester (H_2_DCFDA) were from Thermo Fisher Scientific. Secondary antibodies were from Thermo Fisher Scientific and Jackson ImmunoResearch. Primary antibodies were from Cell Signalling Technologies, Thermo Fisher Scientific, MERCK and ChromoTek. WR99210 was a kind gift from David Jacobus (Jacobus Pharmaceutical, Princeton, U.S.A.). Protease inhibitor cocktail, trimethoprim (TMP) and N-acetylglucosamine (GlcNAc) were from MERCK. Glutaraldehyde, formaldehyde, osmium tetroxide, lead citrate solution, and sodium cacodylate buffer were from Electron Microscopy Sciences. Uranyl acetate was from Loba Chemie and epoxy resin constituents were from TED PELLA. *P. falciparum* 3D7 and D10 strains were obtained from the Malaria Research and Reagent Reference Resource centre (MR4). All the plastic ware was from standard manufacturers such as Corning Inc, Nalgene, Thermo Fisher Scientific and Tarsons. Human blood was collected after obtaining written informed consent from the subjects according to the protocols (IEC-38/2015, IEC-38-R1/2015, IEC-38-R2/2015 and IEC-38-R3/2015) approved by the Institutional Ethics Committee (IEC) of Centre for Cellular and Molecular Biology. All the studies abide by the Declaration of Helsinki principles.

### Methods

#### Sequence analysis and phylogeny

For identification of cullin homologs in *Plasmodium falciparum*, the amino acid sequences of *Homo sapiens* cullins (HsCullin-1 (Q13616), HsCullin-2 (Q13617), HsCullin-3 (Q13618), HsCullin-4A (Q13619), HsCullin-4B (Q13620), HsCullin-5 (Q93034), HsCullin-7 (Q14999), and HsCullin-9 (Q8IWT3)) and *Saccharomyces cerevisiae* cullins (ScCullin-1 (YDL132W), ScCullin-3 (YGR003W), and ScCullin-8 (P47050)) were used as queries in BLAST searches of the *Plasmodium* genome database PlasmoDB (https://plasmodb.org/plasmo/app) and the NCBI non-redundant protein sequence database (https://blast.ncbi.nlm.nih.gov/Blast.cgi). For identification of Rbx1 and Skp1 homologs in *P. falciparum*, the amino acid sequences of *H. sapiens* Rbx1 (HsRbx1 (P62877)) and Skp1 (HsSkp1 (P63208)), and *Saccharomyces cerevisiae* Rbx1 (ScRbx1 (Q08273)) and Skp1 (ScSkp1 (P52286)) were used as queries in BLAST searches of the *Plasmodium* genome database PlasmoDB. To search for cullin, Rbx1 and Skp1 homologs in Apicomplexan parasites, amino acid sequences of *P. falciparum* cullin-1 (PfCullin-1), cullin-2 (PfCullin-2), Rbx1 (PfRbx1) and Skp1 (PfSkp1) were used as queries in the BLAST searches of the eukaryotic pathogen genomics database resource EuPathDB (https://veupathdb.org/veupathdb/app). Sequence alignment was performed using Clustal Omega (https://www.ebi.ac.uk/Tools/msa/clustalo/), and amino acid sequences were analysed for conserved domains and motifs using CD-BLAST program (https://www.ncbi.nlm.nih.gov/Structure/cdd/wrpsb.cgi). For phylogeny, amino acid sequences were aligned using the MAFFT program (https://www.ebi.ac.uk/Tools/msa/mafft/), the phylogenetic tree was constructed using IQ-TREE (http://iqtree.cibiv.univie.ac.at/) with 1000 bootstrap alignments [81, 82], and the tree was visualized with iTOL (https://itol.embl.de/).

#### Parasite culture

Whole human blood was collected, peripheral blood mononuclear cells were removed by washing the blood with RBC storage medium (RPMI 1640 supplemented with 2 g/l glucose, 25 mg/l gentamicin, 300 mg/l glutamine), and RBCs were stored at 50% haematocrit in RBC storage medium at 4°C until used. *P. falciparum* 3D7 and D10 strains were grown at 37°C in a mixed gas environment (5% CO_2_, 5% O_2_ and 90% N_2_) in RPMI 1640-albumax medium with human erythrocytes at 2% haematocrit (RPMI 1640 supplemented with 2 g/l sodium bicarbonate, 2 g/l glucose, 25 mg/ml gentamicin, 300 mg/l glutamine, 100 mM hypoxanthine, 0.5% albumax II) [83]. Parasites were synchronized by sorbitol treatment when the majority of parasites were at ring stage [84], and parasites were isolated from infected RBCs by saponin lysis. Parasite pellets were used immediately in experiments or stored at −80°C until further use. Genomic DNA was isolated from late trophozoite/schizont stage parasites using the Puregene Blood Core kit as recommended by the manufacturer.

#### Construction of transfection plasmids

The sequences of primers used in this study are mentioned in Table S7. The complete open reading frames of PfRbx1 and PfSkp1 were PCR amplified from *P. falciparum* genomic DNA using primers PfRbx-F/PfRbxmyc-R and PfSkp1-F/PfSkp1-R, respectively. PfRbxmyc-R contains Myc-tag coding sequence for addition to the C-terminus of PfRbx1. The PfRbx1myc PCR fragment was cloned into the pGEM-T vector, and subcloned into the pNC plasmid at BglII/XhoI sites to obtain pNC-PfRbx1myc transfection plasmid; pNC has been described earlier [44]. The PfSkp1 PCR fragment was cloned into the pGT-GFP vector at BglII/KpnI sites upstream of the GFP coding sequence [85], excised with BglII/XhoI to obtain PfSkp1/GFP, which was cloned into the similarly digested pNC vector to obtain pNC-PfSkp1/GFP transfection plasmid. PfCullin-1 locus was targeted for single cross over homologous recombination. The PfCullin-1 coding region corresponding to C-terminal 500 amino acid residues was PCR amplified from *P. falciparum* 3D7 genomic DNA using PfCul1Fki/PfCul1Rki primers, digested with ApaI/AgeI, and cloned into the similarly digested pNC-GFP vector upstream of the GFP coding sequence to obtain pNC-PfCullin-1/GFPki transfection plasmid. Double cross-over homologous recombination approach was used to target the PfCullin-2 locus. The PfCullin-2 coding sequence corresponding to C-terminal 409 amino acid residues (flank1) and 867 bp 3’-UTR of PfCullin-2 (flank2) were PCR amplified from *P. falciparum* 3D7 genomic DNA using PfCullin2F1kiFP/PfCullin2F1kiRP and PfCullin2F2F/PfCullin2F2R primers, respectively. The flank1 and flank2 PCR fragments were sequentially cloned into the HBA18/cDD_HA_kd plasmid at NotI-KpnI and AvrII-KasI sites, respectively, to obtain HB-PfCul2/cDD_HA_ plasmid. The HBA18/cDD_HA_kd plasmid has been described earlier [86]. A synthetic gene coding for the *Bacillus subtilis* glmS ribozyme and PvAc-3’U was obtained from (GenScript), and cloned into the HB-PfCul2/cDD_HA_ plasmid at XhoI/AgeI site, replacing the existing PvAc-3’U fragment to generate HBglm-PfCul2/cDD_HA_ transfection plasmid. The coding regions and flanks in all the transfection plasmids were sequenced to ensure that they were free of undesired mutations, and the presence of different regions was confirmed by digestion with region-specific restriction enzymes. The transfection plasmids were purified using the NucleoBond® Xtra Midi plasmid DNA purification kit.

#### Transfection of parasites

Freshly thawed cultures of *P. falciparum* D10 and 3D7 strains were used for transfection. *P. falciparum* D10 early ring stage parasites were transfected with 50-100 μg of the desired transfection plasmid (pNC-PfRbx1myc, pNC-PfSkp1/GFP, pNC-PfCullin-1/GFPki, and HBglm-PfCul2/cDD_HA_) by the infected RBC (iRBC) method [87, 88]. Similarly, *P. falciparum* 3D7 early ring stage parasites were transfected with HBglm-PfCul2/cDD_HA_ transfection plasmid. The transfected cultures were maintained in the presence of 1 μg/ml blasticidin (pNC-PfRbx1myc, pNC-PfSkp1/GFP and pNC-PfCullin-1/GFPki) or 1 nM WR99210 (HBglm-PfCul2/cDD_HA_) from day 2-12, followed by no drug till day 18, and then again in the presence of the drug until the emergence of resistant parasites. For pNC-PfCullin-1/GFPki and HBglm-PfCul2/cDD_HA_ transfected parasites, genomic DNA was isolated at every at 3-4 weeks intervals and assessed for plasmid integration into the target site by PCR using primer sets specific for 5’-integration (PfCullin-1: Cull1-FseqP3/GFPSeqR1, PfCullin-2: PfCul2-5con/PvAc-Rseq), 3’-integration (PfCullin-1: PcDT5U-RP/PfCul1-3con, PfCullin-2; Hrp2-seqF/PfCul2-3con), wild type locus (PfCullin-1: Cull1FseqP1/PfCul1-3con, PfCullin-2: PfCullin2-5con/PfCullin2-3con) and a positive control target (PfCullin-1: PfRbxF/PfRbxR, PfCullin-2: PfAtg18-5con/3con). Once plasmid integration at the target locus was observed (transfected with pNC-PfCullin-1/GFPki or HBglm-PfCul2/cDD_HA_), parasites were cloned by dilution cloning method, and cloned lines were again evaluated for the presence of integration locus by PCR. A GFP-expressing parasite line (GFPcon) was used as a control in various experiments, which has been described earlier [86].

#### Expression and localization of proteins

Recombinant parasites carrying the pNC-PfRbx1myc (PfRbx1myc-epi) or pNC-PfSkp1/GFP (PfSkp1GFP-epi) plasmid would express PfRbx1myc or PfSkp1GFP fusion protein, respectively. Cloned recombinant parasites with integration at the PfCullin-1 locus (PfCul1GFPKI) of PfCullin-2 locus (PfCul2KD_D10_ and PfCul2KD_3D7_) would express PfCullin-1/GFP or PfCullin-2/cDD_HA_ fusion protein, respectively. Recombinant parasites were evaluated for expression and localization of the target fusion proteins. Mixed stage cultures of wild type *P. falciparum* D10, GFPcon, PfRbx1myc-epi and PfSkp1GFP-epi parasites were harvested at 10% parasitemia. The parasite pellets were resuspended in 4× pellet volume of the SDS-PAGE sample buffer (1× buffer contains 50 mM Tris-HCl, 20% glycerol, 2% SDS, 1% β-mercaptoethanol, 0.01% bromophenol blue, pH 6.8), the lysates were run on 12% SDS-PAGE, and the proteins were transferred onto the Immobilon-P membrane. The membrane was blocked (blocking buffer contained 3% skim milk in TBS-T (10 mM Tris, 15 mM NaCl, 0.1% Tween-20, pH 7.4)), incubated with mouse anti-Myc (for PfRbx1myc) or rabbit anti-GFP (for PfSkp1GFP) antibodies (at 1/1000 dilution in blocking buffer), followed by HRP-conjugated goat anti-mouse or anti-rabbit IgG (at 1/20000 dilution in blocking buffer). The signal was developed using the SuperSignal^TM^ Chemiluminescent Substrate. The blot was stripped and re-probed for β-actin as a loading control with mouse anti-β-actin-HRP antibodies (at 1/30000 dilution in blocking buffer).

Cultures of the PfCul1GFPKI, PfCul2KD_D10_ and PfCul2KD_3D7_ parasites were synchronized and harvested at ∼10% parasitemia at different time points during the 48 hour-long asexual erythrocytic development. The parasite pellets were processed for western blotting using mouse anti-HA (for PfCul2KD_D10_ and PfCul2KD_3D7_, at 1/1000 dilution in blocking buffer) or rabbit anti-GFP (for PfCul1GFPKI, at 1/1000 dilution in blocking buffer) antibodies, followed by HRP-conjugated goat anti-mouse or anti-rabbit IgG (at 1/20000 dilution in blocking buffer) as has been described above for PfRbx1myc-epi and PfSkp1GFP-epi parasites. The blot was stripped and re-probed for β-actin with mouse anti-β-actin-HRP antibodies (at 1/30000 dilution in blocking buffer).

For localization of target proteins by immunofluorescence assay (IFA), synchronized cultures of recombinant parasites (PfRbx1myc-epi, PfCul2KD_D10_ and PfCul2KD_3D7_) were collected at various stages during the 48 hour-long asexual erythrocytic development, washed with PBS, immobilized on a poly L-Lysine coated slide, fixed (3% paraformaldehyde and 0.01% glutaraldehyde), permeabilized with 0.1% Triton X-100 (v/v in PBS), and blocked (blocking buffer contained 3% BSA in PBS). The slides were incubated with mouse anti-Myc or mouse anti-HA (1/100 in blocking buffer) antibodies, followed by Cyanine 3-conjugated goat anti-mouse IgG (at 1/1000 dilution in blocking buffer) and DAPI (10 µg/ml). The slides were air-dried, mounted with ProLong Gold anti-fade reagent, and covered with a coverslip. The edges of the mounted area were sealed with nail polish, the slide was observed under 100× objective of ZEISS Axioimager microscope fitted with Zeiss AxioCam HRm camera to capture images using Zeiss AxioCam HRm, which were analysed using the Axiovision software. For live-cell fluorescence microscopy, different stages of PfSkp1GFP-epi and PfCul1GFPKI parasites were collected, immobilized on a poly L-Lysine coated slide, and stained with Hoechst (1 μg/ml). The slides were observed, images were captured and analysed as has been described for IFA. For confocal microscopy, the IFA slides (PfRbx1myc-epi and PfCul2KDD_10_) and fixed slides (PfSkp1GFP-epi and PfCul1GFPKI) were viewed using the Leica TCS SP8 confocal laser scanning microscope, Z-section images were captured, and edited using the Leica Application Suite, Fiji and Adobe Photoshop softwares.

#### Fractionation of recombinant parasites

We prepared cytosolic and chromatin fractions of recombinant parasites by chromatin enrichment for proteomics (ChEP) method [89], and evaluated the fractions for the presence of target proteins by western blotting. Briefly, a 50 ml synchronous culture (at ∼10% parasitemia) of each of the PfRbx1myc-epi, PfSkp1GFP-epi, and PfCul1GFPKI parasites was grown in the presence of 1 μg/ml blasticidin till 30-hour stage, and parasites were purified by saponin lysis method. Similarly, a 50 ml synchronous culture of PfCull2KD_D10_ parasites was grown with 10 μM trimethoprim till 30-hour stage, and parasites were purified by saponin lysis method. The parasite pellets were fixed (1% formaldehyde in PBS), and treated with 0.125 M glycine (in PBS) to quench the reactive groups. The cells were resuspended in 500 µl of nuclear extraction buffer (25 mM Tris-Cl, 10 mM KCl, 0.1 mM EDTA, 0.1 mM EGTA, 1 mM DTT, pH 7.4, protease inhibitor cocktail, and 0.25% NP-40), the lysate was passed twice through a 25G needle, centrifuged at 2300g for 20 min at 4°C, and the supernatant was saved as a cytosolic fraction. The pellet, which contains nuclei, was washed with 500 µl of nuclear extraction buffer with RNAse A (200 µg/ml) followed by twice with PBS. The pellet was washed with Tris-SDS buffer (50 mM Tris-Cl, 10 mM EDTA, pH 7.4, 4% SDS, protease inhibitor cocktail) and urea buffer (10 mM Tris-Cl, 1 mM EDTA, 8 M urea, pH 7.4). The final pellet was resuspended in 500 µl of storage buffer (10 mM Tris-Cl, 1 mM EDTA, 25 mM NaCl, 10% glycerol, pH 7.4, and protease inhibitor cocktail), and the suspension was sonicated (Diagenode Bioruptor UCD-200) at 4°C for 5 min to solubilize chromatin completely. The sonicated sample was centrifuged at 15000g for 30 minutes at 4°C, the supernatant containing solubilized chromatin was mixed with equal volume of 2× SDS-PAGE sample buffer, and incubated at 98°C for 30 min to reverse the cross-links. The chromatin and cytosolic samples were assessed for purity by western blotting using rabbit anti-Histone 2B (1/1000 diluted in blocking buffer) and rabbit anti-α2 subunit of 20S proteasome (at 1/1000 dilution in blocking buffer) antibodies, followed by appropriate secondary antibodies. Both chromatin and cytosolic samples were evaluated for the presence of PfRbx1myc, PfSkp1GFP, PfCullin-1/GFP and PfCullin-2/cDD_HA_ proteins by western blot using mouse anti-Myc (PfRbx1myc), anti-HA (PfCullin-2/cDD_HA_) or anti-GFP (PfSkp1GFP, PfCullin-1/GFP) antibodies, followed by appropriate HRP-conjugated secondary antibodies as has been described in the expression and localization of proteins section.

#### Immunoprecipitation and mass spectrometry

Asynchronous cultures of PfRbx1myc-epi, PfSkp1GFP-epi, PfCul1GFPKI, wild type *P. falciparum* D10 and GFP control parasites were harvested at 10-15% parasitemia, and processed for immunoprecipitation as has been described earlier [86]. Briefly, the parasite pellets were resuspended in 5× pellet volume of the lysis buffer (10 mM Tris, 0.5 mM EDTA, 100 mM NaCl, 0.5% NP-40, pH 7.5, EDTA free protease inhibitor cocktail), subjected to freeze-thaw, the suspension was passed through a 26.5G needle, and the sample was incubated in ice for 30 min. The lysate was centrifuged at 25000g for 30 min at 4°C, the supernatant was transferred to a fresh tube, the pellet was re-extracted with 3× pellet volume of the lysis buffer and this supernatant was combined with the first supernatant. The supernatant was incubated with Myc-TRAP (PfRbx1myc-epi and wild type *P. falciparum* D10) or GFP-TRAP (PfSkp1GFP-epi, PfCul1GFPKI, and GFP control) antibodies (15 µl slurry/2 mg protein; ChromoTek) for 2 hours at 4°C with gentle shaking. The flow through was discarded, and the beads were washed with wash buffer 1 (10 mM Tris, 100 mM NaCl, 0.5 mM EDTA, pH 7.5, EDTA free protease inhibitor cocktail) followed by wash buffer 2 (10 mM Tris, 150 mM NaCl, 0.5 mM EDTA, pH 7.5, EDTA free protease inhibitor cocktail). The beads were boiled in 100 µl of 2× SDS PAGE sample buffer for 15 min to elute the bound proteins. A 20 µl aliquot of the eluate along with appropriate controls (input, flow through and washes) were checked for the presence of bait by western blotting using mouse anti-Myc or rabbit anti-GFP antibodies, followed by with appropriate secondary antibodies as described in the expression and localization of proteins section. The remaining eluate was processed for LC-MS/MS.

Asynchronous cultures of wild type *P. falciparum* D10 and PfCull2KD_D10_ (cultured with 10 μM trimethoprim) parasites were harvested at 10-15% parasitemia, the parasite pellets were lysed (lysis buffer: 10 mM Tris-Cl, 150 mM NaCl, 0.5 mM EDTA, 0.1% NP-40, 10% glycerol, pH 7.5, protease inhibitor cocktail), and processed further as has been described above. The supernatant was incubated with 5 μg of rabbit anti-HA antibodies for overnight with gentle mixing, the suspension was mixed with 10 μl of protein A/G magnetic beads (pre-equilibrated with wash buffer (10 mM Tris, 150 mM NaCl, 0.5 mM EDTA, 0.1% NP-40, 10% glycerol, pH 7.5)) for 2 h at 4°C, and processed further until the elution step as has been described above. A 20 µl aliquot of the eluate along with appropriate controls (input, flow through, washes) were assessed for the presence of PfCullin-2/cDD_HA_ by western blotting using mouse anti-HA antibody, followed by appropriate secondary antibody. The remaining eluate was processed for LC-MS/MS.

For LC-MS/MS, 80 µl of the eluate was stacked on a gradient SDS-PAGE gel until the protein marker ladder started resolving. The protein band was excised, digested with trypsin, peptides were extracted, vacuum dried, resuspended in 11 µl of 2% formic acid, and a 10 µl of the peptide sample was run on the Q Exactive HF (Thermo Fischer Scientific) to perform HCD mode fragmentation and LC-MS/MS analysis as has been described earlier [86]. The raw data files were acquired on the proteome discoverer v2.2 (Thermo Fischer Scientific), analysed, and the predicted peptide sequences were searched against the Uniprot database of *P. falciparum* 3D7 using the Sequest HT algorithm. The analysis criteria used were trypsin specificity, maximum two missed cleavages, and some variable modifications (oxidation of methionine, carbamidomethylation of cysteine, deamidation of asparagine/glutamine). Other criteria were precursor tolerance of 5 ppm, fragmentation tolerance of 0.05 Da, and 1% peptide FDR threshold. The protein hits from Rbx1myc and PfCullin-2/cDD_HA_ data were compared with those from the wild type *P. falciparum* D10 data. The proteins identified in PfSkp1GFP and PfCullin-1/GFP immunoprecipitates were compared with the proteins identified in GFPcon sample. Proteins exclusively present in PfRbx1myc, PfSkp1GFP, PfCullin-1/GFP, and PfCullin2/cDD_HA_ samples were considered genuine if reproducibly present in at least 2 of the 3 biological replicates with minimum 1 unique peptide.

#### Production of recombinant proteins

The complete coding regions of PfRbx1, *P. falciparum* calcyclin binding protein (PfCacyBP), PfSkp1, and the N-terminal coding region of PfCullin-1 (PfCul1N, 1-300 aa) were PCR amplified from *P. falciparum* genomic DNA using PfRbx-F/PfRbx-R, PfCal-Fexp/PfCal-Rexp, PfSkp1-Fexp/PfSkp1-Rexp, PfNCul1-F/PfNCul1-R primers, respectively. The PfCullin-1 C-terminal coding region (PfCul1C, 601-829 aa) and complete coding region of PfFBXO1 were amplified from *P. falciparum* cDNA using PfC-Cul1-Fexp/PfC-Cul1-Rexp and PfFbox-Fexp/PfFbox-Rexp primers, respectively. The PCR fragments were cloned into the pJET1.2 vector, sequenced to ensure that they were free of mutations, and subcloned into the pGEX-6P-1 expression vector at BamHI/XhoI site to obtain pGEX6P1/PfRbx1, pGEX6P1/PfCul1N, pGEX6P1/PfCul1C, pGEX6P1/PfCacyBP, pGEX6P1/PfSkp1, and pGEX6P1/PfFBXO1 expression plasmids. These expression plasmids were transformed into BL21(DE3) *E. coli* cells, which would express the recombinant proteins with an N-terminal GST-tag (GST/Cul1N, GST/Cul1C, GST/FBXO, GST/CacyBP, and GST/Rbx). We also transformed BL21(DE3) *E. coli* cells with pGEX6P-1 to produce GST protein. GST, GST/Rbx, GST/Cul1N and GST/Cul1C were purified from the soluble fraction of the respective IPTG-induced cells under native conditions using the Protino® Glutathione Agarose 4B resin as recommended by the manufacturer and reported earlier [86]. Briefly, the induced cell pellet was resuspended in lysis buffer (PBS with 1 mM DTT and 1 mg/ml lysozyme), the lysate was sonicated (5 s pulses at 20% amplitude for 5–30 min depending on the sample volume), centrifuged at 18000g for 30 min at 4°C, and the supernatant was incubated with Protino® Glutathione Agarose 4B. The resin was washed with PBS and bound proteins were eluted (50 mM Tris, 20 mM GSH, pH 7.5). Elution fractions were run on 12% SDS–PAGE, and fractions containing pure or enriched in recombinant protein were pooled and concentrated (30 kDa cut off Amicon Ultra centricon) with simultaneous buffer exchange to 20 mM Tris–Cl, 50 mM NaCl, pH 8.0 at 4°C. The protein amount in the concentrated sample was estimated by BCA assay, and the proteins were stored at -80°C until further use.

GST/FBXO and GST/CacyBP were purified under denaturing conditions from the respective IPTG-induced cells. Briefly, the induced cell pellet was resuspended in lysis buffer (PBS with 10 mM DTT and 8M urea; 5 ml/g weight of the pellet), sonicated (5 s pulses at 20% amplitude for 5–30 min depending on the sample volume), centrifuged, and the supernatant was dialysed sequentially against decreasing concentrations of the urea buffer (6 M, 5 M, 4 M, 3 M, and 2 M urea in PBS with 10 mM DTT, each for every 2 hours at 4°C). The dialysed sample was incubated with Protino® Glutathione Agarose 4B (pre-equilibrated with 2 M urea in PBS with 10 mM DTT, 0.125 ml resin/5g of initial cell pellet), the resin was washed with 2 M urea buffer, and bound proteins were eluted (2M urea buffer in PBS with 100 mM GSH). Elution fractions were run on 12% SDS–PAGE, and fractions containing pure or enriched in the recombinant protein were pooled, concentrated (30 kDa cut off Amicon Ultra centricon) with simultaneous buffer exchange to 20 mM Tris-Cl, 50 mM NaCl, pH 8.0 at 4°C, the protein amount was estimated by BCA assay, and stored at -80°C until further use PfSkp1 was also subcloned into the pET28a at BamHI/XhoI sites to obtain pET28a-PfSkp1, which was transformed into Rosetta-gami 2(DE3) pLysS cells to express with N-terminal His-tag (_His_Skp). Recombinant _His_Skp was purified from the soluble fraction of IPTG-induced cells using Ni-NTA chromatography under native conditions as has been described earlier [90]. The eluates were run on a 12% SDS-PAGE, fractions containing pure proteins were pooled and concentrated (10 kDa cut off Amicon Ultra centricon) with simultaneous buffer exchange to 20 mM Tris-Cl, 50 mM NaCl, pH 8.0 at 4°C. The protein amount was estimated by BCA, and the concentrated protein sample was stored at -80°C until further use.

PfRbx1 was also subcloned into pSMO at StuI/HindIII sites to obtain pSMO-PfRbx1 plasmid and transformed into Rosetta-gami 2(DE3) pLysS cells, which would express it with an N-terminal His-SUMO-tag (_His_SUMO/Rbx). The pSMO vector has been reported earlier [44]. The recombinant _His_SUMO/Rbx protein was purified from the soluble fraction of IPTG-induced cells by Ni-NTA affinity chromatography under native conditions as has been described earlier [44]. Elution fractions containing the pure protein were pooled and concentrated (10 kDa cut off Amicon Ultra centricon) with simultaneous buffer exchange to 20 mM Tris-Cl, 50 mM NaCl, pH 8.0 at 4°C. The protein amount was estimated by BCA, and the concentrated protein sample was stored at -80°C until further use. In parallel, we also transformed Rosetta-gami 2(DE3) pLysS cells with pSMO plasmid, and purified the _His_SUMO protein as has been reported previously [44].

#### Dot blot protein overlay assay

For assessing interaction of recombinant _His_PfSkp1 with recombinant GST, GST/PfRbx1, GST/PfCacyBP, GST/PfCul1N and GST/FBXO1 proteins, different amounts of purified _His_PfSkp1 were spotted on the nitrocellulose membrane, the membrane was blocked (3% BSA in PBS-T), and then overlaid with 10 μg of purified GST, GST/PfRbx1, GST/PfCacyBP, GST/PfCul1N or GST/FBXO1 for 12 h at 4°C with gentle shaking. The membrane was washed with PBS-T, incubated with mouse anti-GST antibodies (at 1/5000 in PBS-T), followed by HRP conjugated goat anti-mouse antibodies. The signal was developed using the SuperSignal West Pico Chemiluminescent kit. For interaction of recombinant _His_SUMO/PfRbx1 with recombinant GST/PfCul1C, GST/PfCacyBP, GST/PfFBXO1 and GST proteins, different amounts of purified _His_SUMO/PfRbx1 were spotted on the nitrocellulose membrane, the membrane was blocked, overlaid with 10 µg of individual recombinant protein, and processed further as has been described above for _His_PfSkp1. As a control to rule out interaction between GST and SUMO tags, different amounts of recombinant _His_SUMO protein was spotted on the nitrocellulose membrane, the membrane was blocked, and overlaid with 10 µg of recombinant GST/PfCul1C or GST protein, and processed further as has been described above for GST/PfSkp1.

### Co-immunoprecipitation of PfCullin-2 and PfRbx1

Asynchronous cultures of PfRbx1myc-epi and PfCul2KD_D10_ parasites were harvested at ∼15% parasitemia, the parasites were purified by the saponin lysis method, and the parasite pellets were resuspended in 4× pellet volume of the lysis buffer (10 mM Tris-Cl, 150 mM NaCl, 0.5 mM EDTA, 0.1% NP-40, 10% glycerol, pH 7.5, protease inhibitor cocktail). Equal amounts of the lysates were mixed, supplemented (with 2 mM ATP, 5 mM MgCl_2_ and 0.2 mM DTT), incubated at room temperature for 2 hrs with gentle mixing, and processed for immunoprecipitation using Myc-TRAP antibodies as has been described in the immunoprecipitation section. The eluate and appropriate controls (input, flow through, and washes) were assessed for the presence of PfRbx1myc by western blotting using mouse anti-Myc antibodies, followed by appropriate secondary antibodies as has been described in the western blotting section. The blot was stripped and re-probed to check for co-immunoprecipitation of PfCullin-2/cDD_HA_ using rabbit anti-HA antibodies, followed by appropriate secondary antibodies as has been described in the expression and localization of proteins section.

### Ubiquitination activity assays of PfSCF and PfCRL4

PfSkp1GFP-epi and PfCull2KD_D10_ parasites were processed for immunoprecipitation of PfSkp1GFP and PfCullin-2/cDD_HA_, respectively, before the elution step as has been described in the immunoprecipitation section. The bead-bound PfSkp1GFP and PfCullin-2/cDD_HA_ immunoprecipitates were assessed for ubiquitination activity using a ubiquitin conjugation initiation kit according to the manufacturer’s instructions (Boston Biochem). Briefly, each bead-bound immunoprecipitate was resuspended in 36 μl of the supplied ubiquitination reaction containing ubiquitin E1 enzyme, ubiquitin E2 enzyme, ubiquitin, and 0.5 mM DTT. 1 μg of recombinant GST/PfRbx1 was also added to the PfSkp1GFP ubiquitination reaction. Each reaction was divided into two equal aliquots, one was supplemented with MgCl_2_ and ATP and the other was supplemented with water to make up the volume to 20 μl. Both the reactions were incubated at 37°C for 1 h, stopped by adding 20 µl of 4× SDS-PAGE sample buffer, boiled for 15 minutes, and the supernatants were processed for western blotting using mouse anti-ubiquitin antibodies, followed by with HRP-conjugated goat anti-mouse IgG (at 1/20000 dilution in blocking buffer). The blots were stripped and reprobed for the presence of PfSkp1GFP or PfCullin-2/cDD_HA_ with rabbit anti-GFP antibodies (for PfSkp1GFP) or anti-HA antibodies (for PfCullin-2/cDD_HA_), followed by appropriate secondary antibodies as has been described in the expression and localization of proteins section.

### Effect of PfCullin-2 knock-down on parasite development

PfCull2KD_3D7_ parasites were maintained in the presence of 10 μM trimethoprim (+TMP) as has been described in the parasite culture section unless stated otherwise. To check knock-down of PfCullin-2/cDD_HA_ protein, a synchronized culture of PfCull2KD_3D7_ parasites with ∼10% rings was divided into two halves. One half was grown with (+TMP) and the other was grown without (-TMP) 10 μM trimethoprim for three consecutive cycles. Equal aliquots of both the cultures were harvested at the trophozoite stage (at ∼30 hr) in each cycle, and the parasites were assessed for PfCullin-2/cDD_HA_ protein level by western blotting using mouse anti-HA antibodies, followed by HRP-conjugated goat anti-mouse IgG as described in the expression and localization of proteins section. The blot was stripped and re-probed for β-actin as a loading control using mouse anti-β-actin-HRP antibodies.

To assess the effect of PfCullin-2 knock-down on asexual erythrocytic growth, synchronized PfCull2KD_3D7_ parasites were cultured with or without 10 μM TMP in 100 μl volume in a 96-well tissue culture plate, beginning with 4% ring stage parasitemia, for four consecutive cycles. If required, the cultures were diluted 2-4 folds at the end of the 48 hr development cycle and supplemented with fresh medium and RBCs to maintain 3-4% parasitemia. At the end of each cycle, a 50 μl aliquot of each culture was fixed (2% formaldehyde in PBS), stained with YOYO-1 (10 nM YOYO-1 in PBS with 0.1% Triton X-100), and the number of infected cells was determined by counting 10,000 cells using the BD Fortesa FACS Analyzer (λ_ex_ 491 nm and λ_em_ 509 nm). As a control, 100 μl suspension of the day-0 culture was also processed for FACS. The parasitemia at the end of each cycle was adjusted for dilution factor, normalized to day-0 parasitemia, and plotted as fold-multiplication over growth cycles using GraphPad Prism. In parallel, a 50 μl aliquot of each culture at the end of each cycle was processed for Giemsa staining, and assessed for parasite morphology using 100× objective of a light microscope (Axio-Imager Z2).

To evaluate the effect of PfCullin2 knock-down on sexual erythrocytic stages, PfCul2KD_3D7_ gametocytes were obtained by following the “crash method” [91]. Briefly, a synchronized PfCul2KD_3D7_ culture with 5% rings at 4% haematocrit was cultured in the presence of 10 μM TMP until 8-10% parasitemia with daily media change. The culture was divided into two equal aliquots, one was supplemented with 50 mM GlcNAc and 10 μM TMP and the other was supplemented with 50 mM GlcNAc only. Both the cultures were grown for 4 days with daily change of media and the supplements. On day 5 onward, the cultures were maintained with or without TMP, Giemsa-stained smears were made daily, and observed for gametocytes/2000 cells. The number of gametocytes was plotted over days using GraphPad Prism software. On day 4, a 20 ml aliquot of each culture was assessed for PfCullin-2/cDD_HA_ protein level by western blotting using mouse anti-HA antibodies, followed by HRP-conjugated goat anti-mouse IgG as described in the expression and localization of proteins section. The blot was stripped and re-probed for β-actin as a loading control with anti-β-actin-HRP antibodies.

### Electron microscopy

PfCul2KD_3D7_ parasites were evaluated for morphology and integrity of membraned structures by scanning electron microscopy (SEM) and transmission electron microscopy (TEM). Parasites were processed for SEM as has been reported earlier [92]. Briefly, synchronised PfCul2KD_3D7_ parasites were grown with (+TMP) or without (-TMP) 10 μM trimethoprim until the 3^rd^ cycle late trophozoite/schizont stage, and 10 ml culture with ∼10 % parasitemia was processed for saponin lysis to purify the parasites. The purified parasites were washed with 500 μl of RPMI 1640, resuspended in 50 μl RPMI 1640, and mixed with 50 μl fixative (2.5% paraformaldehyde, 5% glutaraldehyde, 0.2 M sodium cacodylate buffer, pH 7.4), and incubated for 1 hr at room temperature with slow mixing. The cells were washed twice with sodium cacodylate buffer and gradually dehydrated by incubating with increasing concentration of ethanol (50%, 70%, 95%, and 100%), each incubation was for 3 minutes with fresh solvent. The cells were resuspended in 100 μl water, an aliquot was diluted 100-fold with water and dried on a glass coverslip at room temperature. The coverslip was mounted on the stub and gold spurring was done using the Quorum 150T ES plus. The cells were observed in scanning electron microscope (Hitachi science systems) at 3-5 μm resolution.

For TEM, synchronised PfCul2KD_3D7_ parasites were grown with (+TMP) or without (-TMP) 10 μM TMP until the 3^rd^ cycle late trophozoite/schizont stages, and the parasites were fixed as has been described above for SEM. The fixed cells were washed three times with 0.1 M cacodylate buffer (pH 7.4), and incubated in 1% osmium tetroxide for 30 min to 1 hr at room temperature. The cells were washed twice with cacodylate buffer, stained *en bloc* with 2% uranyl acetate for 20 min, washed twice with the cacodylate buffer, and gradually dehydrated with increasing concentration of ethanol (50%, 70%, 95%, and 100%), each incubation was for 3 min with fresh solvent. The dehydrated sample was incubated and washed twice in propylene oxide for 5 min each. The sample was incubated with a 1:1 and 3:1 mixture of Spurr’s epoxy resin and propylene oxide for 15 and 20 min, respectively, followed by a final wash with pure epoxy resin. Finally, the cells were embedded in epoxy resin with polymerising agent and kept at 62°C for 12-14 hr. The sample was trimmed and sectioned with a glass knife using the ultramicrotome (LEICA EM UC7), and placed on a copper grid. The sample grids were post-stained with uranyl acetate for 1 hr, followed by with lead citrate for 2 min with intermittent washing with water. The sample grids were dried, loaded on the transmission electron microscope (Talos L120C), images were captured with CETA camera at 500 nm, and analysed by Acquisition-Velox software.

### Label free quantification of proteins

Synchronized PfCull2KD_3D7_ parasites were cultured with (+TMP) or without (-TMP) 10 μM trimethoprim until the 3^rd^ cycle late trophozoite/early schizont stage, and parasites were purified by saponin lysis method. The parasite pellets were resuspended in lysis buffer (100 mM Tris, 4% SDS, 200 mM DTT, 1% Triton X-100, pH 7.5), centrifuged, and the supernatant protein content was estimated by amido black assay. 150 μg of the +TMP or -TMP parasite lysate was mixed with 0.02% bromophenol blue (with 10% glycerol), and ran on 12% SDS-PAGE gel. The gel was stained and each protein lane was cut into 8 slices, and each slice was processed for mass spectrometry as has been described in the mass spectrometry section. The MS data of each of the 8 slices of +TMP and -TMP samples were run separately against the *P. falciparum* Uniprot database, and abundance values were obtained using the label free quantification tool of Sequest HT. The abundance values from two biological replicates were analysed to obtain log2-fold change of protein levels in -TMP sample as compared to the +TMP sample, and *P*-values for significance of the difference between +TMP and -TMP data sets were calculated using the Student’s T-test. A log2-fold change of ≥+1 was considered as upregulated, ≤-1 was considered as downregulated, and that between +1 to -1 was considered as unchanged. Upregulated and downregulated proteins with *P*-values ≤0.05 were further analysed for differential regulation using the VolcanoseR online tool, and pathway enrichment was obtained by gene set enrichment analysis (GSEA) with the programme R.

### Assessment of ER stress, DNA damage and mitochondrial membrane potential

ER stress was measured as has been described earlier [93]. Briefly, PfCul2KD_3D7_ parasites were grown with (+TMP) or without (-TMP) 10 μM trimethoprim for 3 cycles. Equal number of 3^rd^ cycle parasites of both the cultures were incubated with different concentrations of DTT (5, 10, and 20 mM) or equal amount of water for 1 hr, followed with 10 μM of the peroxide sensitive dye 2’,7’-dichlorodihydrofluorescein diacetate acetyl ester (H_2_DCFDA) for 30 min in the dark with gentle shaking at room temperature. The fluorescence was measured using the SpectraMax iD3 multimode plate reader (λ_em_ = 530 nm and λ_ex_ = 488 nm).

For measuring the effect of DNA-damaging agent, synchronised PfCul2KD_3D7_ parasites were grown with (+TMP) or without (-TMP) 10 μM trimethoprim untill the 3^rd^ cycle mid trophozoite stage, followed by in the presence of 0.005% DMSO or 0.005% MMS for 6 hr. The cultures were washed with cultured medium, supplemented with fresh medium, grown further for 12 hrs, Giemsa stained smears were made, and about 1000 cells were observed for fresh rings.

Mitochondrial stress was measured as membrane potential by treating parasites with JC-1, a cationic mitochondrial membrane potential dye. It forms red aggregates in healthy/polarised mitochondria and remains as a green monomer if mitochondria are unhealthy. The ratio of red/green fluorescence indicates the health status of mitochondria. Asynchronous cultures of PfCul2KD_3D7_ parasites were grown with (+TMP) or without (-TMP) 10 μM trimethoprim for 3 cycles, and equal number of the 3^rd^ cycle parasites were incubated in 2 µM JC-1 (in RPMI 1640) for 20 min at 37°C. The parasites were washed with culture medium, and fluorescence was measured using SpectraMax iD3 multimode plate reader (red: λ_em_ = 590 nm and λ_ex_ = 514 nm; green: λ_em_ = 529 nm and λ_ex_ = 514 nm).

### Assessment of DNA replication and mitotic structures

PfCul2KD_3D7_ parasites were evaluated for DNA replication by estimating the total DNA content and the number of nuclei. Synchronized PfCul2KD_3D7_ parasites were grown with (+TMP) or without (-TMP) 10 μM trimethoprim for 3 cycles, beginning with equal parasitemia. Equal aliquots of the cultures were taken at 10, 36, and 48 hrs during the first 2 cycles and at 10, 28, 32, 36, 40, and 44 hrs during the 3^rd^ cycle. Half of the each aliquot was fixed (4% paraformaldehyde in PBS) for 45 min at room temperature, treated with 0.1% Triton X-100 (in PBS) to permeabilize the cells, treated with RNase A (0.3 mg/ml) for 20 min at 37°C to remove RNA, and washed with PBS. The samples were stained with SYBR green I (1:2000 v/v in PBS) and the fluorescence of each sample was measured using SpectraMax iD3 multimode plate reader (λem = 530 nm and λex = 488 nm). For microscopy, parasites from the other half of the 3^rd^ cycle aliquots were smeared on glass slides, fixed (3% paraformaldehyde and 0.01% glutaraldehyde in PBS), permeabilized with 0.1% Triton X-100 (v/v in PBS), stained with DAPI, observed under 100× objective of ZEISS Axioimager, and images were captured with Zeiss AxioCam HRm camera. The number of nuclei/cell were counted for each time point.

For the analysis of mitotic structures (spindle fibres and centriolar plaques), parasites were collected at 40 hr stages during the 3^rd^ cycle, and processed for IFA as has been described in the expression and localization section. Briefly, for spindle fibres, parasites were stained with alexa-488-conjugated mouse anti-α-tubulin antibodies (at 1/1000 dilution in blocking buffer) and DAPI (10 μg/ml). For centriolar plaques, parasites were stained with rabbit anti-centrin-1 antibodies (at 1/250 dilution in blocking buffer), followed by with alexa-488-conjugated goat anti-rabbit IgG (at 1/1000 in blocking buffer) and DAPI (10 μg/ml). The slides were observed under 100× objective of ZEISS Axioimager and images were captured with Zeiss AxioCam HRm camera. About 100 parasites were observed for staining patterns.

### Statistical analysis

The data were analyzed to determine significance of difference (*P*-value) between two groups using the t-test. Statistical significance is indicated for data sets showing significant *P*-values (* is p<0.05-0.01, ** is p<0.01-0.001, *** is p<0.001).

### Data availability

The mass spectrometry proteomics data have been deposited to the ProteomeXchange Consortium via the PRIDE partner repository with the dataset identifiers PXD040195 (PfCullin-1), PXD040090 (PfRbx1), PXD040194 (PfSkp1), PXD040268 (PfCullin-2) and PXD040197 (LFQ).

## Acknowledgements

We thank the staff of CCMB’s Advanced Microscopy and Imaging Facility, Proteomics Facility, and CryoEM facility for assistance with experiments related to imaging, mass spectrometry, and EM imaging, respectively.

## Funding information

This study was supported by grants from the Department of Biotechnology, India (BT/PR11497/MED/29/854/2014) and Council of Scientific and Industrial Research, India (MLP0148) to PS. ZR, GSR and SG the recipients of fellowships from the Department of Biotechnology, India, Council of Scientific and Industrial Research, India, and University Grants Commission, India, respectively. PP was supported with funds from Council of Scientific and Industrial Research, India (MLP0148). DD was supported with funds from Department of Biotechnology, India (BT/PR11497/MED/29/854/2014).

## Conflict of interest

The authors do not have any conflict of interest with the contents of this article.

## Supplementary Information

### Supplementary figures and figure legends

**Figure S1.**
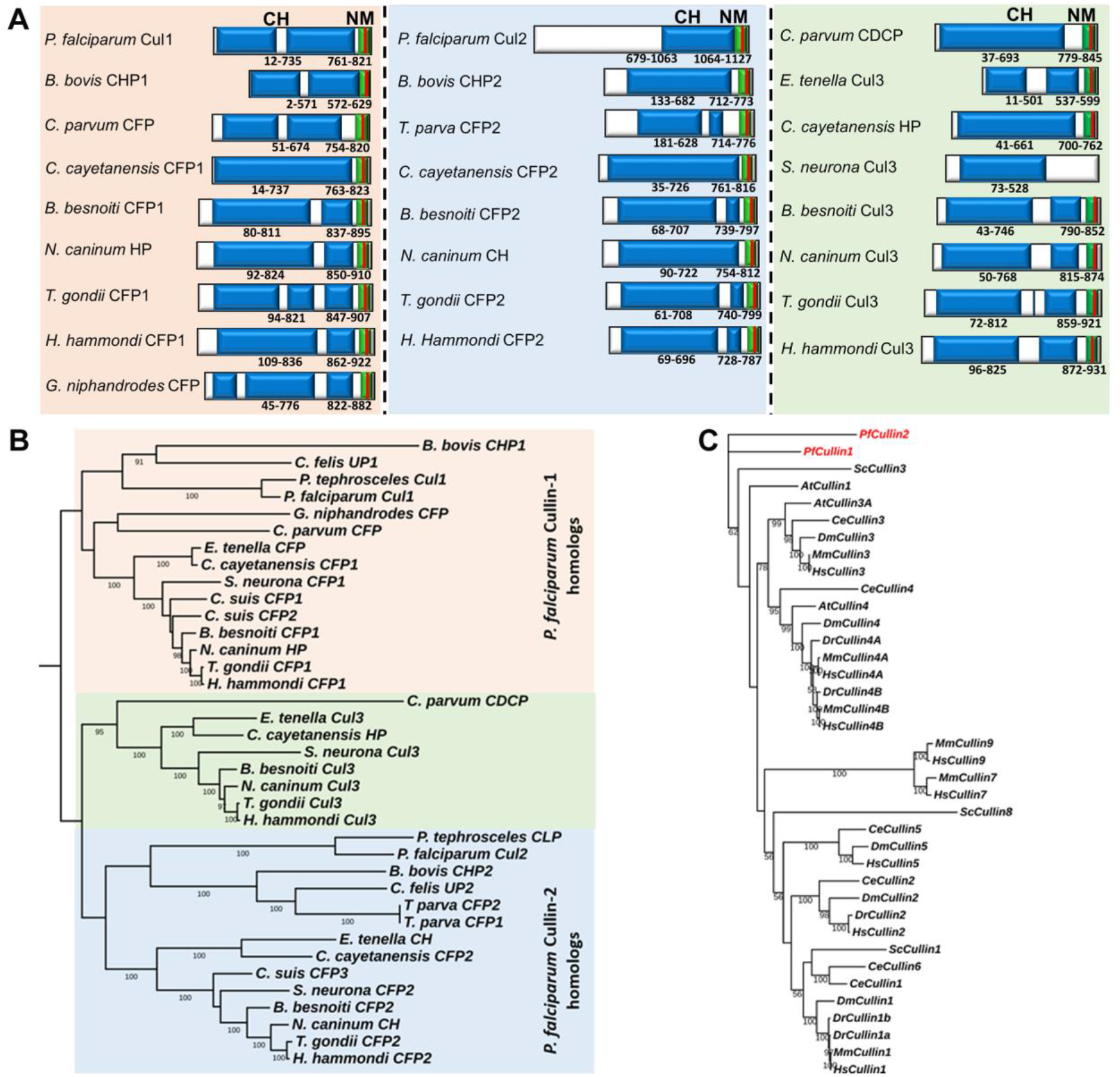
Domain organisation and phylogenetic relatedness of Apicomplexan cullin homologs. The amino acid sequences of Apicomplexan cullin homologs were analysed for conserved domains and phylogenetic relatedness within Apicomplexa and with homologs in model organisms. **A.** The schematic represents domain organization of the indicated cullins with the boundaries of cullin homology domain (CH) and neddylation motif (NM). The red bar in NM is the conserved putative lysine residue that is conjugated to NEDD8. **B.** The phylogram shows that PfCulin-1 (in pink) or PfCulin-2 (in blue) homologs of the indicated Apicomplexan parasites cluster together on separate nodes. The cullins (in green) represent the third cullin homolog in the indicated Apicomplexan parasites. The annotations are: CHP, conserved hypothetical protein; UP, unspecified product; CFP, cullin-family protein; HP, hypothetical protein; CDCP, cullin domain-containing protein; CLP, cullin like protein; CH, cullin homology domain; Cul, cullin. **C.** The phylogram of PfCullins and their homologs in model organisms shows that PfCullin-1 and PfCullin-2 are distantly related to their homologs in the indicated model organisms. The number on branches denote the strength of clustering (bootstrap value).

**Figure S2:**
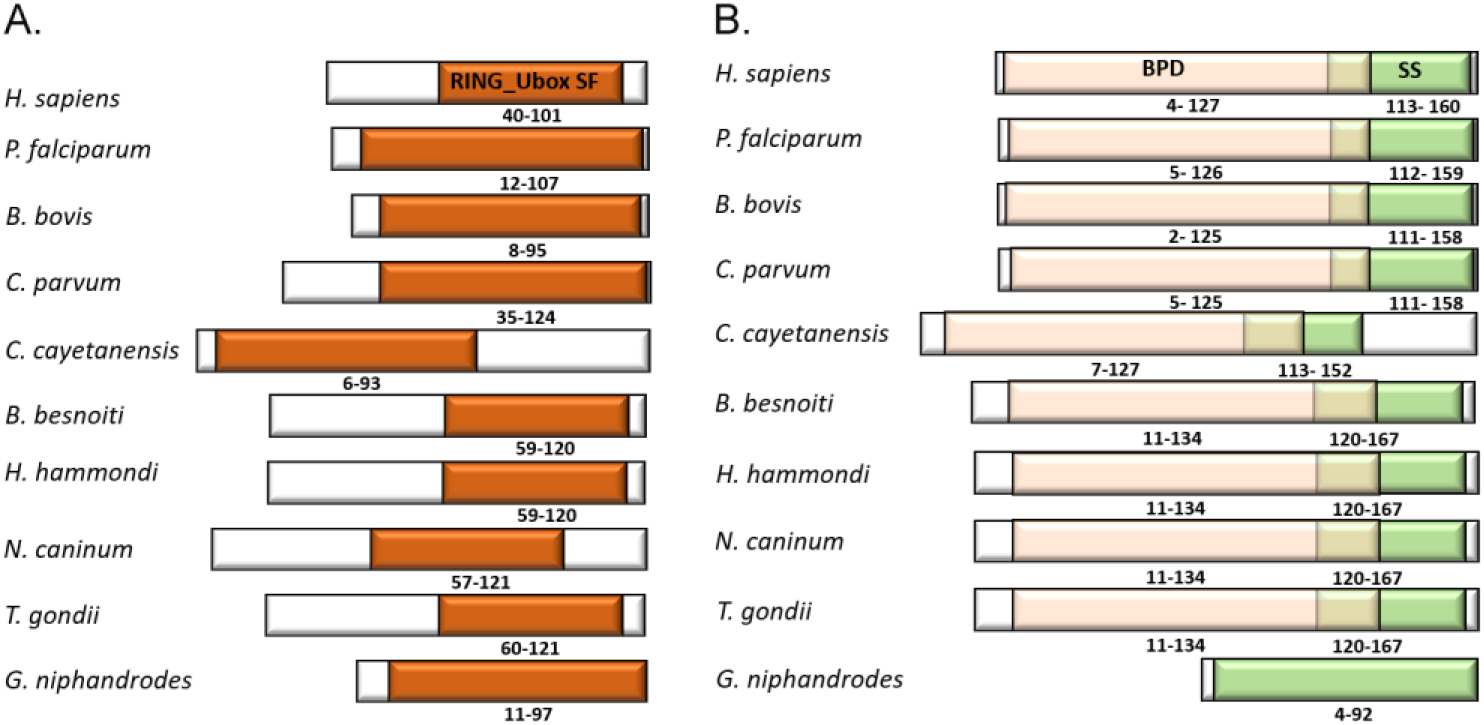
Domain organisation of Rbx1 and Skp1 homologs. The amino acid sequences of putative Apicomplexan Rbx1 and Skp1 were analysed for conserved domains and phylogenetic relatedness within the Apicomplexa. **A.** The schematic represents domain organization of the indicated Rbx1 protein with the boundary of RING_Ubox superfamily (RING_Ubox SF) domain, which contains the consensus zinc finger motif [C-X2-C-X(9-39)-C-X(1-3)-H-X(2-3)-(N/C/H)-X2-C-X(4-48)-C-X2-C]. **B.** The schematic represents domain organization of the indicated Skp1 protein with the boundaries of the BTB (Broad-Complex, Tramtrack and Bric a brac)/POZ (poxvirus and zinc finger) domain (BPD), Skp1 superfamily (SS) domain, and the sequence shared by BPD and SS regions.

**Figure S3.**
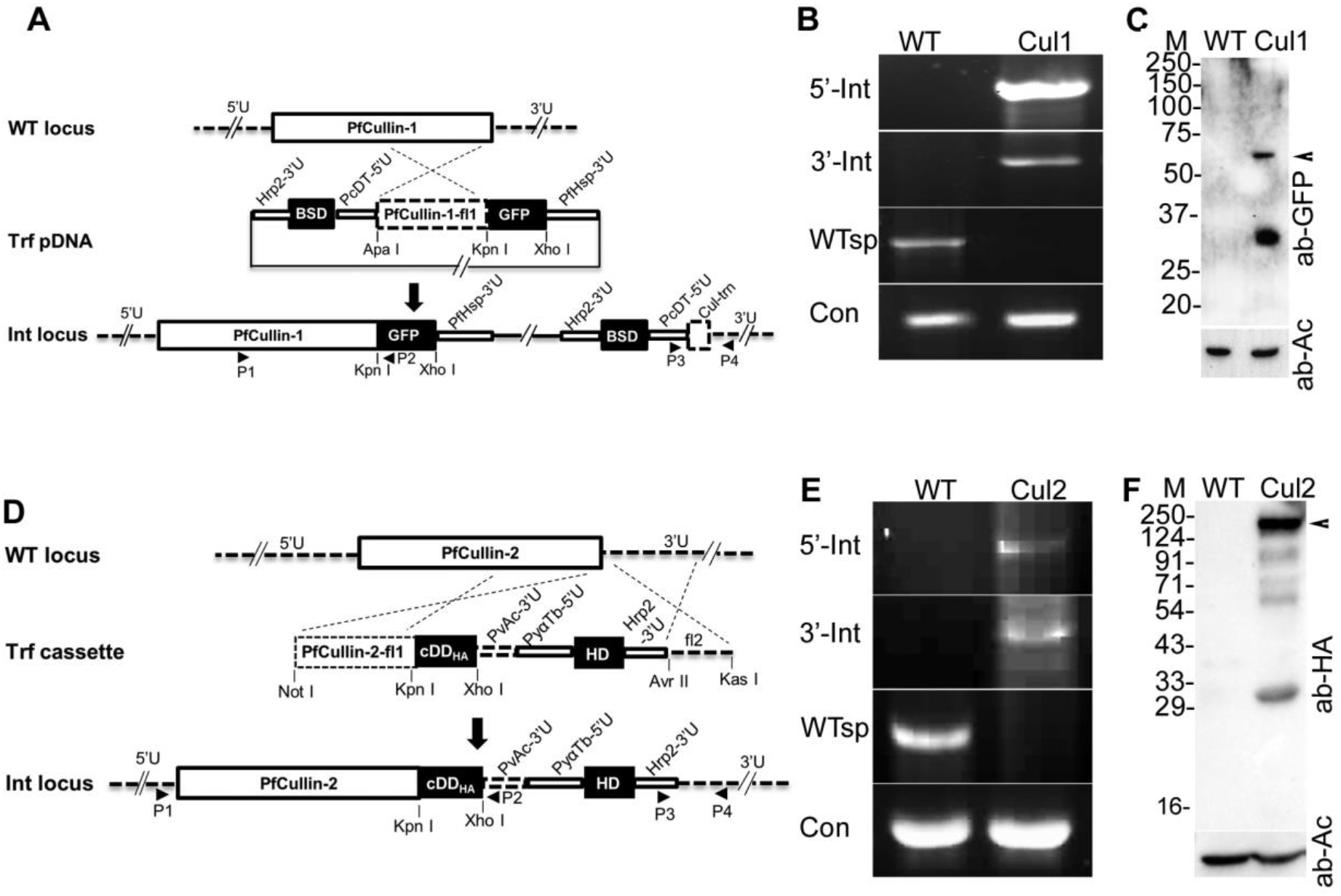
Generation of PfCul1GFPKI and PfCul2KDD10 parasites. The endogenous PfCullin-1 and PfCullin-2 coding regions were replaced with PfCullin-1/GFP and PfCullin-2/cDD_HA_ coding sequences in *P. falciparum* D10 strain, respectively. Cloned recombinant lines were assessed for integration of the desired DNA into the target locus by PCR. **A.** The schematic represents integration of transfection plasmid DNA (Trf pDNA) into the endogenous PfCullin-1 locus (WT locus), generating the integration locus (Int locus). Rectangular boxes represent the coding regions for PfCullin-1, PfCullin-1-fl1, GFP and BSD. The untranslated regions of PfCullin-1 (5ʹU and 3ʹU), the location and orientation of primers (horizontal arrows), the restriction endonuclease sites (vertical lines), and regulatory regions in the linear transfection construct (PfHsp86-3ʹU, PcDT-5ʹU and PfHrp2-3ʹU) are indicated. **B.** The ethidium bromide-stained agarose gel shows PCR products amplified from the genomic DNA of wild type (WT) and PfCul1GFPKI (Cul1) parasites using locus specific primers (P1 and P2 for 5’ integration (5’-Int), P3 and P4 for 3’ integration (3’-Int), P1 and P4 for wild type (WTsp), and primers specific for the PfRbx1 gene were used as a positive control (Con)). **C.** The lysates of wild type *P. falciparum* D10 (WT) and PfCul1GFPKI (Cul1) parasites were assessed for expression of PfCullin-1/GFP by western blotting using anti-GFP (ab-GFP) and anti-β-actin (ab-Ac) antibodies. The arrow indicates the size of truncated PfCullin-1/GFP, and sizes of protein markers (M) are in kDa. **D.** The schematic represents integration of linear transfection construct (Trf cassette) into the endogenous PfCullin-2 locus (WT locus), resulting into the generation of integration locus (Int locus). Rectangular boxes represent the coding regions for PfCullin-2, PfCullin-2-fl1, *E. coli* mutant DHFR with HA-tag (cDD_HA_) and human DHFR (HD). The flanking untranslated regions (5ʹU and 3ʹU), the location and orientation of primers (horizontal arrows), the restriction endonuclease sites (vertical lines) and regulatory regions in the linear transfection construct (PfCullin-2-flank2 (fl2), PvAc-3ʹU, PyαTb-5ʹU, and PfHrp2-3ʹU (Hrp2-3ʹU)) are indicated. Locus specific primers (P1 and P2 for 5’ integration (5’-Int), P3 and P4 for 3’ integration (3’-Int), P1 and P4 for wild type (WTsp), and primers specific for the PfAtg18 gene were used as a positive control (Con)) were used in PCR to differentiate integration and wild type loci. **E.** The ethidium bromide-stained agarose gel shows PCR products amplified from the wild type (WT) and PfCul2KDD10 (Cul2) genomic DNA using the indicated primer sets. **F.** The lysates of wild type *P. falciparum* D10 (WT) and PfCul2KDD10 (Cul2) parasites were assessed for expression of PfCullin-2/cDD_HA_ by western blotting using anti-HA (ab-HA) and anti-β-actin (ab-Ac) antibodies. The arrow indicates the size of full-length PfCullin-2/cDD_HA_. The sizes of protein markers (M) are in kDa.

**Figure S4.**
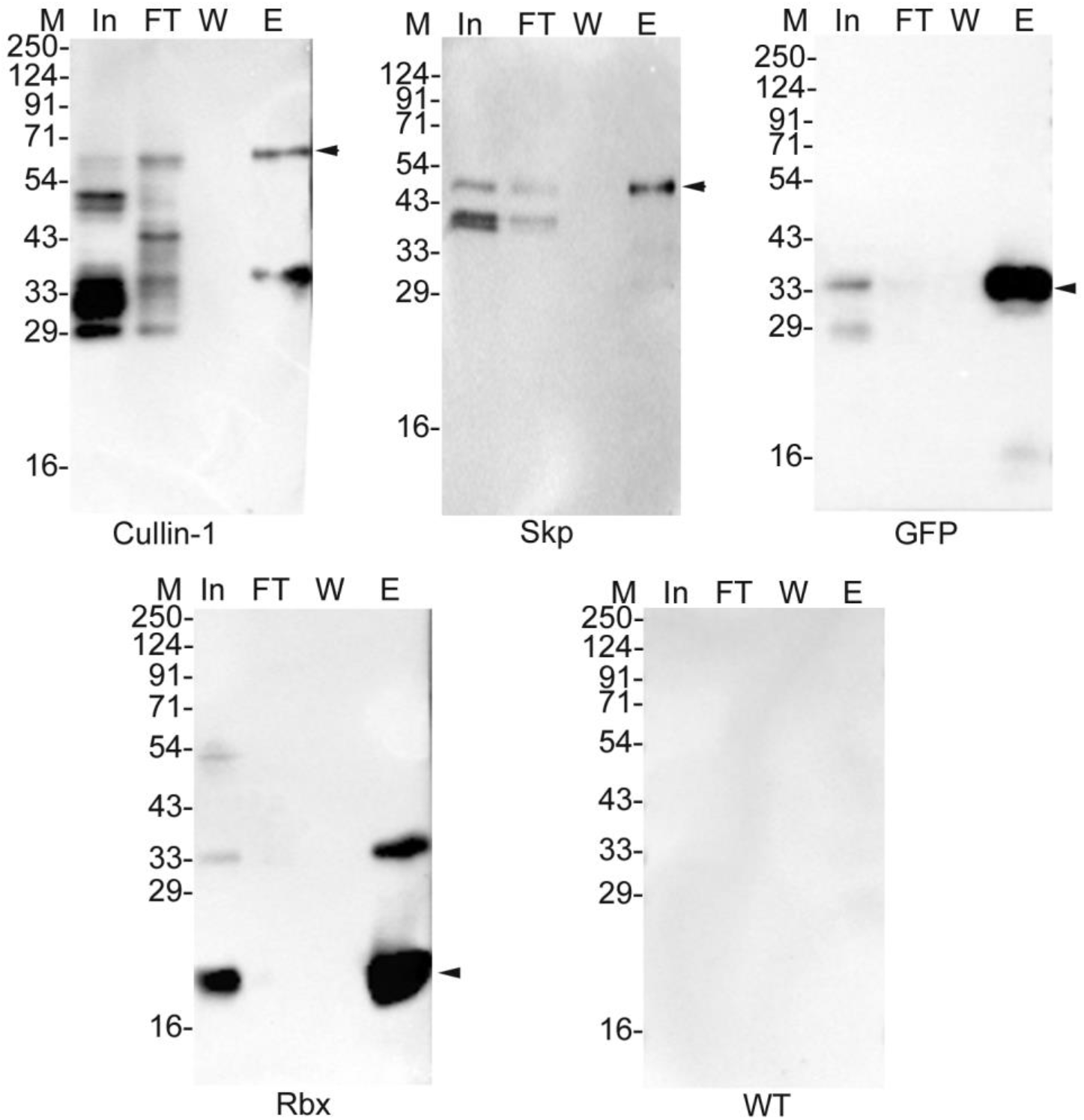
Immunoprecipitation. Lysates of PfCul1GFPKI (Cullin-1) PfSkp1/GFP-epi (Skp), and GFP-expressing control (GFP) parasites were processed for immunoprecipitation using GFP-Trap antibodies. Lysates of PfRbx1myc-epi (Rbx) and wild type *P. falciparum* D10 (WT) parasites were processed for immunoprecipitation using Myc-Trap antibodies. Aliquots of the input lysate (In), flow through (FT), wash (W), and eluate (E) for all the immunoprecipitations were assessed for the presence of target proteins using mouse anti-Myc (for Rbx and WT) or rabbit anti-GFP (for Cullin-1, Skp, GFP) antibodies by western blotting. Arrow heads indicate target proteins, and sizes of protein markers (M) are in kDa.

**Figure S5.**
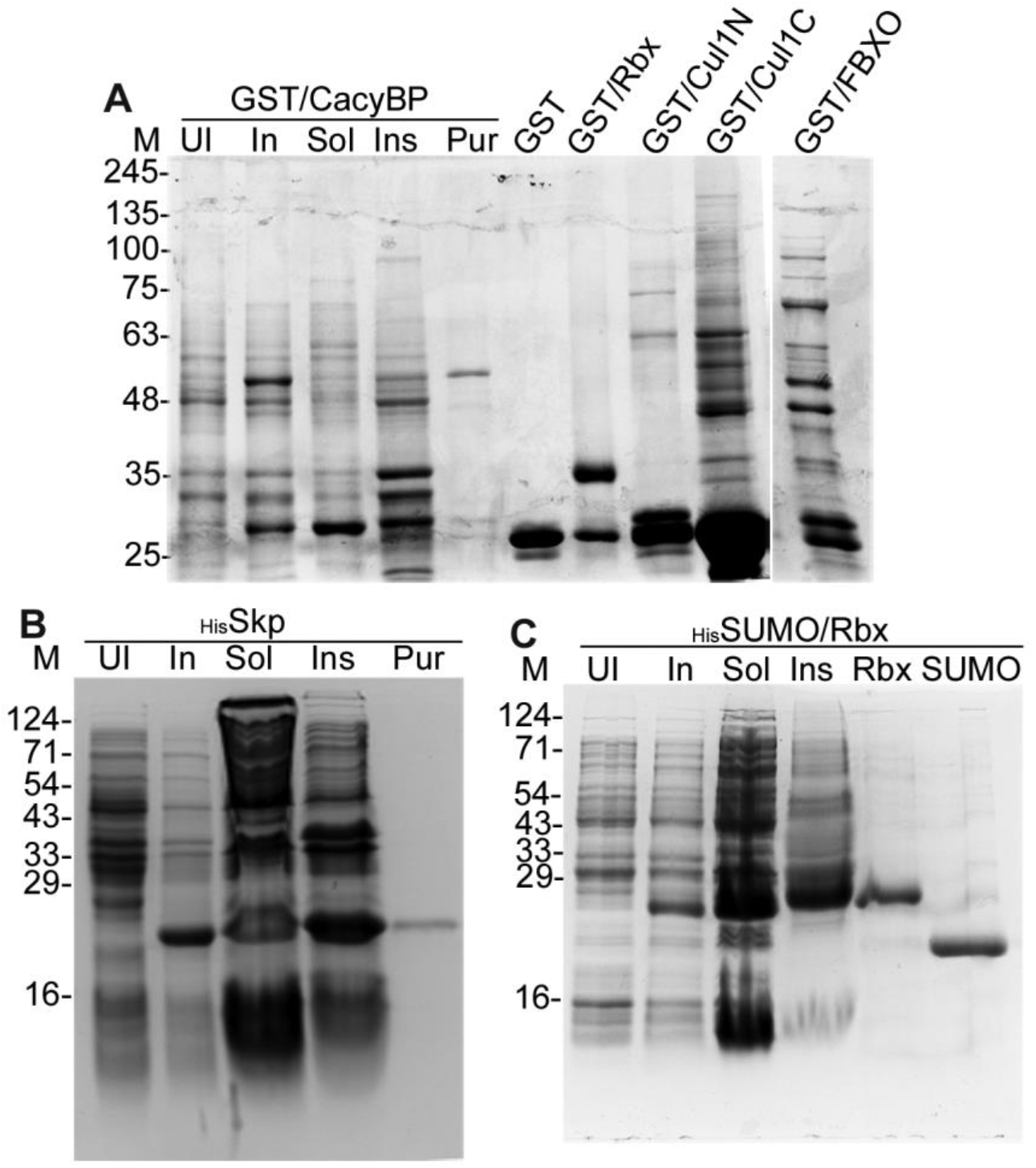
Production of recombinant PfSCF subunits. **A.** GST alone or GST-tagged CacyBP (GST/CacyBP), PfRbx1 (GST/Rbx), PfCullin-1 N-terminal (GST/Cul1N) and C-terminal (GST/Cul1C) fragments, and PfFBXO1 (GST/FBXO) were expressed in *E. coli*. The coomassie stained SDS-PAGE gel contains total lysates of un-induced (UI) and IPTG-induced (In) cells, soluble (Sol) and insoluble (Ins) fractions of the induced cells, and purified GST/CacyBP. The lanes GST, GST/Rbx, GST/Cul1N, GST/Cul1C and GST/FBXO contain the respective purified or enriched proteins. **B.** 6×His-tagged PfSkp1 (HisSkp) was expressed in *E. coli* cells. The coomassie stained SDS-PAGE gel contains total lysates of un-induced (UI) and IPTG-induced (In) cells, soluble (Sol) and insoluble (Ins) fractions of the induced cells, and purified protein (Pur). **C.** 6×His-SUMO-tagged PfRbx1 (HisSUMO/Rbx) or 6×His-SUMO (HisSUMO) proteins were expressed in *E. coli*. The coomassie stained SDS-PAGE gel contains total lysates of un-induced (UI) and IPTG-induced (In) cells, soluble (Sol) and insoluble (Ins) fractions of the induced cells, and purified HisSUMO/Rbx (Rbx) and HisSUMO (SUMO) proteins. The protein size markers (M) are in kDa.

**Fig S6.**
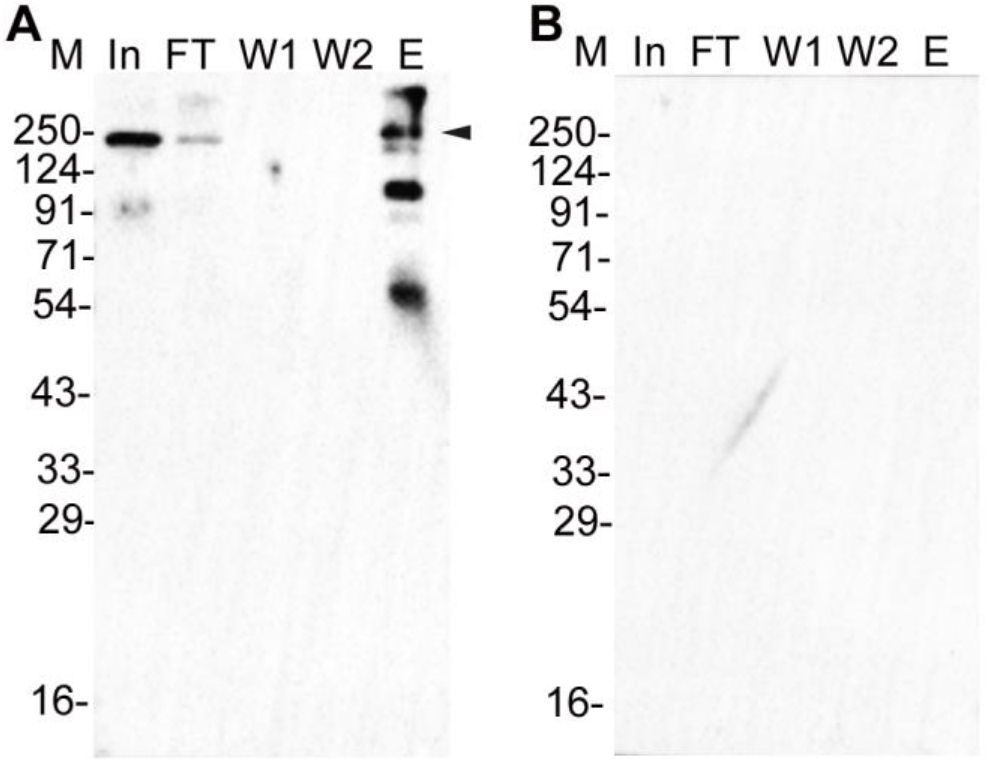
Immunoprecipitation of PfCullin-2. Lysates of PfCul2KDD10 (**A**) and wild type *P. falciparum* D10 (**B**) parasites were processed for immunoprecipitation using rabbit anti-HA antibodies. Aliquots of the parasite lysate inputs (In), flow through (FT), washes (W1, W2), and eluate (E) samples were probed with mouse anti-HA antibodies by western blotting. The sizes of protein markers (M) are in kDa.

**Fig S7.**
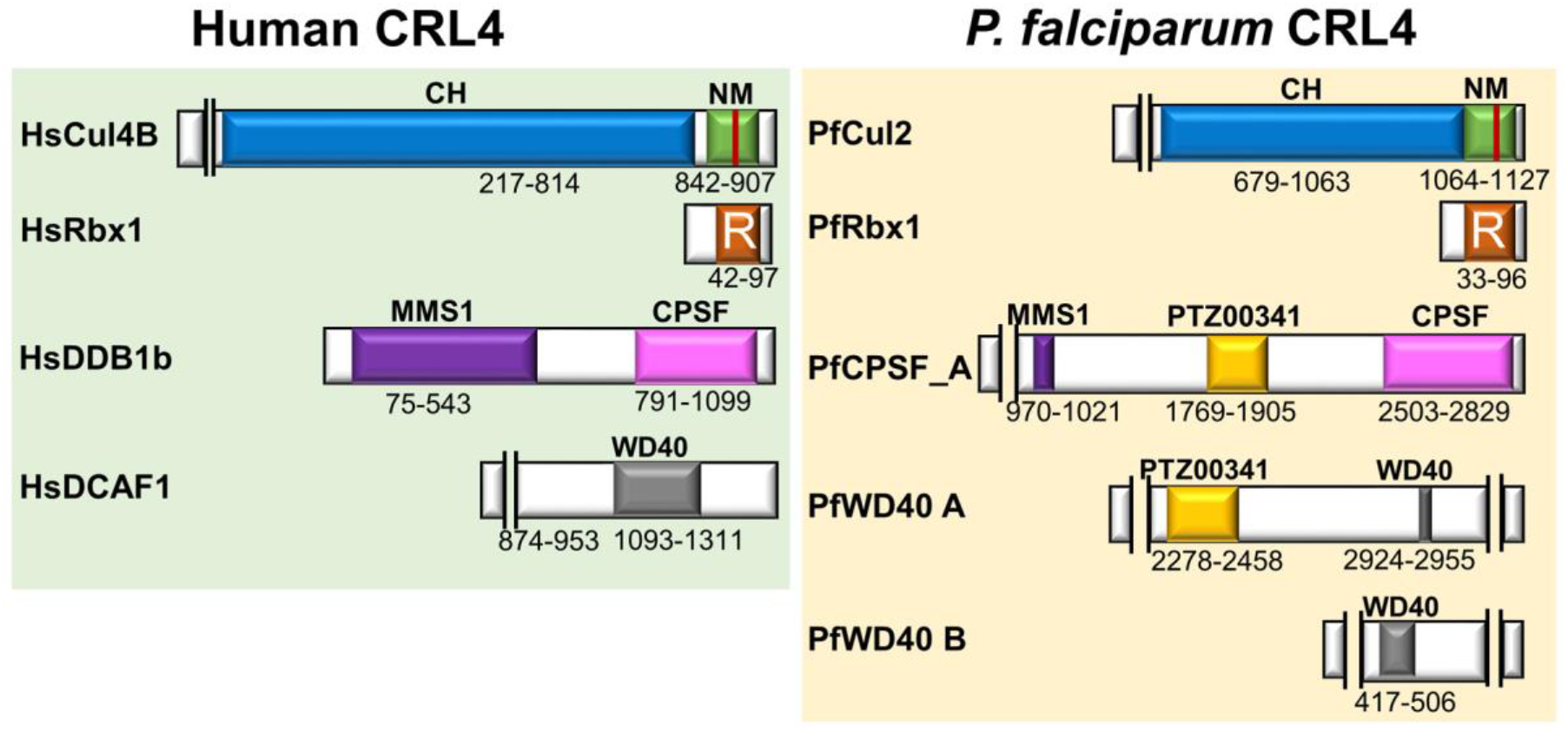
Domain organization of CRL4 subunits. The schematic represents predicted domain architecture of the core subunits of human CRL4B and *P. falciparum* CRL4. The human CRL4B contains Cullin-4B (HsCul4B), Rbx1 (HsRbx1), DNA damage-specific binding protein 1b (HsDDB1b) and DDB1-Cullin-4-associated factor 1 (HsDCAF1). The *P. falciparum* CRL4 complex appears to contain Cullin-2 (PfCul2), Rbx1 (PfRbx1), cleavage and polyadenylation specific factor subunit A (PfCPSF_A) and WD40 repeat proteins (PfWD40 A and PfWD40 B). The shaded boxes are conserved domains/motifs, which include cullin homology domain (CH) and neddylation motif (NM) in cullins, RING finger domain (R) in Rbx1, methyl methanesulfonate-sensitivity protein 1 (MMS1) and CPSF domains in HsDDB1 and PfCPSF_A proteins, WD40 repeat domain in HsDCAF1 and PfWD40 proteins. PfCPSF_A and PfWD40 A proteins also contain PTZ00341 domain of unknown function, which is present in ring-infected erythrocyte surface antigen (RESA) and several other *Plasmodium* proteins. The number below each domain corresponds to the number of boundary amino acid residues. The protein size is scaled to 1 cm for 100 aa; some regions are interrupted by two vertical line to fit the protein size.

**Fig S8.**
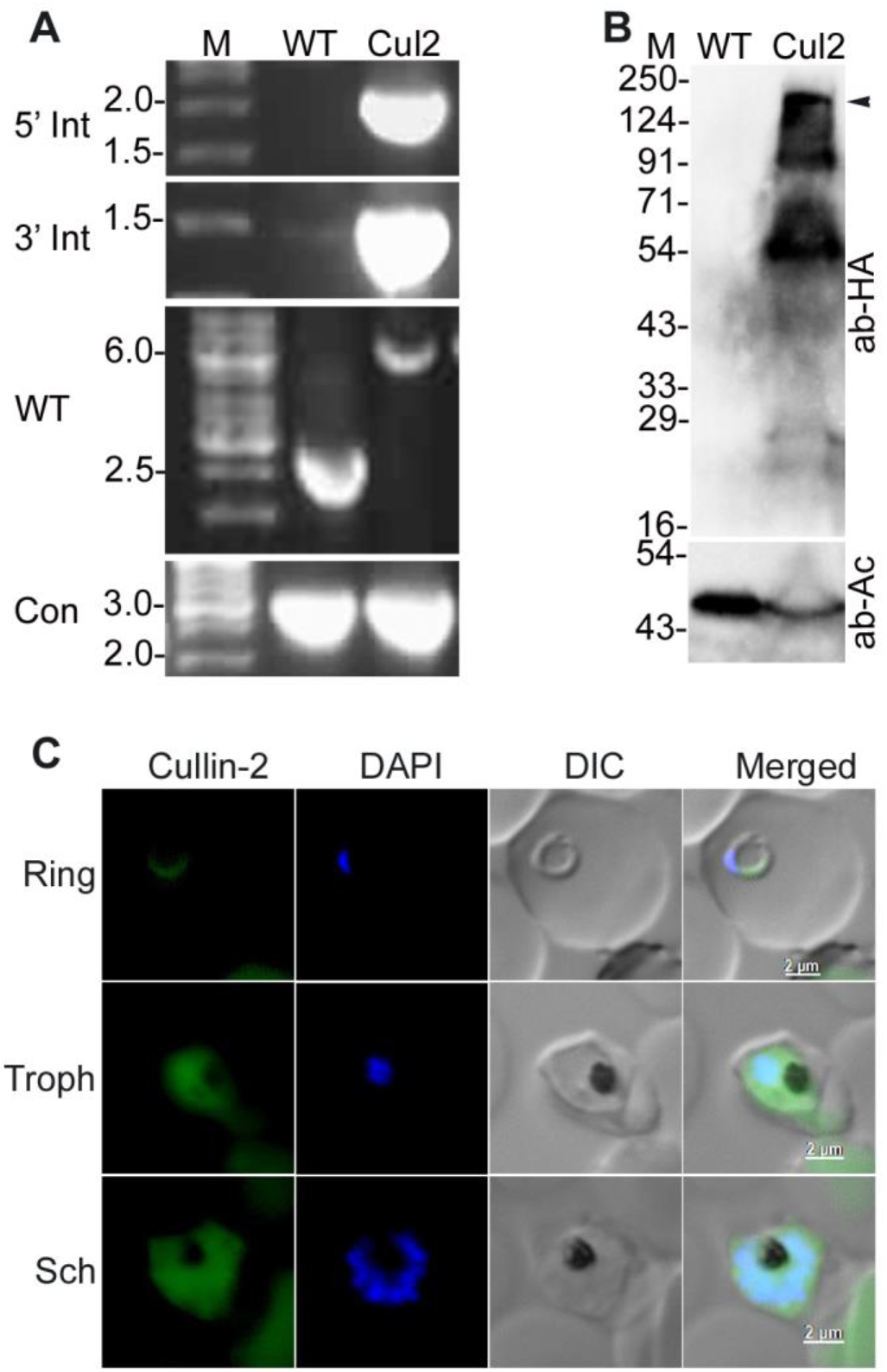
Generation of PfCul2KD_3D7_ parasites. The endogenous PfCullin-2 coding region was replaced with PfCullin-2/cDD_HA_ coding sequence as shown in Figure S3C. A cloned line was assessed for the presence of integration locus by PCR and expression of PfCullin-2/cDD_HA_ protein by western blotting. **A.** The ethidium bromide-stained agarose gel shows PCR products amplified from the wild type (WT) and PfCul2KD_3D7_ (Cul2) genomic DNA using the indicated primer sets. DNA size markers (M) are in kbp. **B.** The lysates of wild type *P. falciparum* 3D7 (WT) and PfCul2KD_3D7_ (Cul2) parasites were assessed for expression of PfCullin-2/cDD_HA_ by western blotting using anti-HA (ab-HA) and anti-β-actin (ab-Ac) antibodies. The arrow indicates the size of full-length PfCullin-2/cDD_HA_. **C.** Ring, trophozoite (Troph), and schizont (Sch) stages of PfCul2KD_3D7_ parasites were assessed for localization of PfCullin-2/cDD_HA_ by IFA using anti-HA antibodies. The images are for PfCullin-2/cDD_HA_ signal (Cullin-2), nuclear staining (DAPI), parasite and RBC boundaries (DIC), and overlap of all the three images (Merged). Scale bar is in the merged image.

**Figure S9.**
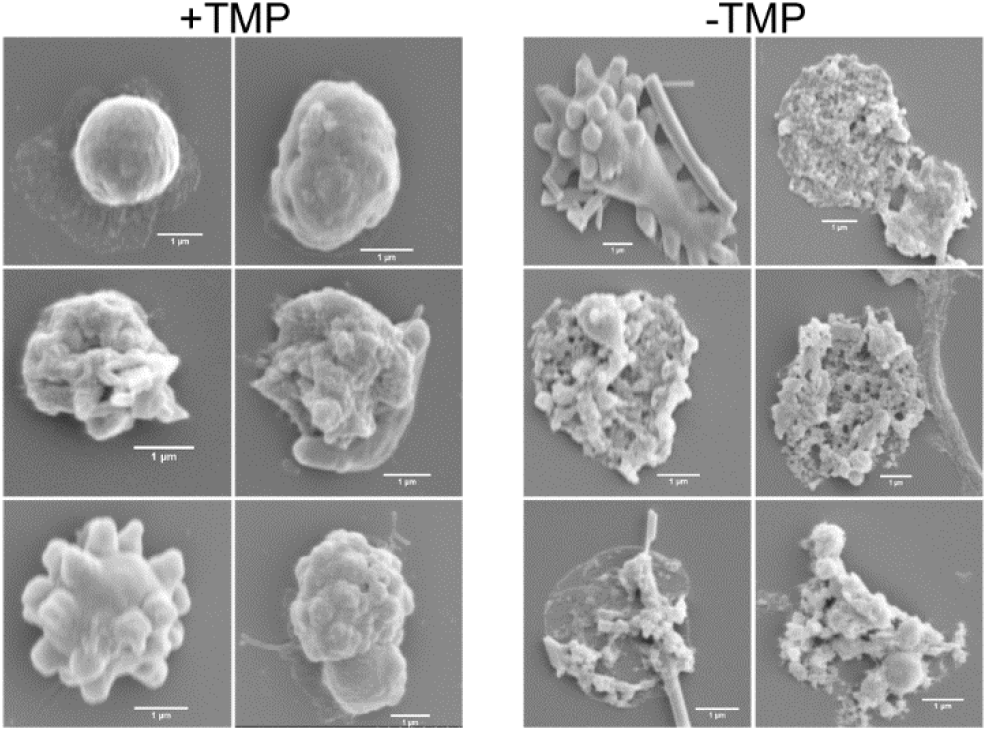
Scanning electron micrographs of PfCul2KD_3D7_ parasites. PfCul2KD_3D7_ parasites were cultured with (+TMP) or without (-TMP) for three cycles, the 3^rd^ cycle parasites were isolated by saponin lysis, and processed for SEM. The images show outer surface of the parasites, which appears smooth in case of +TMP, but distorted in case of -TMP parasites.

**Figure S10.**
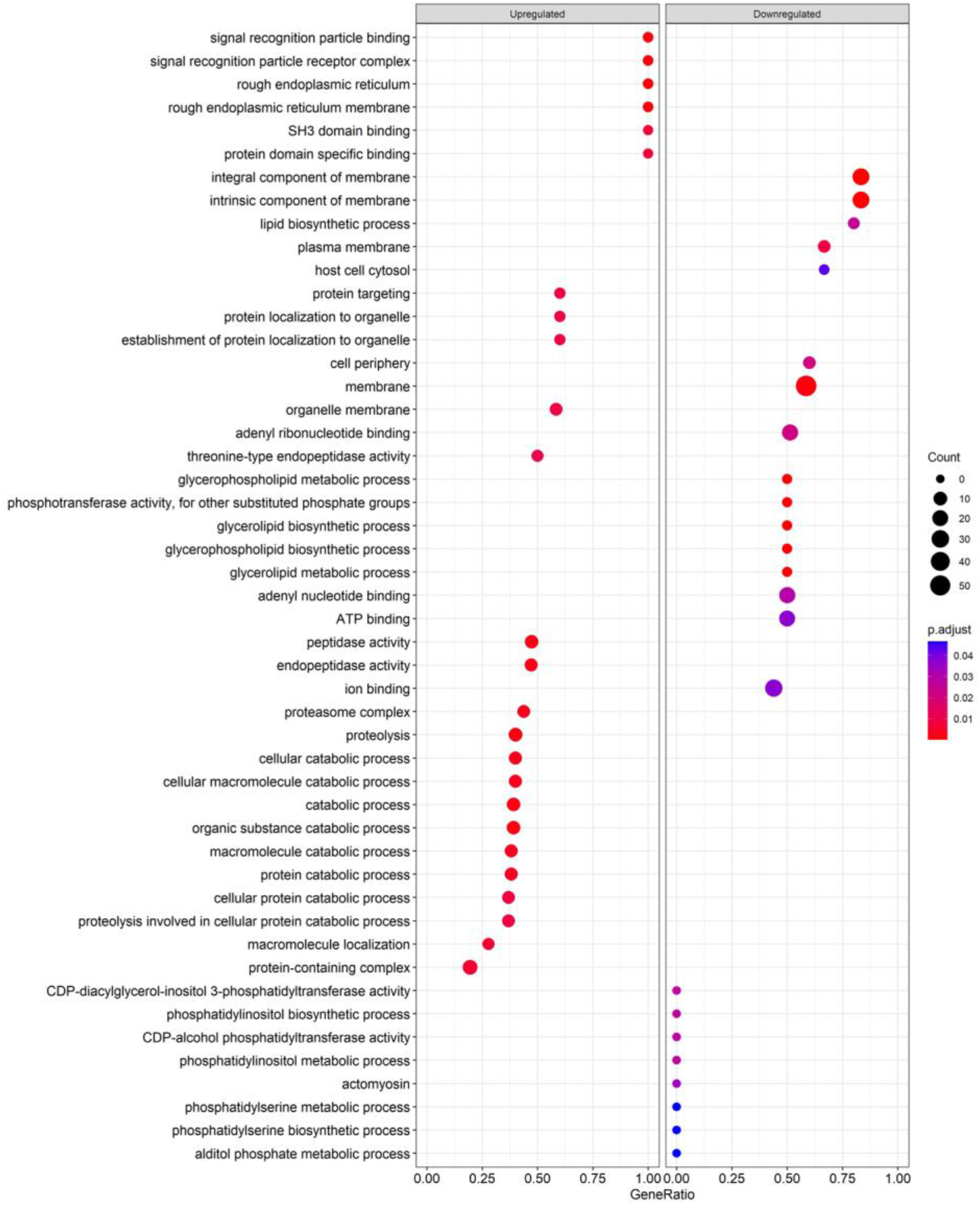
Pathway enrichment of differentially regulated proteins in PfCul2KD_3D7_ trophozoites. PfCul2KD_3D7_ parasites were cultured with or without trimethoprim for three cycles, and the 3^rd^ cycle trophozoites were processed for LFQ. Differentially regulated proteins were used for gene set enrichment analysis (GSEA) using the cluster profiler for gene ontology (GO) term. The plot shows biological pathways on Y-axis and gene enrichment values on X-axis. The size and color of the circle denotes the number of GO terms involved in a particular pathway and their confidence, respectively. Bigger the size and darker the red color of the circle corresponds to higher confidence and enrichment of the pathways.

### Supplementary tables

**Table S1.**
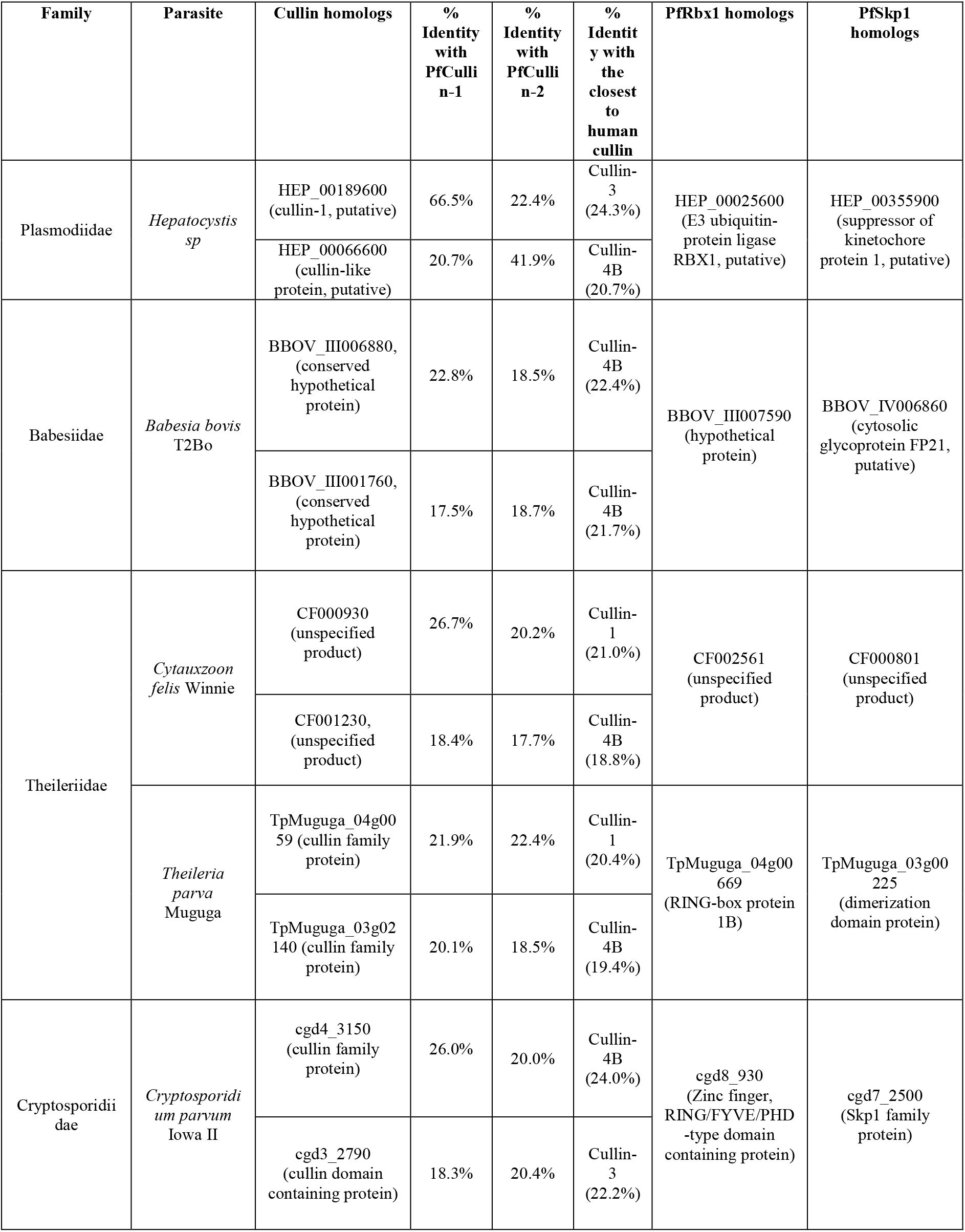
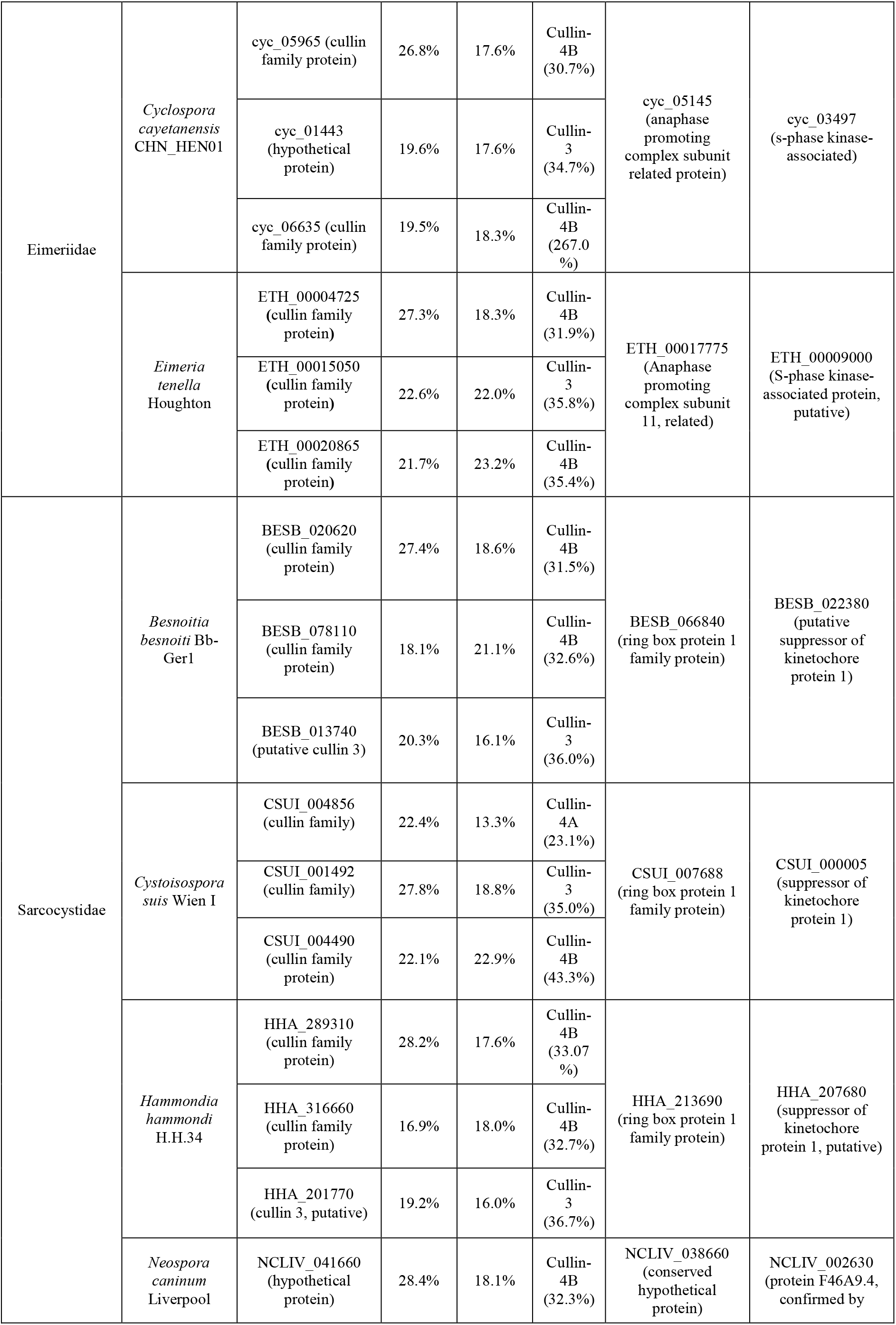
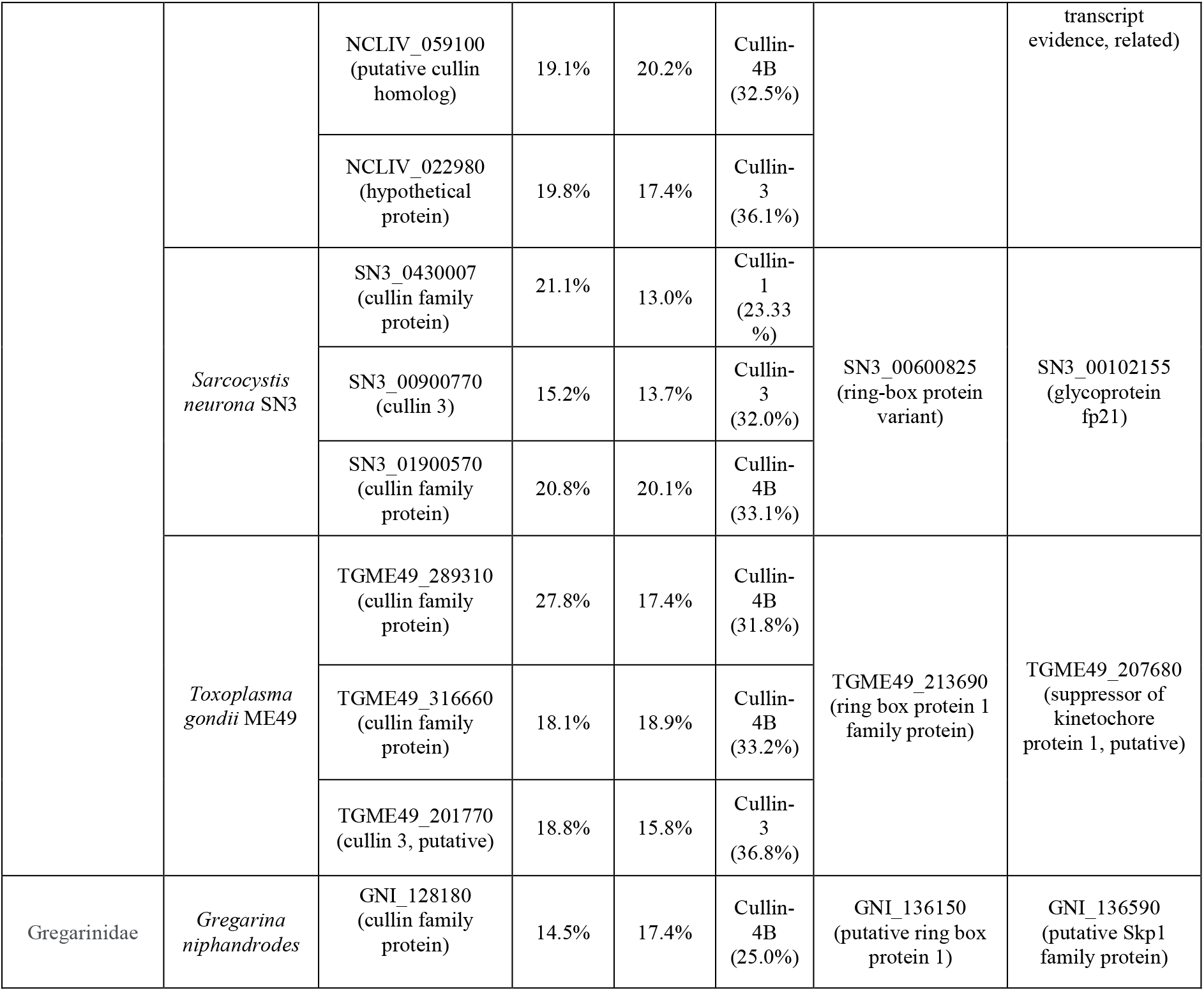
Putative CRL subunits in apicomplexan parasites. PfCullin-1, PfCullin-2, PfRbx1 and PfSkp1 homologs in reference Apicomplexan parasite strains of different families.

**Table S2.**
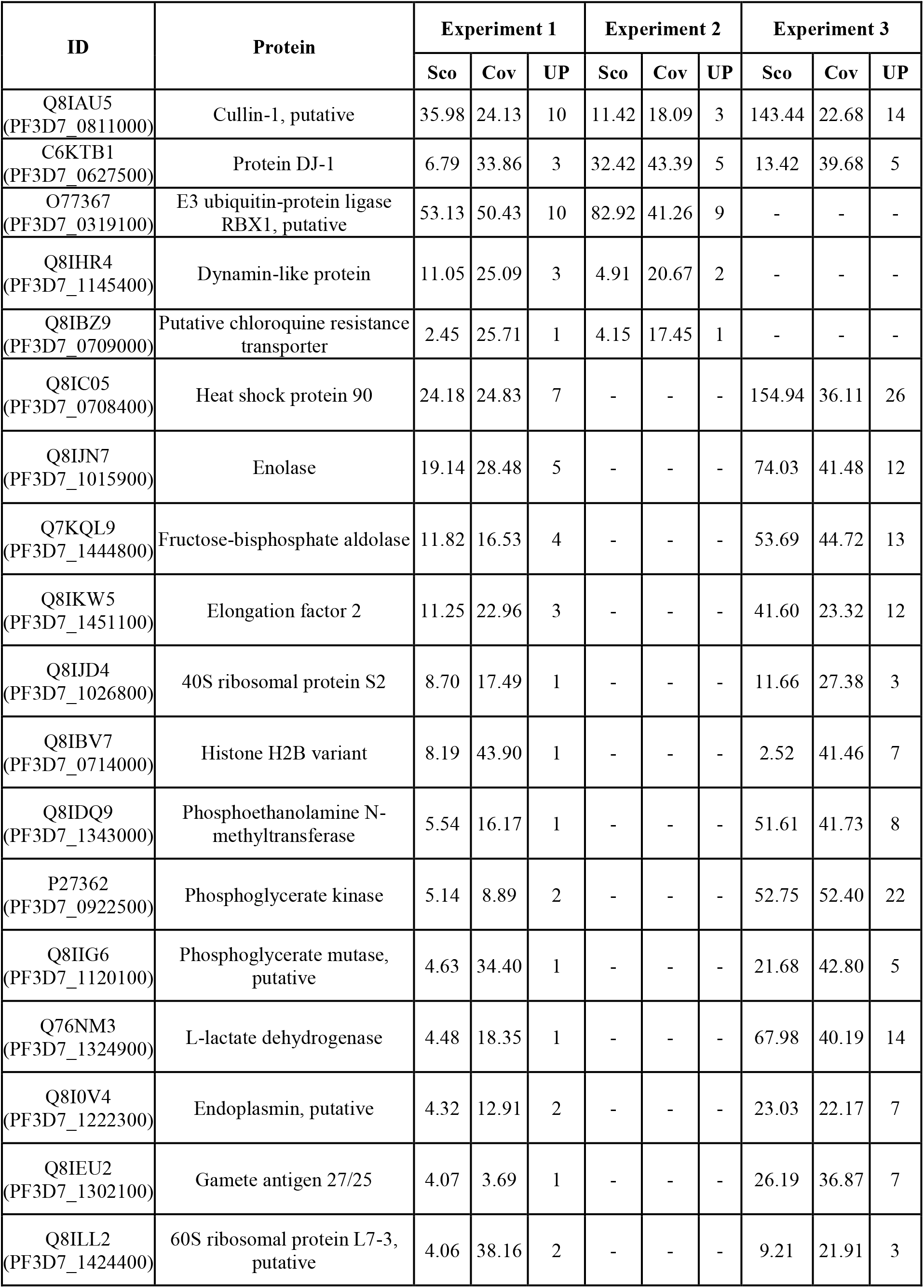
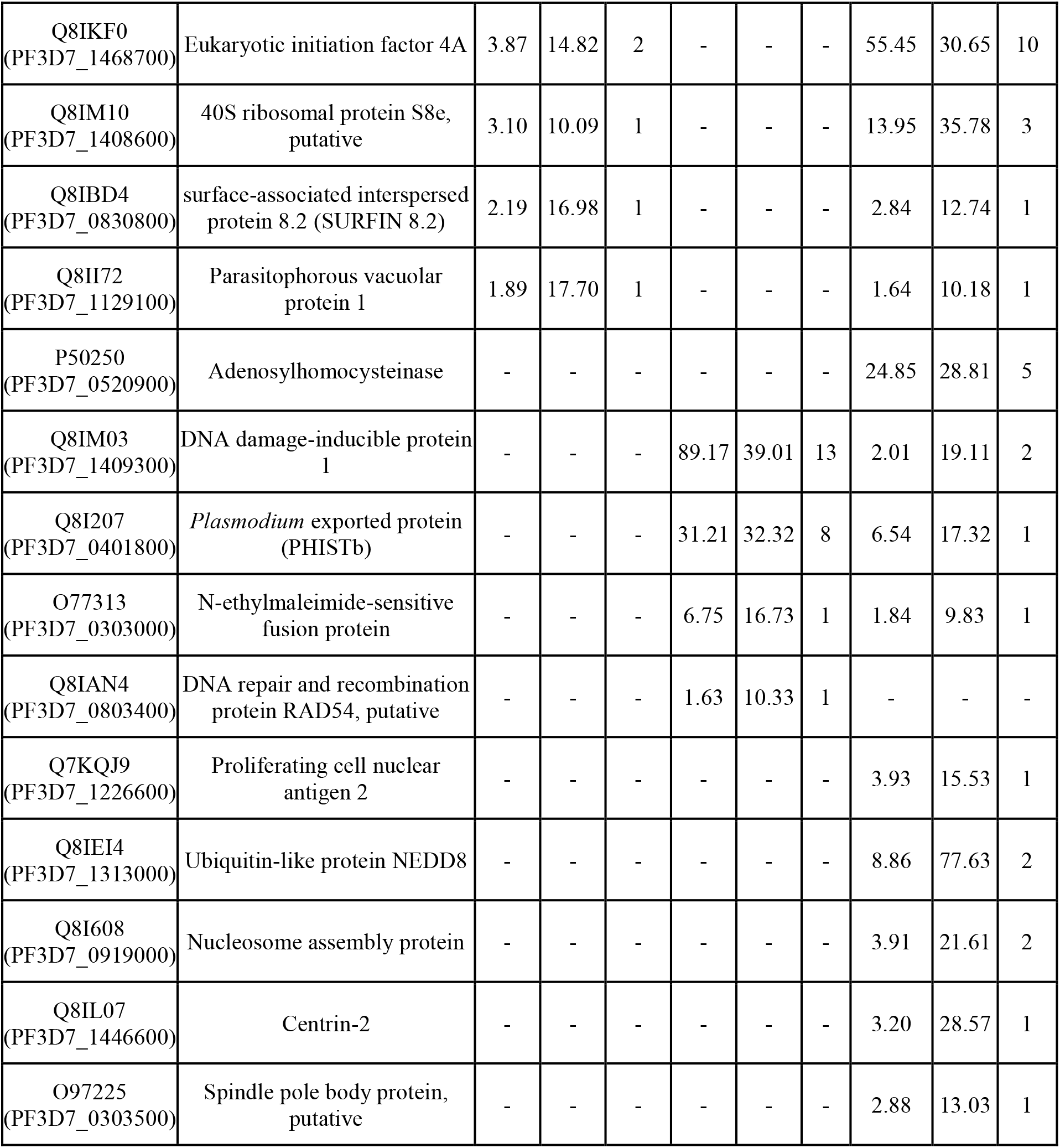
Proteins identified in the PfCullin1/GFP immunoprecipitate. Shown are the Uniprot (PlasmoDB) ID, score (Sc), coverage (Co) and unique peptides (UP) for each protein identified in three independent biological repeats.

**Table S3.**
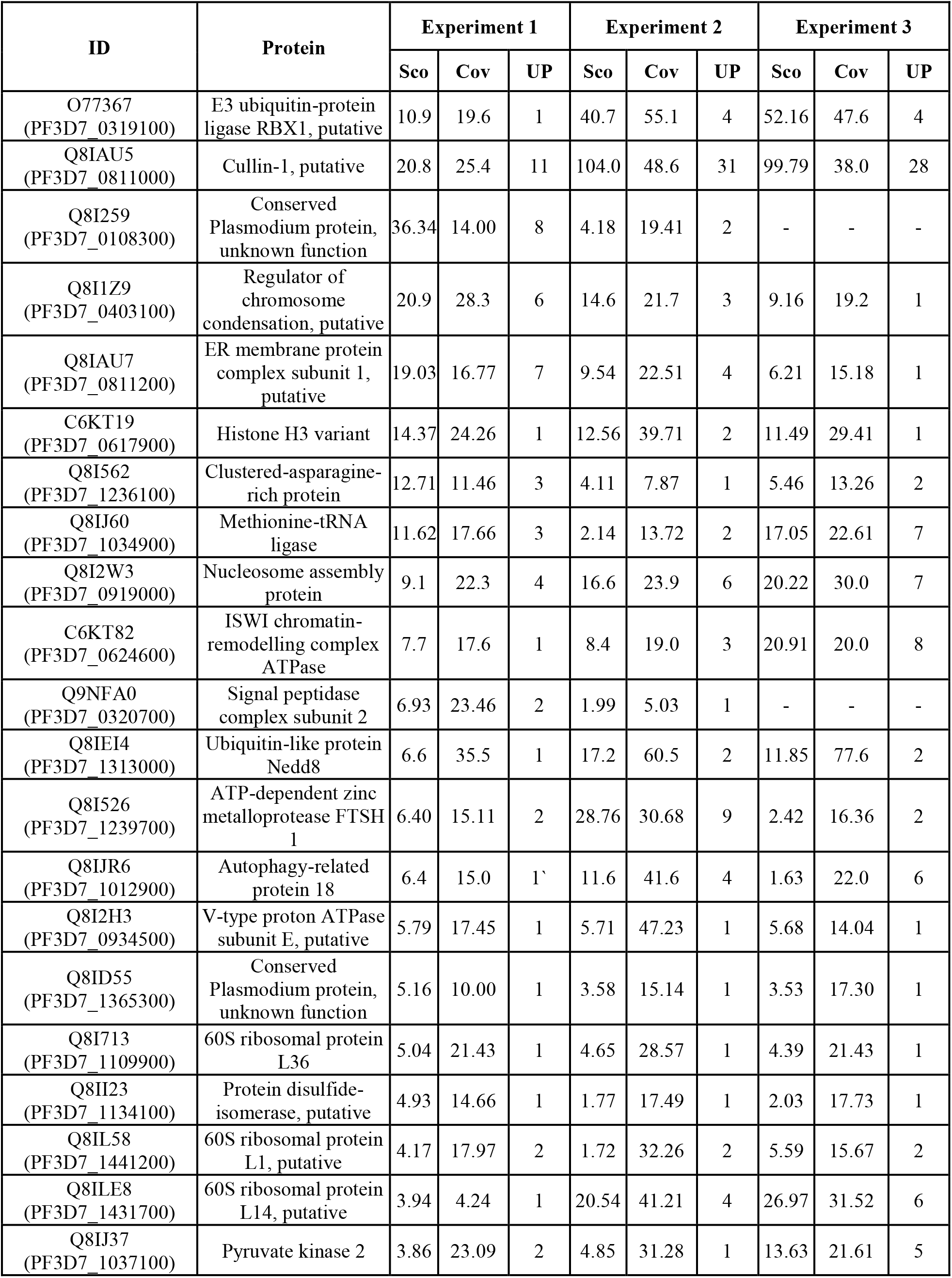
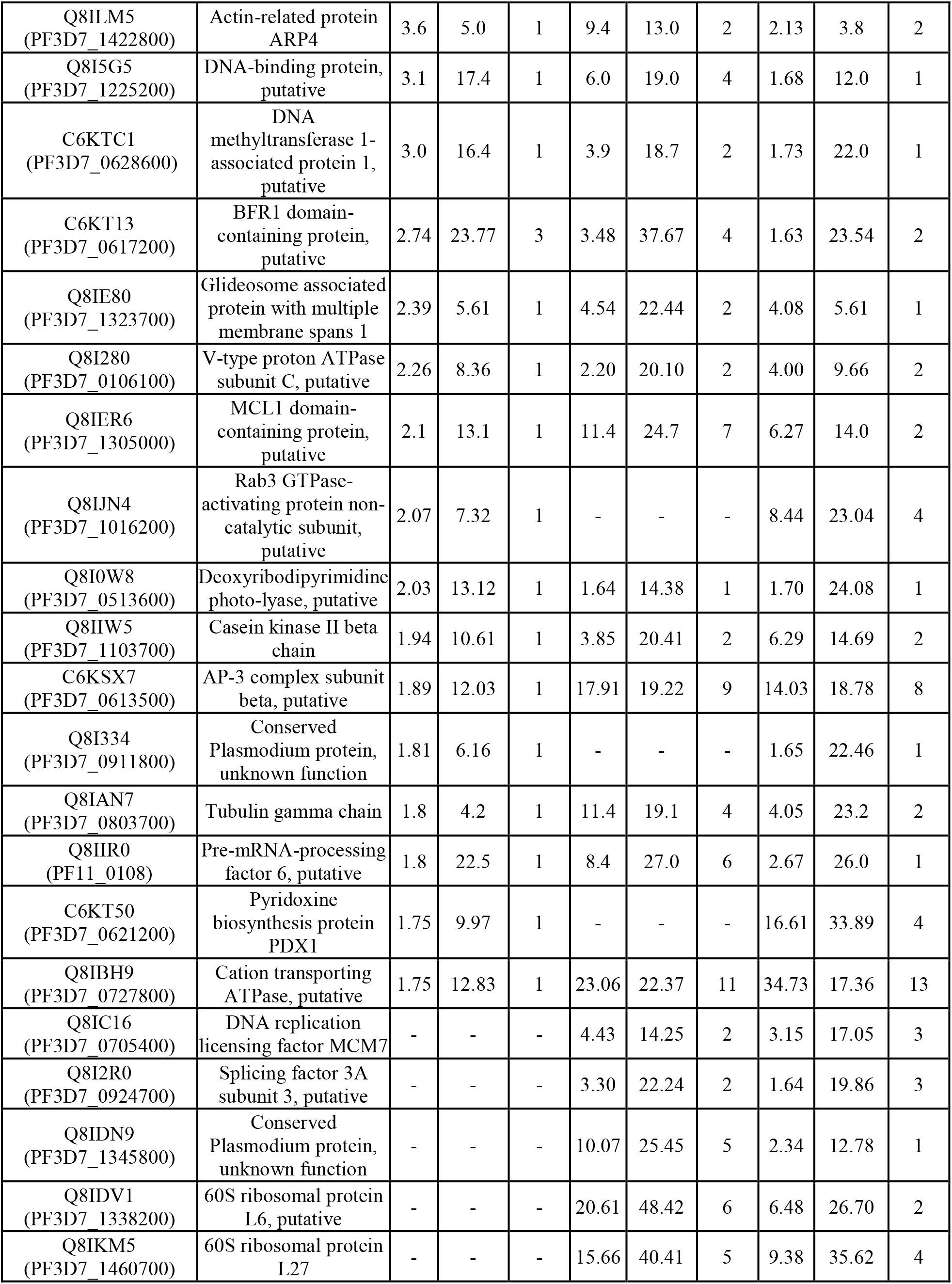
Proteins identified in the PfRbx1myc immunoprecipitate. Shown are the Uniprot (PlasmoDB) ID, score (Sc), coverage (Co) and unique peptides (UP) for proteins present in three independent biological repeats.

| ID | Protein | Experiment 1 |  |  | Experiment 2 |  |  | Experiment 3 |  |  |
| --- | --- | --- | --- | --- | --- | --- | --- | --- | --- | --- |
|  |  | Sc | Cov | UP | Sc | Cov | UP | Sc | Cov | UP |
| O77367<br>(PF3D7_0319100) | E3 ubiquitin-protein ligase RBX1, putative | 10.9 | 19.6 | 1 | 40.7 | 55.1 | 4 | 52.16 | 47.6 | 4 |
| Q8IAU5<br>(PF3D7_0811000) | Cullin-1, putative | 20.8 | 25.4 | 11 | 104.0 | 48.6 | 31 | 99.79 | 38.0 | 28 |
| Q8I259<br>(PF3D7_0108300) | Conserved Plasmodium protein, unknown function | 36.34 | 14.00 | 8 | 4.18 | 19.41 | 2 | - | - | - |
| Q8I1Z9<br>(PF3D7_0403100) | Regulator of chromosome condensation, putative | 20.9 | 28.3 | 6 | 14.6 | 21.7 | 3 | 9.16 | 19.2 | 1 |
| Q8IAU7<br>(PF3D7_0811200) | ER membrane protein complex subunit 1, putative | 19.03 | 16.77 | 7 | 9.54 | 22.51 | 4 | 6.21 | 15.18 | 1 |
| C6KT19<br>(PF3D7_0617900) | Histone H3 variant | 14.37 | 24.26 | 1 | 12.56 | 39.71 | 2 | 11.49 | 29.41 | 1 |
| Q8I562<br>(PF3D7_1236100) | Clustered-asparagine-rich protein | 12.71 | 11.46 | 3 | 4.11 | 7.87 | 1 | 5.46 | 13.26 | 2 |
| Q8IJ60<br>(PF3D7_1034900) | Methionine-tRNA ligase | 11.62 | 17.66 | 3 | 2.14 | 13.72 | 2 | 17.05 | 22.61 | 7 |
| Q8I2W3<br>(PF3D7_0919000) | Nucleosome assembly protein | 9.1 | 22.3 | 4 | 16.6 | 23.9 | 6 | 20.22 | 30.0 | 7 |
| C6KT82<br>(PF3D7_0624600) | ISWI chromatin-remodelling complex ATPase | 7.7 | 17.6 | 1 | 8.4 | 19.0 | 3 | 20.91 | 20.0 | 8 |
| Q9NFA0<br>(PF3D7_0320700) | Signal peptidase complex subunit 2 | 6.93 | 23.46 | 2 | 1.99 | 5.03 | 1 | - | - | - |
| Q8IEI4<br>(PF3D7_1313000) | Ubiquitin-like protein Nedd8 | 6.6 | 35.5 | 1 | 17.2 | 60.5 | 2 | 11.85 | 77.6 | 2 |
| Q8I526<br>(PF3D7_1239700) | ATP-dependent zinc metalloprotease FTSH 1 | 6.40 | 15.11 | 2 | 28.76 | 30.68 | 9 | 2.42 | 16.36 | 2 |
| Q8IJR6<br>(PF3D7_1012900) | Autophagy-related protein 18 | 6.4 | 15.0 | 1 | 11.6 | 41.6 | 4 | 1.63 | 22.0 | 6 |
| Q8I2H3<br>(PF3D7_0934500) | V-type proton ATPase subunit E, putative | 5.79 | 17.45 | 1 | 5.71 | 47.23 | 1 | 5.68 | 14.04 | 1 |
| Q8ID55<br>(PF3D7_1365300) | Conserved Plasmodium protein, unknown function | 5.16 | 10.00 | 1 | 3.58 | 15.14 | 1 | 3.53 | 17.30 | 1 |
| Q8I713<br>(PF3D7_1109900) | 60S ribosomal protein L36 | 5.04 | 21.43 | 1 | 4.65 | 28.57 | 1 | 4.39 | 21.43 | 1 |
| Q8II23<br>(PF3D7_1134100) | Protein disulfide-isomerase, putative | 4.93 | 14.66 | 1 | 1.77 | 17.49 | 1 | 2.03 | 17.73 | 1 |
| Q8IL58<br>(PF3D7_1441200) | 60S ribosomal protein L1, putative | 4.17 | 17.97 | 2 | 1.72 | 32.26 | 2 | 5.59 | 15.67 | 2 |
| Q8ILE8<br>(PF3D7_1431700) | 60S ribosomal protein L14, putative | 3.94 | 4.24 | 1 | 20.54 | 41.21 | 4 | 26.97 | 31.52 | 6 |
| Q8IJ37<br>(PF3D7_1037100) | Pyruvate kinase 2 | 3.86 | 23.09 | 2 | 4.85 | 31.28 | 1 | 13.63 | 21.61 | 5 |
| Q8ILM5<br>(PF3D7_1422800) | Actin-related protein<br>ARP4 | 3.6 | 5.0 | 1 | 9.4 | 13.0 | 2 | 2.13 | 3.8 | 2 |
| Q8I5G5<br>(PF3D7_1225200) | DNA-binding protein,<br>putative | 3.1 | 17.4 | 1 | 6.0 | 19.0 | 4 | 1.68 | 12.0 | 1 |
| C6KTC1<br>(PF3D7_0628600) | DNA<br>methyltransferase 1-associated protein 1,<br>putative | 3.0 | 16.4 | 1 | 3.9 | 18.7 | 2 | 1.73 | 22.0 | 1 |
| C6KT13<br>(PF3D7_0617200) | BFR1 domain-containing protein,<br>putative | 2.74 | 23.77 | 3 | 3.48 | 37.67 | 4 | 1.63 | 23.54 | 2 |
| Q8IE80<br>(PF3D7_1323700) | Glideosome associated<br>protein with multiple<br>membrane spans 1 | 2.39 | 5.61 | 1 | 4.54 | 22.44 | 2 | 4.08 | 5.61 | 1 |
| Q8I280<br>(PF3D7_0106100) | V-type proton ATPase<br>subunit C, putative | 2.26 | 8.36 | 1 | 2.20 | 20.10 | 2 | 4.00 | 9.66 | 2 |
| Q8IER6<br>(PF3D7_1305000) | MCL1 domain-containing protein,<br>putative | 2.1 | 13.1 | 1 | 11.4 | 24.7 | 7 | 6.27 | 14.0 | 2 |
| Q8IJN4<br>(PF3D7_1016200) | Rab3 GTPase-<br>activating protein non-<br>catalytic subunit,<br>putative | 2.07 | 7.32 | 1 | - | - | - | 8.44 | 23.04 | 4 |
| Q8I0W8<br>(PF3D7_0513600) | Deoxyribodipyrimidine<br>photo-lyase, putative | 2.03 | 13.12 | 1 | 1.64 | 14.38 | 1 | 1.70 | 24.08 | 1 |
| Q8IIW5<br>(PF3D7_1103700) | Casein kinase II beta<br>chain | 1.94 | 10.61 | 1 | 3.85 | 20.41 | 2 | 6.29 | 14.69 | 2 |
| C6KSX7<br>(PF3D7_0613500) | AP-3 complex subunit<br>beta, putative | 1.89 | 12.03 | 1 | 17.91 | 19.22 | 9 | 14.03 | 18.78 | 8 |
| Q8I334<br>(PF3D7_0911800) | Conserved<br>Plasmodium protein,<br>unknown function | 1.81 | 6.16 | 1 | - | - | - | 1.65 | 22.46 | 1 |
| Q8IAN7<br>(PF3D7_0803700) | Tubulin gamma chain | 1.8 | 4.2 | 1 | 11.4 | 19.1 | 4 | 4.05 | 23.2 | 2 |
| Q8IIR0<br>(PF11_0108) | Pre-mRNA-processing<br>factor 6, putative | 1.8 | 22.5 | 1 | 8.4 | 27.0 | 6 | 2.67 | 26.0 | 1 |
| C6KT50<br>(PF3D7_0621200) | Pyridoxine<br>biosynthesis protein<br>PDX1 | 1.75 | 9.97 | 1 | - | - | - | 16.61 | 33.89 | 4 |
| Q8IBH9<br>(PF3D7_0727800) | Cation transporting<br>ATPase, putative | 1.75 | 12.83 | 1 | 23.06 | 22.37 | 11 | 34.73 | 17.36 | 13 |
| Q8IC16<br>(PF3D7_0705400) | DNA replication<br>licensing factor MCM7 | - | - | - | 4.43 | 14.25 | 2 | 3.15 | 17.05 | 3 |
| Q8I2R0<br>(PF3D7_0924700) | Splicing factor 3A<br>subunit 3, putative | - | - | - | 3.30 | 22.24 | 2 | 1.64 | 19.86 | 3 |
| Q8IDN9<br>(PF3D7_1345800) | Conserved<br>Plasmodium protein,<br>unknown function | - | - | - | 10.07 | 25.45 | 5 | 2.34 | 12.78 | 1 |
| Q8IDV1<br>(PF3D7_1338200) | 60S ribosomal protein<br>L6, putative | - | - | - | 20.61 | 48.42 | 6 | 6.48 | 26.70 | 2 |
| Q8IKM5<br>(PF3D7_1460700) | 60S ribosomal protein<br>L27 | - | - | - | 15.66 | 40.41 | 5 | 9.38 | 35.62 | 4 |

**Table S4.** Proteins identified in the PfSkp1GFP immunoprecipitate. Shown are the Uniprot (PlasmoDB) ID, score (Sc), coverage (Co) and unique peptides (UP) for each protein present in three independent biological repeats.

| Accession ID | Protein | Experiment 1 |  |  | Experiment 2 |  |  | Experiment 3 |  |  |
| --- | --- | --- | --- | --- | --- | --- | --- | --- | --- | --- |
|  |  | Sco | Cov | UP | Sco | Cov | UP | Sco | Cov | UP |
| Q8ID38<br>(PF3D7_1367000) | Suppressor of kinetochore protein 1, putative (Skp1) | 138.3 | 24.1 | 4 | 175.7 | 43.8 | 5 | 158.7 | 22.2 | 4 |
| Q8I542<br>(PF3D7_1238100) | Calcyclin binding protein, putative | 1.6 | 18.0 | 1 | 2.0 | 12.7 | 1 | 18.0 | 3.5 | 1 |
| Q8I207<br>(PF3D7_0401800) | Plasmodium exported protein PHISTb | - | - | - | 8.1 | 21.4 | 2 | 78.5 | 39.3 | 17 |
| C6KTB1<br>(PF3D7_0627500) | Protein DJ-1 | 8.8 | 41.3 | 2 | 18.9 | 47.1 | 6 | 4.1 | 37.0 | 2 |
| Q8IHR4<br>(PF3D7_1145400) | Dynamin-like protein | 9.2 | 30.1 | 3 | 5.5 | 17.3 | 2 | 5.4 | 23.4 | 3 |
| Q8IAU5<br>(PF3D7_0811000) | Cullin-1, putative | 3.9 | 13.5 | 1 | 31.5 | 24.7 | 16 | - | - | - |
| Q7KQK2<br>(PF3D7_1211800) | PfpUB polyubiquitin | 15.8 | 76.6 | 3 | 10.7 | 35.4 | 3 | - | - | - |
| C6KT37<br>(PF3D7_0619700) | F-box protein FBXO1, putative | 2.2 | 27.3 | 2 | 7.8 | 31.1 | 4 | - | - | - |
| Q8I2U5<br>(PF3D7_0920800) | Inosine-5'-monophosphate dehydrogenase | 25.23 | 27.25 | 7 | 2.08 | 14.12 | 1 | - | - | - |
| Q8IIF0<br>(PF3D7_1121600) | Exported protein 1 | 9.96 | 17.28 | 2 | 6.11 | 6.79 | 2 | - | - | - |
| Q8IK02<br>(PF3D7_1003500) | 40S ribosomal protein S20e, putative | 3.71 | 51.69 | 1 | 4.79 | 20.34 | 1 | - | - | - |
| Q8ILV2<br>(PF3D7_1414300) | 60S ribosomal protein L10, putative | 1.78 | 15.07 | 1 | 2.60 | 9.13 | 1 | - | - | - |
| C6KST5<br>(PF3D7_0608700) | T-complex protein 1 subunit zeta | 1.61 | 18.78 | 1 | 7.72 | 25.60 | 3 | - | - | - |
| O77372<br>(PF3D7_0318500) | conserved Plasmodium protein | 18.45 | 30.21 | 1 | - | - | - | 8.26 | 18.25 | 1 |
| Q9U0N1<br>(PF3D7_0113000) | Glutamic acid-rich protein GARP | 9.34 | 17.38 | 2 | - | - | - | 21.73 | 20.95 | 4 |
| Q8ILQ9<br>(PF3D7_1419100) | ATP-dependent RNA helicase DDX55 | 3.35 | 31.03 | 1 | - | - | - | 3.66 | 28.63 | 1 |
| Q8IM03<br>(PF3D7_1409300) | DNA damage-inducible protein 1 | - | - | - | 77.2 | 38.7 | 13 | 90.8 | 40.1 | 14 |
| Q8IAN4<br>(PF3D7_0803400) | DNA repair and recombination protein RAD54,<br>putative | - | - | - | 3.08 | 13.48 | 1 | 1.91 | 13.88 | 1 |

**Table S5.**
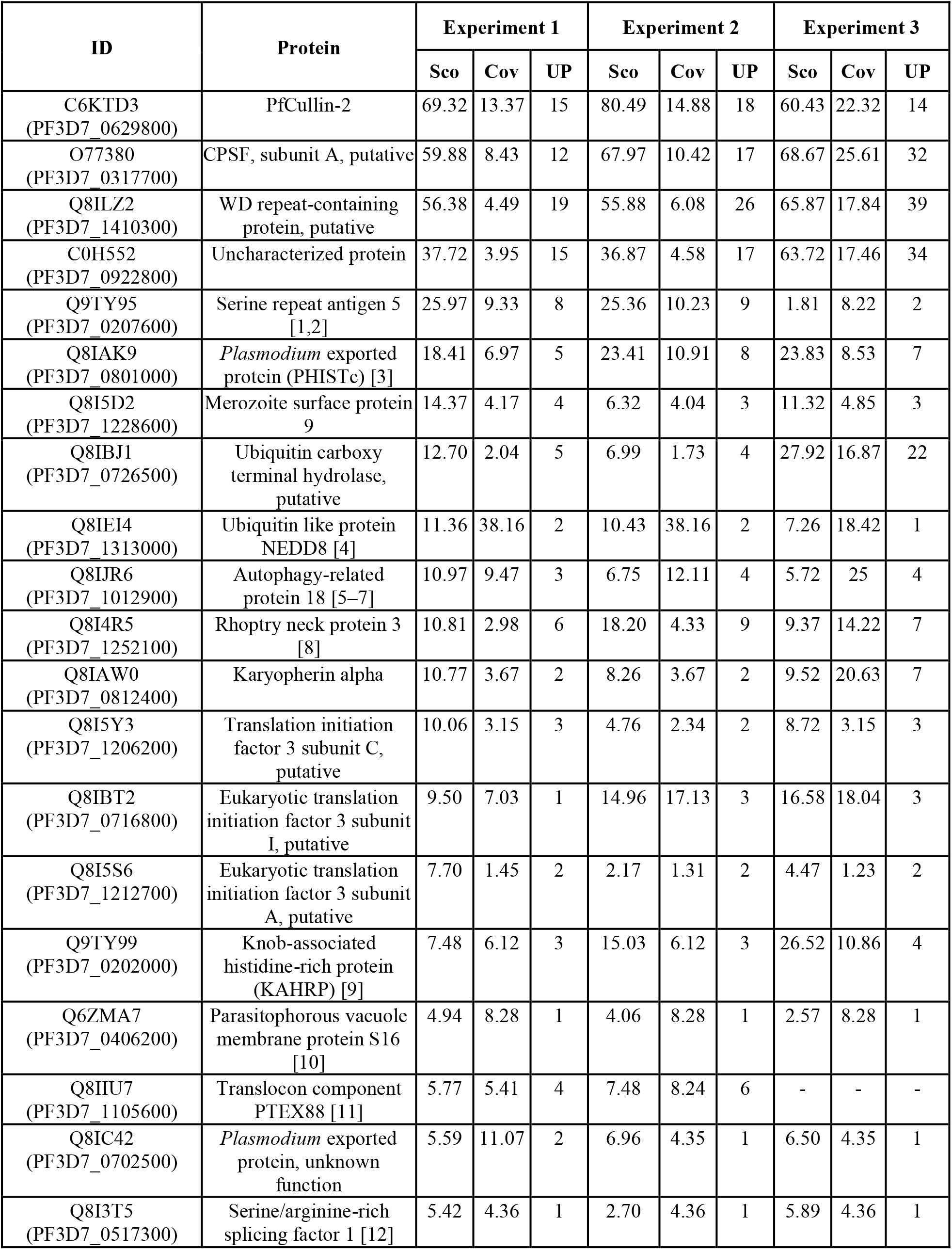
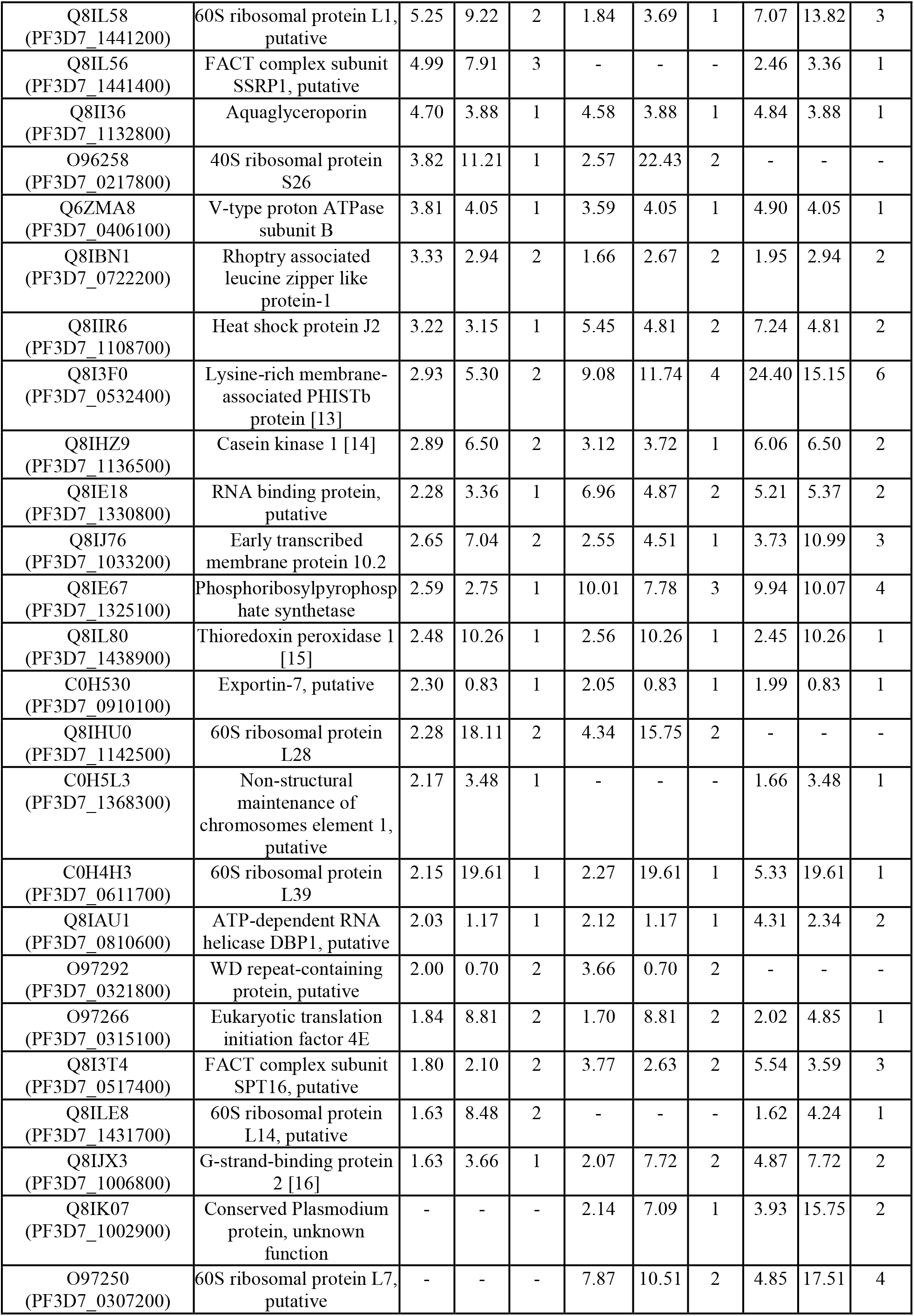
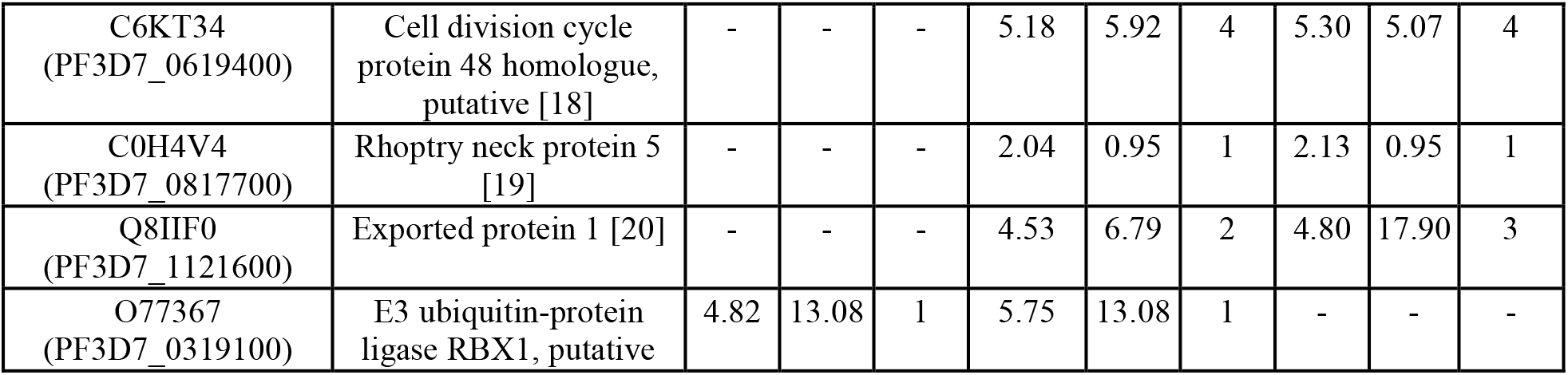
Proteins identified in the PfCullin-2/cDD_HA_ immunoprecipitate. Shown are the Uniprot (PlasmoDB) ID, score (Sco), coverage (Cov) and unique peptides (UP) for proteins present in three independent biological repeats. The characterized *Plasmodium* proteins are referenced.

**References**

1. Collins CR, Hackett F, Atid J, Tan MSY, Blackman MJ. The *Plasmodium falciparum* pseudoprotease SERA5 regulates the kinetics and efficiency of malaria parasite egress from host erythrocytes. PLoS Pathog. 2017;13: e1006453-. Available: https://doi.org/10.1371/journal.ppat.1006453

2. Stallmach R, Kavishwar M, Withers-Martinez C, Hackett F, Collins CR, Howell SA, et al. Plasmodium falciparum SERA5 plays a non-enzymatic role in the malarial asexual blood-stage lifecycle. Mol Microbiol. 2015;96: 368–387. doi:https://doi.org/10.1111/mmi.12941

3. Nagaoka H, Kanoi BN, Morita M, Nakata T, Palacpac NMQ, Egwang TG, et al. Characterization of a Plasmodium falciparum PHISTc protein, PF3D7_0801000, in blood-stage malaria parasites. Parasitol Int. 2021;80: 102240. doi:https://doi.org/10.1016/j.parint.2020.102240

4. Bhattacharjee M, AN, SR et al. Characterization of Plasmodium falciparum NEDD8 and identification of cullins as its substrates.

5. Priyanka B, Anuj T, Vandana T, Asif M, Pushkar S. Autophagy-Related Protein ATG18 Regulates Apicoplast Biogenesis in Apicomplexan Parasites. mBio. 2017;8: e01468-17. doi:10.1128/mBio.01468-17

6. Renu Sudhakar DDSTSGPSS. Plasmodium falciparum Atg18 localizes to the food vacuole via interaction with the multi-drug resistance protein 1 and phosphatidylinositol 3-phosphate.

7. Breglio KF, AR, ER et al. A single nucleotide polymorphism in the Plasmodium falciparum atg18 gene associates with artemisinin resistance and confers enhanced parasite survival under nutrient deprivation.

8. M LL, Yvonne A, S SE, Matthias G, Joshua Z, Takafumi T, et al. Deletion of Plasmodium falciparum Protein RON3 Affects the Functional Translocation of Exported Proteins and Glucose Uptake. mBio. 2019;10: e01460-19. doi:10.1128/mBio.01460-19

9. Breglio KF, AR, ER et al. A single nucleotide polymorphism in the Plasmodium falciparum atg18 gene associates with artemisinin resistance and confers enhanced parasite survival under nutrient deprivation.

10. Darin Kongkasuriyachai et al. 2004. Functional analysis of Plasmodium falciparum parasitophorous vacuole membrane protein (Pfs16) during gametocytogenesis and gametogenesis by targeted gene disruption.

11. Chisholm SA MELRDMGS et al. (2016). Contrasting Inducible Knockdown of the Auxiliary PTEX Component PTEX88 in P. falciparum and P. berghei Unmasks a Role in Parasite Virulence.

12. Eshar S, AL, RA, RP, FY, M-GY, KR, LM and DR (2015),. Regulatory elements of the PfSR1 protein from P. falciparum.

13. Proellocks NI HSBD et al. A lysine-rich membrane-associated PHISTb protein involved in alteration of the cytoadhesive properties of Plasmodium falciparum-infected red blood cells.

14. Batty MB SRD-SDDCG-BJF. Interaction of Plasmodium falciparum casein kinase 1 with components of host cell protein trafficking machinery.

15. Komaki-Yasuda K, OM, NK, KI, & KS (2013). Identification of a Novel and Unique Transcription Factor in the Intraerythrocytic Stage of Plasmodium falciparum.

16. Niikura M FTFKISAHKF. G-strand binding protein 2 is involved in asexual and sexual development of Plasmodium berghei.

17. Spork S, HJA, MK, SM, KWA, CT, SG, MUG, & PJM (2009). An Unusual ERAD-Like Complex Is Targeted to the Apicoplast of Plasmodium falciparum.

18. Curtidor H PLA-PGVMPMPMA. Plasmodium falciparum rhoptry neck protein 5 peptides bind to human red blood cells and inhibit parasite invasion.

19. Lisewski AM, Quiros JP, Mittal M, Putluri N, Sreekumar A, Haeggström JZ, et al. Potential role of Plasmodium falciparum exported protein 1 in the chloroquine mode of action. Int J Parasitol Drugs Drug Resist. 2018;8: 31–35. doi:https://doi.org/10.1016/j.ijpddr.2017.12.003

**Table S6.**
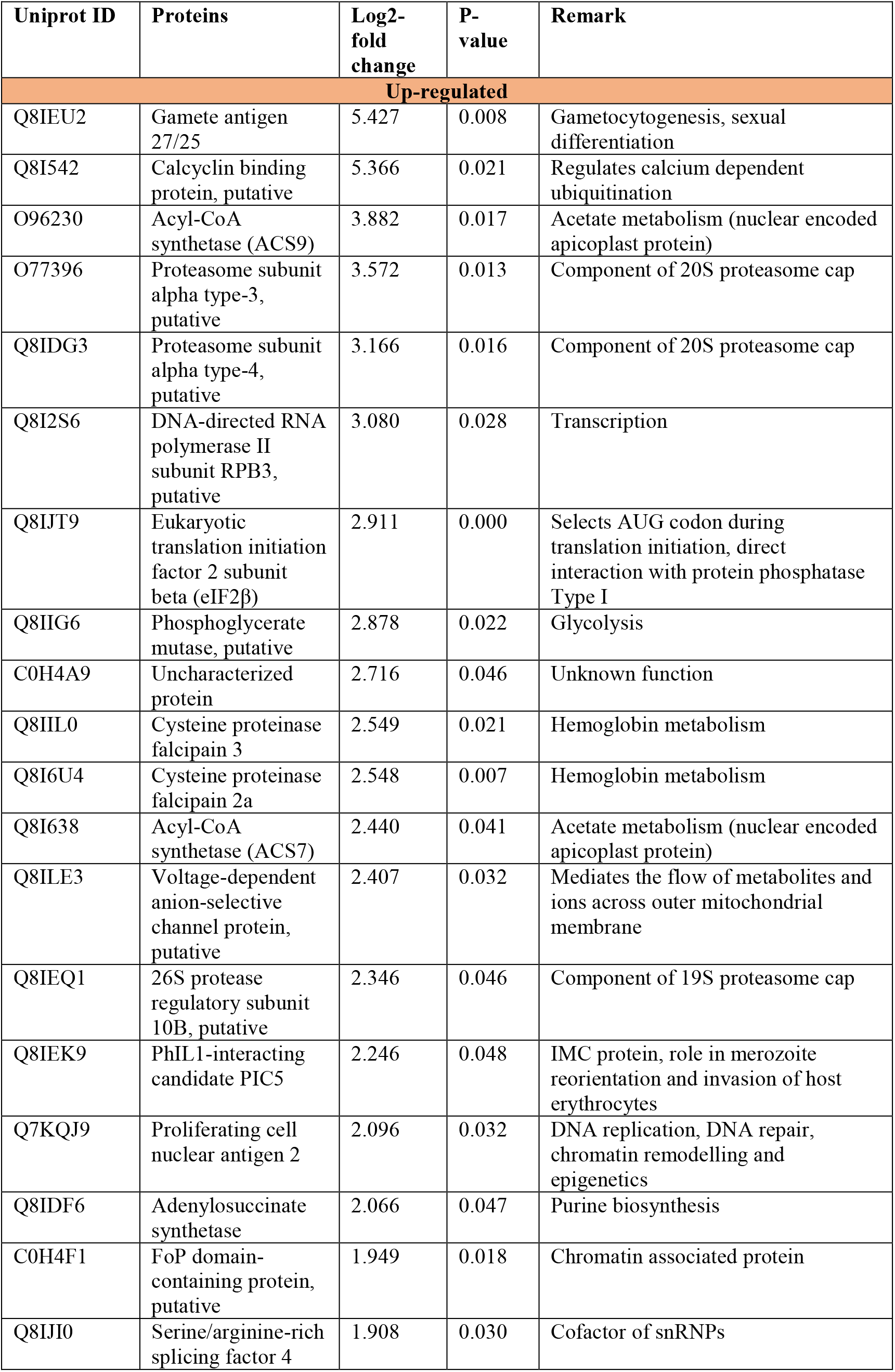
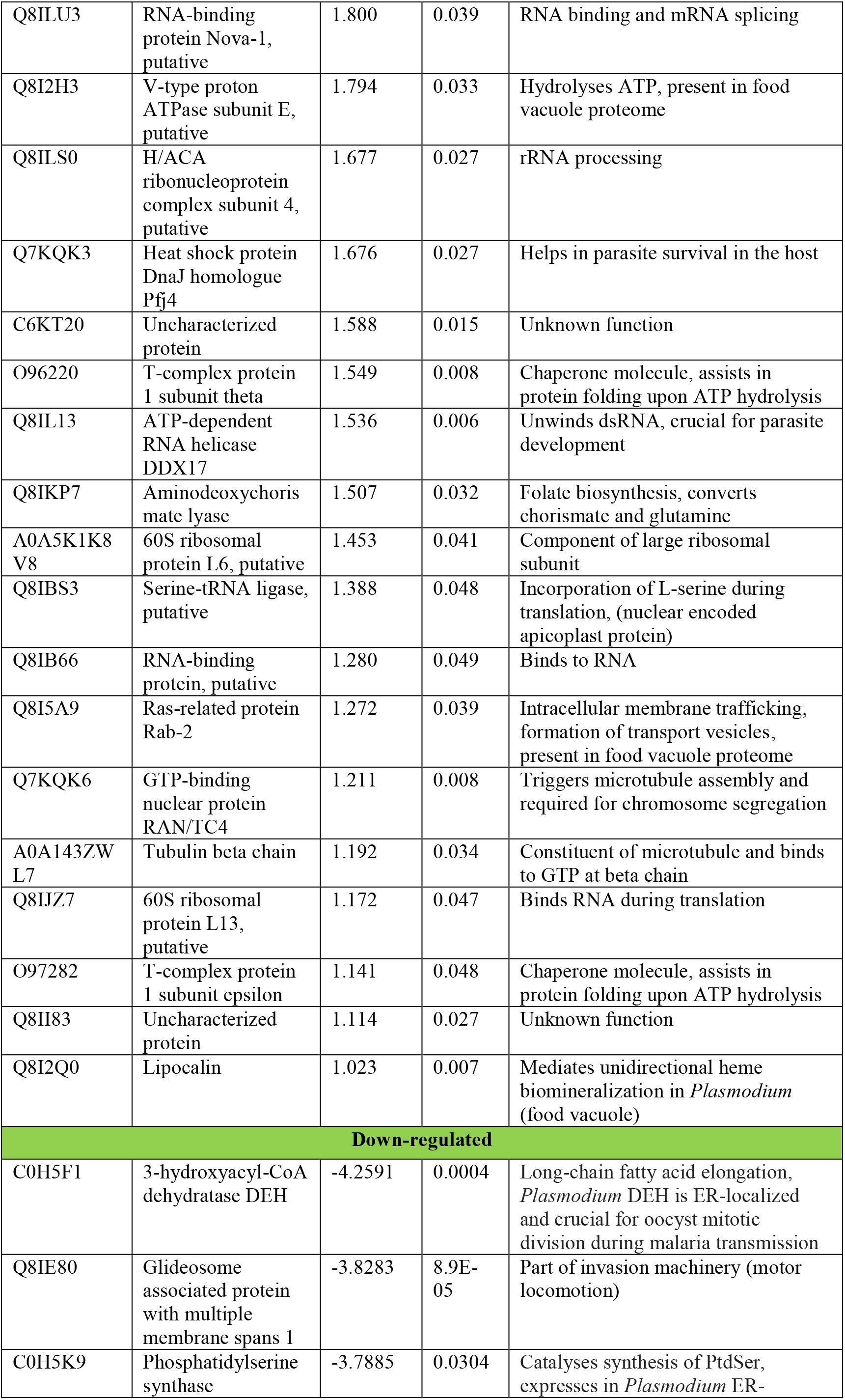
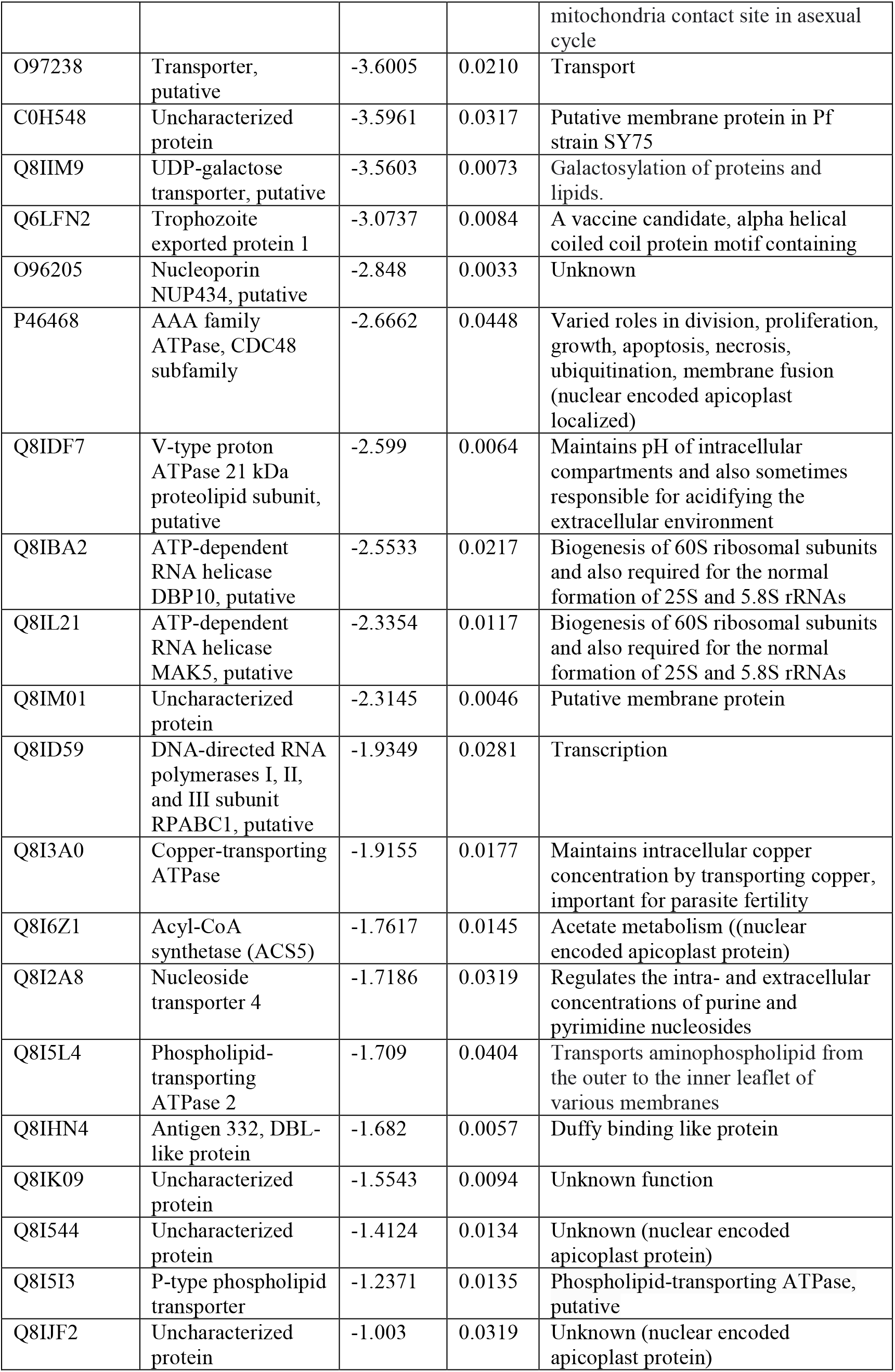
High confidence up- and down-regulated proteins in PfCullin-2-depleted trophozoites upon. Proteins shown/predicted to be targeted to the apicoplast and mitochondria are highlighted.

| Uniprot ID | Proteins | Log2-fold change | P-value | Remark |
| --- | --- | --- | --- | --- |
| **Up-regulated** |  |  |  |  |
| Q8IEU2 | Gamete antigen 27/25 | 5.427 | 0.008 | Gametocytogenesis, sexual differentiation |
| Q8I542 | Calcyclin binding protein, putative | 5.366 | 0.021 | Regulates calcium dependent ubiquitination |
| O96230 | Acyl-CoA synthetase (ACS9) | 3.882 | 0.017 | Acetate metabolism (nuclear encoded apicoplast protein) |
| O77396 | Proteasome subunit alpha type-3, putative | 3.572 | 0.013 | Component of 20S proteasome cap |
| Q8IDG3 | Proteasome subunit alpha type-4, putative | 3.166 | 0.016 | Component of 20S proteasome cap |
| Q8I2S6 | DNA-directed RNA polymerase II subunit RPB3, putative | 3.080 | 0.028 | Transcription |
| Q8IJT9 | Eukaryotic translation initiation factor 2 subunit beta (eIF2β) | 2.911 | 0.000 | Selects AUG codon during translation initiation, direct interaction with protein phosphatase Type I |
| Q8IIG6 | Phosphoglycerate mutase, putative | 2.878 | 0.022 | Glycolysis |
| C0H4A9 | Uncharacterized protein | 2.716 | 0.046 | Unknown function |
| Q8IIL0 | Cysteine proteinase falcipain 3 | 2.549 | 0.021 | Hemoglobin metabolism |
| Q8I6U4 | Cysteine proteinase falcipain 2a | 2.548 | 0.007 | Hemoglobin metabolism |
| Q8I638 | Acyl-CoA synthetase (ACS7) | 2.440 | 0.041 | Acetate metabolism (nuclear encoded apicoplast protein) |
| Q8ILE3 | Voltage-dependent anion-selective channel protein, putative | 2.407 | 0.032 | Mediates the flow of metabolites and ions across outer mitochondrial membrane |
| Q8IEQ1 | 26S protease regulatory subunit 10B, putative | 2.346 | 0.046 | Component of 19S proteasome cap |
| Q8IEK9 | PhIL1-interacting candidate PIC5 | 2.246 | 0.048 | IMC protein, role in merozoite reorientation and invasion of host erythrocytes |
| Q7KQJ9 | Proliferating cell nuclear antigen 2 | 2.096 | 0.032 | DNA replication, DNA repair, chromatin remodelling and epigenetics |
| Q8IDF6 | Adenylosuccinate synthetase | 2.066 | 0.047 | Purine biosynthesis |
| C0H4F1 | FoP domain-containing protein, putative | 1.949 | 0.018 | Chromatin associated protein |
| Q8IJI0 | Serine/arginine-rich splicing factor 4 | 1.908 | 0.030 | Cofactor of snRNPs |
| Q8ILU3 | RNA-binding protein Nova-1, putative | 1.800 | 0.039 | RNA binding and mRNA splicing |
| Q8I2H3 | V-type proton ATPase subunit E, putative | 1.794 | 0.033 | Hydrolyses ATP, present in food vacuole proteome |
| Q8ILS0 | H/ACA ribonucleoprotein complex subunit 4, putative | 1.677 | 0.027 | rRNA processing |
| Q7KQK3 | Heat shock protein DnaJ homologue Pfj4 | 1.676 | 0.027 | Helps in parasite survival in the host |
| C6KT20 | Uncharacterized protein | 1.588 | 0.015 | Unknown function |
| O96220 | T-complex protein 1 subunit theta | 1.549 | 0.008 | Chaperone molecule, assists in protein folding upon ATP hydrolysis |
| Q8IL13 | ATP-dependent RNA helicase DDX17 | 1.536 | 0.006 | Unwinds dsRNA, crucial for parasite development |
| Q8IKP7 | Aminodeoxychorismate lyase | 1.507 | 0.032 | Folate biosynthesis, converts chorismate and glutamine |
| A0A5K1K8 V8 | 60S ribosomal protein L6, putative | 1.453 | 0.041 | Component of large ribosomal subunit |
| Q8IBS3 | Serine-tRNA ligase, putative | 1.388 | 0.048 | Incorporation of L-serine during translation, (nuclear encoded apicoplast protein) |
| Q8IB66 | RNA-binding protein, putative | 1.280 | 0.049 | Binds to RNA |
| Q8I5A9 | Ras-related protein Rab-2 | 1.272 | 0.039 | Intracellular membrane trafficking, formation of transport vesicles, present in food vacuole proteome |
| Q7KQK6 | GTP-binding nuclear protein RAN/TC4 | 1.211 | 0.008 | Triggers microtubule assembly and required for chromosome segregation |
| A0A143ZW L7 | Tubulin beta chain | 1.192 | 0.034 | Constituent of microtubule and binds to GTP at beta chain |
| Q8IJZ7 | 60S ribosomal protein L13, putative | 1.172 | 0.047 | Binds RNA during translation |
| O97282 | T-complex protein 1 subunit epsilon | 1.141 | 0.048 | Chaperone molecule, assists in protein folding upon ATP hydrolysis |
| Q8II83 | Uncharacterized protein | 1.114 | 0.027 | Unknown function |
| Q8I2Q0 | Lipocalin | 1.023 | 0.007 | Mediates unidirectional heme biomineralization in *Plasmodium* (food vacuole) |
| **Down-regulated** |  |  |  |  |
| C0H5F1 | 3-hydroxyacyl-CoA dehydratase DEH | -4.2591 | 0.0004 | Long-chain fatty acid elongation, *Plasmodium* DEH is ER-localized and crucial for oocyst mitotic division during malaria transmission |
| Q8IE80 | Glideosome associated protein with multiple membrane spans 1 | -3.8283 | 8.9E-05 | Part of invasion machinery (motor locomotion) |
| C0H5K9 | Phosphatidylserine synthase | -3.7885 | 0.0304 | Catalyses synthesis of PtdSer, expresses in *Plasmodium* ER- mitochondria contact site in asexual cycle |
| O97238 | Transporter, putative | -3.6005 | 0.0210 | Transport |
| C0H548 | Uncharacterized protein | -3.5961 | 0.0317 | Putative membrane protein in Pf strain SY75 |
| Q8IIM9 | UDP-galactose transporter, putative | -3.5603 | 0.0073 | Galactosylation of proteins and lipids. |
| Q6LFN2 | Trophozoite exported protein 1 | -3.0737 | 0.0084 | A vaccine candidate, alpha helical coiled coil protein motif containing |
| O96205 | Nucleoporin NUP434, putative | -2.848 | 0.0033 | Unknown |
| P46468 | AAA family ATPase, CDC48 subfamily | -2.6662 | 0.0448 | Varied roles in division, proliferation, growth, apoptosis, necrosis, ubiquitination, membrane fusion (nuclear encoded apicoplast localized) |
| Q8IDF7 | V-type proton ATPase 21 kDa proteolipid subunit, putative | -2.599 | 0.0064 | Maintains pH of intracellular compartments and also sometimes responsible for acidifying the extracellular environment |
| Q8IBA2 | ATP-dependent RNA helicase DBP10, putative | -2.5533 | 0.0217 | Biogenesis of 60S ribosomal subunits and also required for the normal formation of 25S and 5.8S rRNAs |
| Q8IL21 | ATP-dependent RNA helicase MAK5, putative | -2.3354 | 0.0117 | Biogenesis of 60S ribosomal subunits and also required for the normal formation of 25S and 5.8S rRNAs |
| Q8IM01 | Uncharacterized protein | -2.3145 | 0.0046 | Putative membrane protein |
| Q8ID59 | DNA-directed RNA polymerases I, II, and III subunit RPABC1, putative | -1.9349 | 0.0281 | Transcription |
| Q8I3A0 | Copper-transporting ATPase | -1.9155 | 0.0177 | Maintains intracellular copper concentration by transporting copper, important for parasite fertility |
| Q8I6Z1 | Acyl-CoA synthetase (ACS5) | -1.7617 | 0.0145 | Acetate metabolism ((nuclear encoded apicoplast protein) |
| Q8I2A8 | Nucleoside transporter 4 | -1.7186 | 0.0319 | Regulates the intra- and extracellular concentrations of purine and pyrimidine nucleosides |
| Q8I5L4 | Phospholipid-transporting ATPase 2 | -1.709 | 0.0404 | Transports aminophospholipid from the outer to the inner leaflet of various membranes |
| Q8IHN4 | Antigen 332, DBL-like protein | -1.682 | 0.0057 | Duffy binding like protein |
| Q8IK09 | Uncharacterized protein | -1.5543 | 0.0094 | Unknown function |
| Q8I544 | Uncharacterized protein | -1.4124 | 0.0134 | Unknown (nuclear encoded apicoplast protein) |
| Q8I5I3 | P-type phospholipid transporter | -1.2371 | 0.0135 | Phospholipid-transporting ATPase, putative |
| Q8IJF2 | Uncharacterized protein | -1.003 | 0.0319 | Unknown (nuclear encoded apicoplast protein) |

**Table S7.**
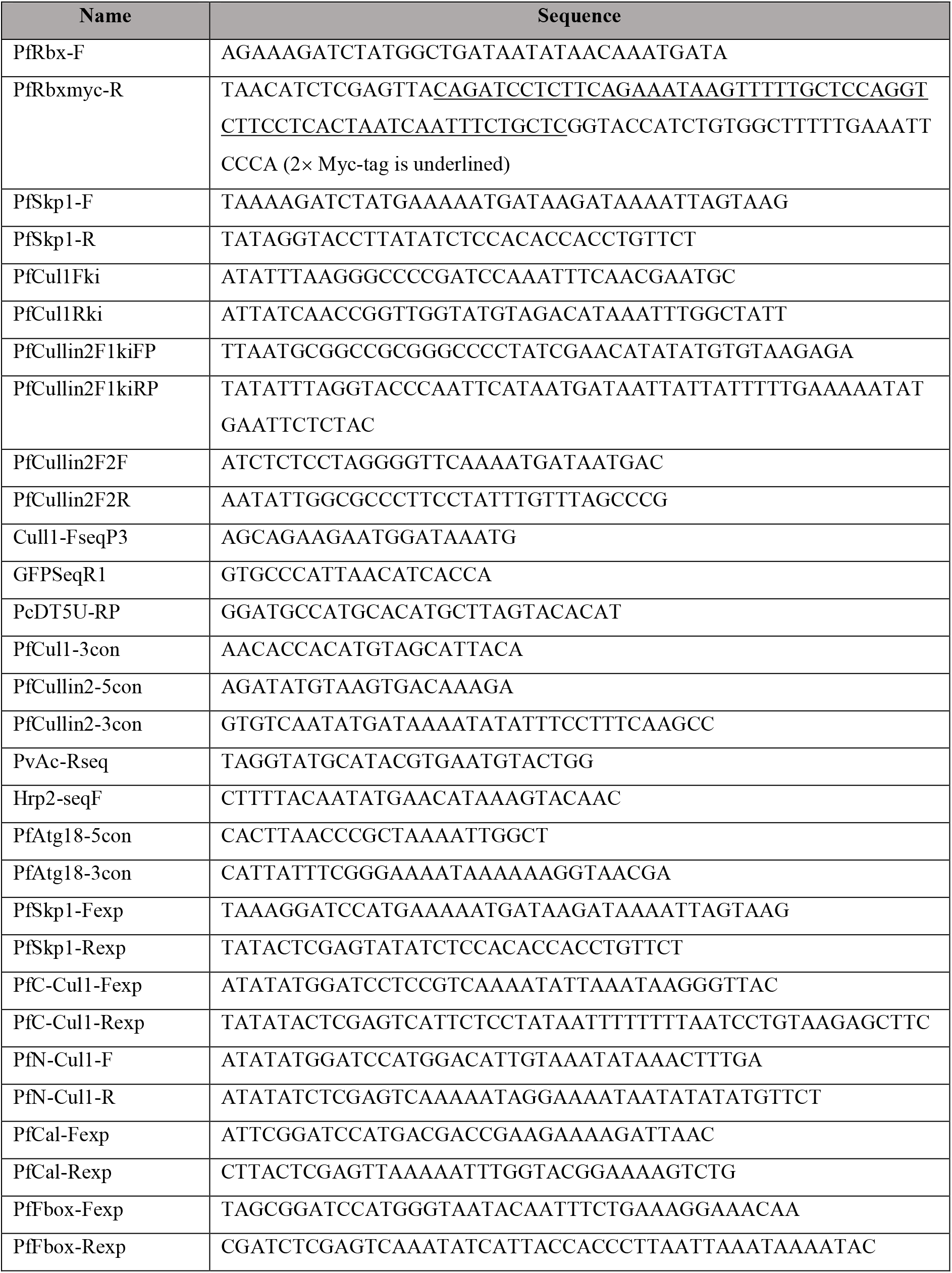
Primer used in the study.

## Notes

### Competing Interest Statement

The authors have declared no competing interest.

